# RNA m6A mediates *de novo* DNA methylation and heritable epigenetic silencing in human cells

**DOI:** 10.64898/2026.10.21.760774

**Authors:** Nikita S. Divekar, Ngan T. Phan, Luis G. Palma, Justin P. Lubin, Anna E. Christenson, Scarleth C. Ulloa, Peter J. Colias, Da Xu, Jamin Liu, Shaoni Mukhopadhyay, Bernardo D. Moreno, Amy S. Y. Lee, Sarah E. Chasins, James K. Nuñez

## Abstract

DNA methylation is the most abundant epigenetic modification on the human genome and changes dynamically during embryogenesis, aging, and disease. How DNA methylation is established *de novo* remains incompletely understood. Using orthogonal CRISPR platforms to initiate *de novo* DNA methylation coupled with genome-wide CRISPR screens and proximity-dependent proteomics, we discover the interplay between RNA m6A modifications and *de novo* DNA methylation to program heritable epigenetic memory in human cells. The RNA m6A writer complex is recruited to chromatin with DNA methyltransferases and histone repressors during the transient establishment phase of heritable epigenetic silencing. Inhibition of RNA m6A during early pluripotent transition leads to defects in *de novo* DNA methylation, resulting in upregulation of transposable elements that pose a threat to genome integrity during early development. Our work reveals the vital coordination between the epitranscriptome and *de novo* DNA methylation to establish proper heritable epigenetic silencing in human cells.

## INTRODUCTION

Precise regulation of gene expression is essential for cellular identity, lineage differentiation, and genomic stability. Human cells maintain genome integrity partly by enforcing heterochromatin, a condensed, transcriptionally silent state at genomic sites encoding DNA repeats, lineage-specific genes, and transposable elements (TEs). DNA methylation of cytosines is the most prevalent epigenetic chemical modification on the human genome and coats constitutive heterochromatic domains ^1–3^. DNA methylation undergoes dynamic changes during early human development and human aging, and is misregulated in diseases such as cancer and neurodevelopmental disorders ^4,5^. How human cells dictate which genomic sites to catalyze DNA methylation remains incompletely understood.

One of the most dynamic changes in DNA methylation patterning in humans occurs during early embryogenesis ^5^. After fertilization, the human genome undergoes global epigenetic resetting by first erasing the majority of DNA methylation in pre-implantation blastocysts, followed by *de novo* DNA methylation establishment after the blastocyst implants into the uterus ^6,7^. The period of low DNA methylation is accompanied by the expression of TEs that pose a threat to genome integrity ^8^. For example, human endogenous retrovirus K (HERV-K) is the youngest class of transposons in the human genome with ∼100 insertions that encode full-length proviral copies. HERV-K virus-like particles are produced in pre-implantation embryos, and then silenced upon *de novo* DNA methylation after implantation ^9,10^. Most recently, HERV-K activation has been shown to be essential for proper embryonic development ^11^.

Epigenetic silencing of HERVs is initiated by KRAB zinc finger proteins (ZFPs) that recruit the TRIM28 corepressor and SETDB1 to catalyze trimethylation of histone H3 lysine 9 (H3K9me3), which hallmarks constitutive heterochromatin ^12,13^. H3K9me3 is closely associated with DNA methylation, which is catalyzed by the *de novo* DNA methyltransferases DNMT3A and DNMT3B, and DNMT1 sustains epigenetic memory by heritable maintenance during cell division ^5,14–16^. Furthermore, accumulating evidence suggests coupling of H3K9me3 and DNA methylation in a reinforcing feedback loop ^17–23^. For instance, in mouse embryonic stem cells, H3K9 methylation is required for the proper establishment of DNA methylation at specific retrotransposons ^18,19,24,25^. Although the specific writers, erasers, and readers of H3K9me3 and DNA methylation are known, the upstream signaling pathways and auxiliary factors that coordinate the interplay and simultaneous deposition of these heritable marks are not fully understood ^13,26,27^.

TEs account for over 50% of the human genome, thus making it challenging to study the molecular steps governing their *de novo* epigenetic silencing at each genomic site ^28^. Bypassing this widespread complexity requires the ability to manipulate individual loci in isolation. Recent advances in targeted epigenome editing have enabled programmable epigenetic changes at defined genomic loci by tethering a DNA-binding domain, such as catalytically dead dCas9, to chromatin effectors to alter the local epigenetic environment of target genes ^18,29–32^. For example, synthetic co-targeting of DNMT3A, DNMT3L, and KRAB to a specified gene promoter induces *de novo* establishment of DNA methylation and H3K9me3 that is maintained by cells heritably through cell division and cell differentiation, harnessing the natural KRAB-TRIM28-SETDB1 signaling pathway for silencing HERVs ^33^. Recently, we pioneered the CRISPRoff technology that fuses dCas9 to DNMT3A, DNMT3L, and KRAB into one polypeptide, thereby allowing us to initiate *de novo* heritable epigenetic silencing at any site on the human genome with only two components: CRISPRoff and sgRNA ^34^.

Here, we aimed to understand the molecular steps that underlie *de novo* DNA methylation and downstream heritable epigenetic memory in human cells. Using orthogonal CRISPR platforms for epigenome editing, genome-wide screens, proximity-dependent proteomics, as well as pluripotent stem cell models of *de novo* DNA methylation, we discover a surprising interplay between proteins that regulate RNA m6A and *de novo* DNA methyltransferases to program heritable epigenetic silencing in human cells. Our work illuminates the coordination between RNA epitranscriptomics and *de novo* DNA methylation, particularly in epigenetic repression of transposable elements during early embryonic development.

## RESULTS

### Inducible epigenetic silencing by CRISPRi and CRISPRoff

To identify genes that are required for establishing short-term (CRISPRi) and long-term (CRISPRoff) epigenetic silencing (**Figure 1A**), we took advantage of our previously engineered doxycycline (dox) inducible CRISPRi (iCRISPRi) and CRISPRoff (iCRISPRoff) in K-562 leukemia cell lines that express sgRNAs targeting the promoter of the endogenous gene *CD55* (**Figures 1B** and **S1A**) ^34^. CD55 is nonessential for cell viability, localizes to the cell surface, and its protein levels can be measured quantitatively in single cells by antibody staining in live cells. Dox-induction of iCRISPRi and iCRISPRoff cells results in CD55 protein depletion in almost every cell by 72 hours after dox treatment (**Figure 1C**).

**Figure 1.**
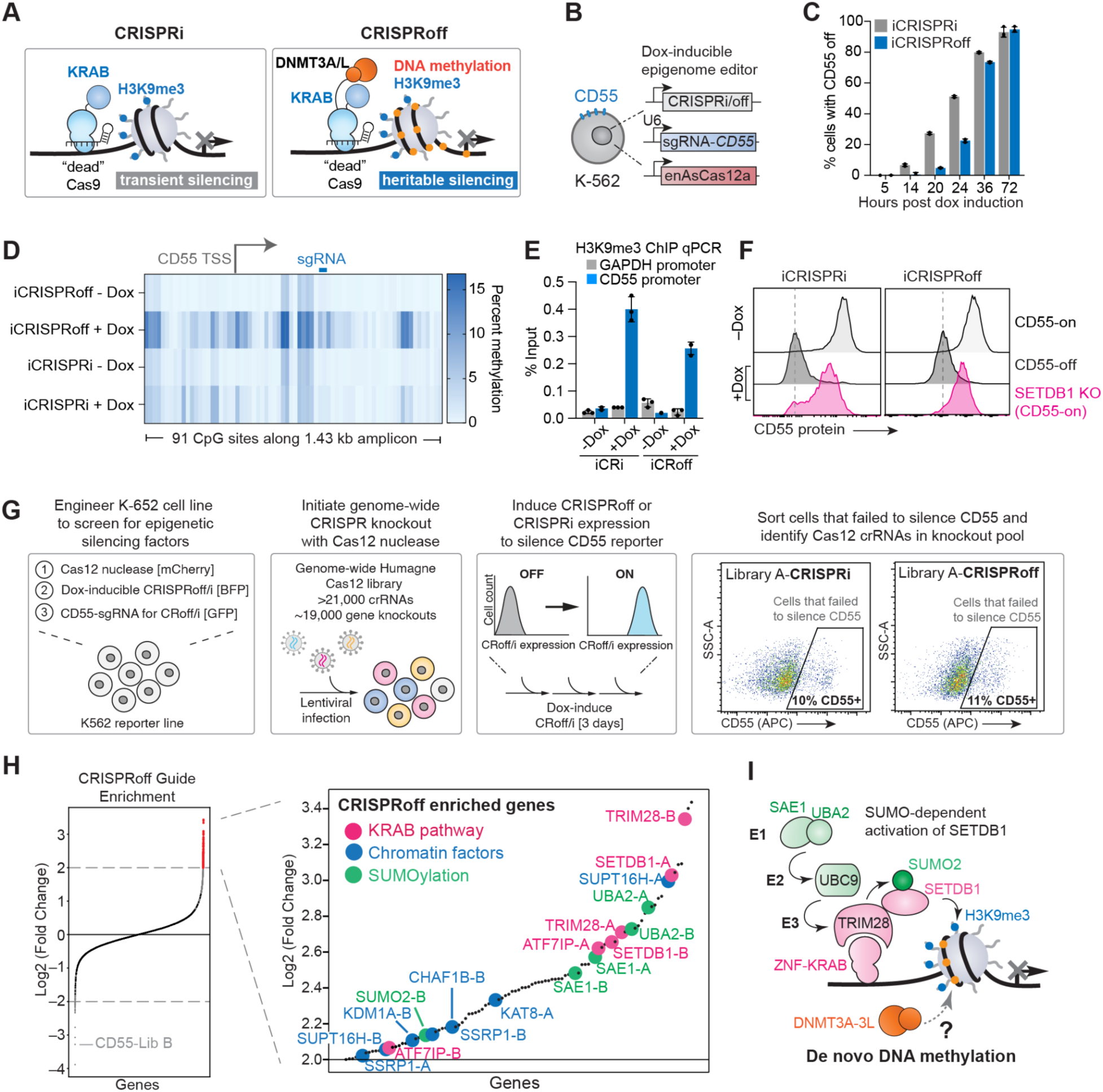
Genome-wide CRISPRoff screens identify regulators of heritable epigenetic silencing. (A) Schematic of CRISPRi and CRISPRoff epigenetic editors and their effects on chromatin. (B) Schematic of the engineered cell line used for the genome-wide CRISPRoff screen. (C) Time course of CD55 silencing following doxycycline-induced expression of CRISPRi (gray) or CRISPRoff (blue) editors in K-562 cells. (D) DNA methylation profiling across CpG sites at the *CD55* promoter in the absence of doxycycline and after 3 days of doxycycline-induced editor expression. (E) H3K9me3 ChIP-qPCR analysis day 3 post-dox induction at the *GAPDH* promoter (gray, negative control) and the *CD55* promoter (blue) following induction of CRISPRi or CRISPRoff. (F) Flow cytometry histograms of CD55 protein levels in single cells in the absence of doxycycline (light gray), after 72 hours of doxycycline treatment (dark gray), and after 72 hours of doxycycline treatment in SETDB1 knockout cells (pink). (G) Schematic of the genome-wide CRISPRi/off CRISPR screens, including representative flow cytometry plots from the Library A screening populations at the time of cell sorting. (H) Log2 fold-change (L2FC) ranking of sgRNAs enriched in the CRISPRoff failure-to-silence population relative to the unsorted population. sgRNAs are colored by functional pathways, including KRAB-associated factors, chromatin regulators, and SUMOylation. (I) Model of SUMO-dependent pathway required for KRAB-mediated epigenetic silencing.

We next validated that induction of CRISPRi and CRISPRoff leads to deposition of repressive epigenetic marks at the *CD55* promoter. First, we performed targeted enzymatic methyl (EM)-seq to profile DNA methylation changes within a 1.5-kb region of the promoter around the CD55-sgRNA binding site. While uninduced cells have little to no DNA methylation, induction of CRISPRoff establishes DNA methylation at the promoter (**Figures 1D** and **S1B**)^35^. We also detect DNA methylation upon CRISPRi induction albeit at lower levels, likely due to the recruitment of endogenous DNA methyltransferases (DNMTs) upon KRAB-mediated establishment of H3K9me3. Chromatin immunoprecipitation followed by quantitative PCR (ChIP-qPCR) confirms the enrichment of H3K9me3 at the *CD55* locus upon CRISPRi and CRISPRoff induction (**Figure 1E**).

Next, we engineered the inducible lines to constitutively express the engineered CRISPR-Cas12 nuclease from *Acidaminococcus* (enAsCas12a) that allows us to perform targeted genetic knockouts to determine the dependency factors of CRISPRi and CRISPRoff ^36,37^. We posited that knockout of *SETDB1* – the methyltransferase enzyme that catalyzes H3K9me3 deposition – would fail to establish CRISPRi and CRISPRoff-mediated silencing. Indeed, addition of *SETDB1*-targeting CRISPR RNAs (crRNAs) into iCRISPRi and iCRISPRoff cell lines severely impacts CD55 silencing upon dox induction (**Figures 1F** and **S1C**). Together, our engineered orthogonal CRISPR cell-based platform can quantitatively establish epigenetic repression at an endogenous locus, thus providing us with a tractable platform to systematically dissect the establishment phase of heritable epigenetic silencing.

### CRISPR screens to identify factors required for establishing *de novo* epigenetic silencing

To identify genes that are important for CRISPRi and CRISPRoff-mediated epigenetic silencing, we performed parallel genome-wide CRISPR screens using the enAsCas12 Humagne Set A and Set B libraries that target 20,081 protein-coding genes ^37^. We first packaged the crRNA libraries in lentiviral particles, transduced into iCRISPRi and iCRISPRoff K-562 cells, selected for crRNA-expressing cells with puromycin, and induced CRISPRi and CRISPRoff with doxycycline to silence CD55 (**Figures 1G** and **S1D**). After three days of continuous doxycycline-treatment to establish epigenetic silencing, we detected robust dCas9-editor expression as measured by BFP fused to the editors. CD55 protein staining resulted in 10-11% (iCRISPRi) and 11-15% (iCRISPRoff) of cells that failed to silence CD55 (**Figures 1G, S1E,** and **S1F**). We sorted cells that failed to silence *CD55*, sequenced the crRNAs, and compared the guides in the enriched population to those in the total unsorted population.

We ranked our genetic hits by calculating the log2 fold-change of each crRNA from the Set A and B libraries for the CRISPRi and CRISPRoff screens (**Tables S1** and **S2**). Among the strongest enriched genetic hits for both CRISPRi and CRISPRoff are members of the KRAB-mediated pathway, such as SETDB1, TRIM28, and ATF7IP (**Figures 1H** and **S1G**). Notably, our CRISPRoff genetic hits recapitulate the epigenetic repression pathway by identifying members of the SUMOylation pathway that is required for TRIM28-mediated SUMOylation of SETDB1 to stimulate its H3K9me3 methyltransferase activity (**Figure 1I**). Additionally, we identified factors that are implicated in heterochromatin formation, including the FACT complex (SSRP1-SUPT16H), a component the CAF-1 histone chaperone (CHAF1B), and the KDM1A/LSD1 histone demethylase ^18,38–40^.

To validate the genetic hits, we selected the top 100 hits with an enrichment score greater than 4 from the primary screens and performed a secondary screen in an arrayed format (one knockout per well). We chose crRNAs from both Set A and B libraries as our primary screens revealed crRNA-dependent activities with each library (**Figures 1H** and **S1G**). We repeated the iCRISPRi and iCRISPRoff approach to silence CD55 in K-562 cells and calculated a failure to silence score (FSS) by dividing CD55 protein expression in cells treated with doxycycline by that without doxycycline (**Figure 2A**). This approach enabled us to identify false-positive knockout genes from our primary screen that activate our CD55 reporter irrespective of CRISPRi/off induction, such as the mRNA polyA tail factor *NUDT21* that significantly increases CD55 levels upon knockout and thus yields a low FSS (**Figure S2A**). In contrast, knockout of genes in the KRAB pathway (*TRIM28*, *ATF7IP*, *SETDB1*) and SUMOylation (*SUMO2*, *UBA2*, *SAE*) do not increase CD55 levels in the absence of CRISPRi or CRISPRoff induction. However, their knockouts negatively impact CRISPRi- or CRISPRoff-mediated repression of CD55 that yields higher FSS, which indicates their role in establishing epigenetic silencing (**Figures 2B, 2C,** and **S2B**).

**Figure 2.**
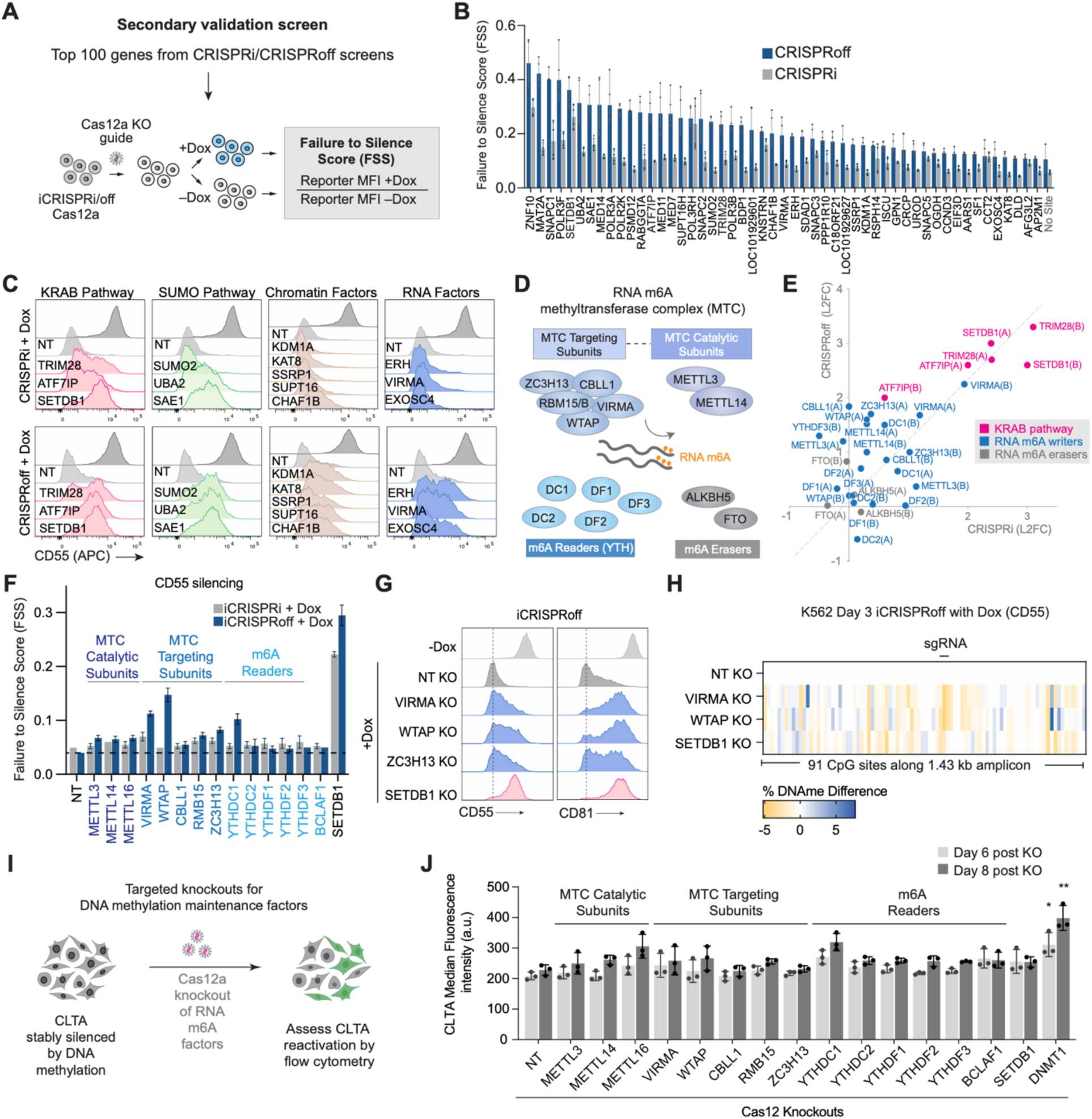
The RNA m6A pathway is involved in *de novo* DNA methylation. (A) Schematic illustrating the secondary validation screen workflow and calculation of the Failure-to-Silence Score (FSS) for each knockout gene. (B) Comparison of FSS values following knockout of candidate genes in the inducible CRISPRi and CRISPRoff cell reporters. Only factors exhibiting FSS values greater than the “No Site” CRISPR RNA negative control are shown. Individual data points indicate biological replicates; data are represented as mean ± s.d. (C) Representative flow cytometry histograms from inducible CRISPRi and CRISPRoff cell lines following knockout of candidate genes, grouped by functional pathways. (D) Schematic of RNA m6A pathway components, categorized into methyltransferase complex (MTC) targeting subunits, catalytic subunits, YTH domain-containing m6A readers and m6A erasers. (E) Relative log2 fold-change (L2FC) scores of KRAB-associated factors (pink), and RNA m6A writers (blue) and erasers (grey) from the CRISPRi and CRISPRoff primary screens. (F) Comparison of FSS values for CD55 expression following knockout of MTC targeting components and RNA m6A reader proteins in CRISPRi (grey) and CRISPRoff (blue) secondary validation screens. (G) Flow cytometry histograms of CD55 and CD81 following knockout of NT (non-targeting), MTC targeting components (VIRMA, WTAP, and ZC3H13), and the H3K9me3 methyltransferase (SETDB1). (H) DNA methylation profiling showing relative gains or losses of DNA methylation across the *CD55* promoter in NT, VIRMA, WTAP, and SETDB1 knockout in K-562s. (I) Schematic of the maintenance assay in CLTA-GFP HEK cells. (J) Quantification of CLTA reactivation after knockout of m6A factors, SETDB1, and DNA methylation maintenance factor DNMT1 in CLTA-silenced cells. \**p* < 0.05, \*\**p* < 0.01, unpaired t-test. n = 3 biological replicates, error bars ± 1 s.d.

We next analyzed genes that were more enriched in CRISPRoff compared to CRISPRi to select candidate factors that potentially interplay with *de novo* DNA methylation establishment (**Figures 2B** and **S2C**). The CRISPRoff FSSs are consistently higher than CRISPRi with notable exceptions such as positive control *SETDB1* (high FSS in CRISPRoff and CRISPRi) and negative control crRNAs (weak in both CRISPRoff and CRISPRi). Among the top genes with the strongest FSS for CRISPRoff compared to CRISPRi are *PSMD12* (proteasomal subunit), *SUMO2* (SUMOylation), and *MAT2A*, the catalytic subunit for S-adenosylmethionine (SAM) synthesis that is required for both DNA methylation and H3K9me3 processes (**Figure S2D**). Similar to the primary screen, chromatin factors KDM1A, SSRP1, SUPT16H, and CHAF1B yield stronger phenotypes with CRISPRoff compared to CRISPRi (**Figure 2C**).

#### The RNA m6A writer complex participates in establishing heritable epigenetic silencing

Surprisingly, we observed that knockout of genes that are implicated in RNA-centric pathways yields strong phenotypes with CRISPRoff compared to CRISPRi: *EXOSC4* (RNA degradation), *ERH* (RNA regulation), and *VIRMA* (RNA m6A modification) (**Figure 2C**) ^41–44^. We focused our study on RNA m6A as it is the most abundant epitranscriptomic modification in human cells and plays a critical role in RNA stability, degradation, and protein synthesis ^45,46^.

VIRMA is a subunit of the m6A methyltransferase complex (m6A writer complex or MTC), a multi-protein complex consisting of the MAC catalytic subunits (METTL3, METTL14) and MACOM (VIRMA, WTAP, CBLL1, ZC3H13) ^44,47–50^. The m6A modification is recognized by reader YTH reader proteins (YTHDC1, YTHDC2, YTHDF1, YTHDF2, YTHDF3), which can act redundantly to process the RNA^51–53^ (Figure2D). Our primary CRISPR screens yielded enrichment of players in the RNA m6A pathway, especially the subunits of the MTC (VIRMA, WTAP, ZC3H13, CBLL1) and the catalytic subunits METTL3 and METTL14 (**Figure 2E**).

Recent work in mammalian cells has shown that TE transcripts are highly enriched in RNA m6A, which contributes to their epigenetic silencing through H3K9me3 deposition ^54–61^. Moreover, METTL3-METTL14 can recruit DNMT1 to regulate gene body methylation and gene expression ^62^. However, the role of RNA m6A in mediating *de novo* DNA methylation has not been described. To determine if the RNA m6A pathway is required for establishing *de novo* epigenetic silencing in human cells, we individually knocked out members of the MTC and readers in K-562 cells and induced expression of CRISPRi and CRISPRoff to silence *CD55*. Indeed, knockouts of the MTC subunits *VIRMA* and *WTAP* and the *YTHDC1* reader leads to strong defects in CD55 silencing (**Figures 2F, 2G**, **S3A,** and **S3B**). We additionally performed DNA methylation profiling of the *CD55* promoter in *VIRMA* and *WTAP* knockout cells followed by CRISPRoff induction, which show strong defect in *de novo* DNA methylation at levels similar to *SETDB1* knockout cells (**Figures 2H** and **S3C**). Together, our data point to the role of the RNA m6A pathway in mediating the *de novo* establishment of heritable epigenetic silencing in human cells.

### RNA m6A dependency is consistent across different gene targets and cell types

We next tested whether *de novo* epigenetic silencing is dependent on the RNA m6A pathway in other genetic and cellular contexts. In K-562 cells, targeting CRISPRoff to a different endogenous gene (*CD81*) after knockouts of the RNA m6A players leads to defect in *de novo* epigenetic silencing in the *VIRMA*, *WTAP*, and *ZC3H13* knockouts (**Figures 2G** and **S3D-S3E**). Given that RNA m6A pathway regulates RNA stability, we sought to exclude the possibility that the observed defect in establishing epigenetic silencing is an artifact of destabilized transcript encoding the CRISPR editors or the accompanying sgRNA. Thus, we delivered CRISPRoff and sgRNAs as ribonucleoprotein complexes (RNP) packaged in virus-like particles to bypass the need for DNA or RNA expression of CRISPRoff ^63^ (**Figure S3F)**. In HEK293T cells, we first knocked out *VIRMA*, *WTAP,* and *SETDB1* and delivered CRISPRoff RNPs with sgRNAs targeting the endogenous *CLTA* gene to initiate *de novo* epigenetic silencing. The *CLTA* gene is tagged endogenously with GFP, which provides us with a quantitative readout of epigenetic silencing ^34,64^. Consistent with our data in K-562 cells, knockout of *VIRMA* and *WTAP* negatively affects *de novo* silencing of CLTA (**Figure S3G**). DNA methylation profiling of the *CLTA* promoter shows reduced *de novo* DNA methylation at the *CLTA* promoter in *VIRMA* and *WTAP* knockout cells, similar to the *SETDB1* knockout positive control (**Figure S3H**). Together, we observe that establishing *de novo* epigenetic silencing is dependent on the RNA m6A factors across different endogenous genes and cell types.

### RNA m6A factors are not important for maintenance of epigenetic silencing

DNMT3A and DNMT3B primarily catalyze *de novo* DNA methylation, whereas DNMT1 faithfully maintains DNA methylation during cell division ^26^. To determine whether the RNA m6A factors are also involved in the maintenance phase of epigenetic silencing, we performed a small-scale arrayed screen by individually knocking out the RNA m6A MTC subunits and readers in HEK293T cells with CLTA pre-silenced stably by CRISPRoff (**Figure 2I**). We passaged the cells for 8 days and quantified reactivation of CLTA expression by flow cytometry. Indeed, knockout of the maintenance DNA methyltransferase *DNMT1* reactivates CLTA expression. In contrast, knockout of RNA m6A factors do not significantly reactivate CLTA expression (**Figure 2J**). These studies suggest that the RNA m6A factors are not required for the maintenance of DNA methylation but instead play a role in the establishment phase of *de novo* epigenetic silencing.

### Proteomics detects RNA m6A factors at the establishment site of epigenetic silencing

CasID-mediated proteomics labeling of proteins at human endogenous retroviruses (HERV) genomic loci have identified the RNA m6A writer METTL3 and m6A readers YTHDC1, YTHDF1, along with chromatin factors involved in heterochromatin such as HP1α, HP1β, DNMT1, and KRAB-ZFPs ^65^. Additionally, ERV reporter-based screen assays and co-immunoprecipitation of Mettl3 in mouse ES cells have shown that Setdb1 interacts with the Yth reader proteins and Mettl3, suggesting that the RNA m6A factors co-localize with chromatin repressors to maintain epigenetic silencing ^54,61^. We next sought to probe the proteins that localize to a genomic site during the transient establishment phase of *de novo* epigenetic silencing.

Previously, we performed proximity-dependent labeling of KRAB interactors by fusing APEX2 to the N-terminus KRAB-dCas9, transfecting the construct in HEK293T cells to initiate epigenetic silencing of the *CLTA* promoter, followed by biotin labeling of proximal proteins 24 hours post-delivery into cells (**Figure 3A**) ^66^. Analysis of the data yields strong enrichment of the RNA m6A writer complex and readers, including members of the RNA m6A writer complex METTL3, METTL14, VIRMA, WTAP, ZCH3H13 and RBM15B, along with histone repressors and DNA methyltransferases (**Figures 3B** and **S4A**; **Table S3**). To further validate our findings, we performed an independent APEX2 labeling with KRAB-APEX2 at the C-terminus of dCas9. In agreement with the N-terminal APEX2-KRAB dataset, we detect enrichment of known KRAB interactors (e.g. TRIM28), DNMT3A and the RNA m6A writer complex subunits WTAP, VIRMA, and ZC3H13 (**Figures S4B** and **S4C**; **Table S4**).

**Figure 3.**
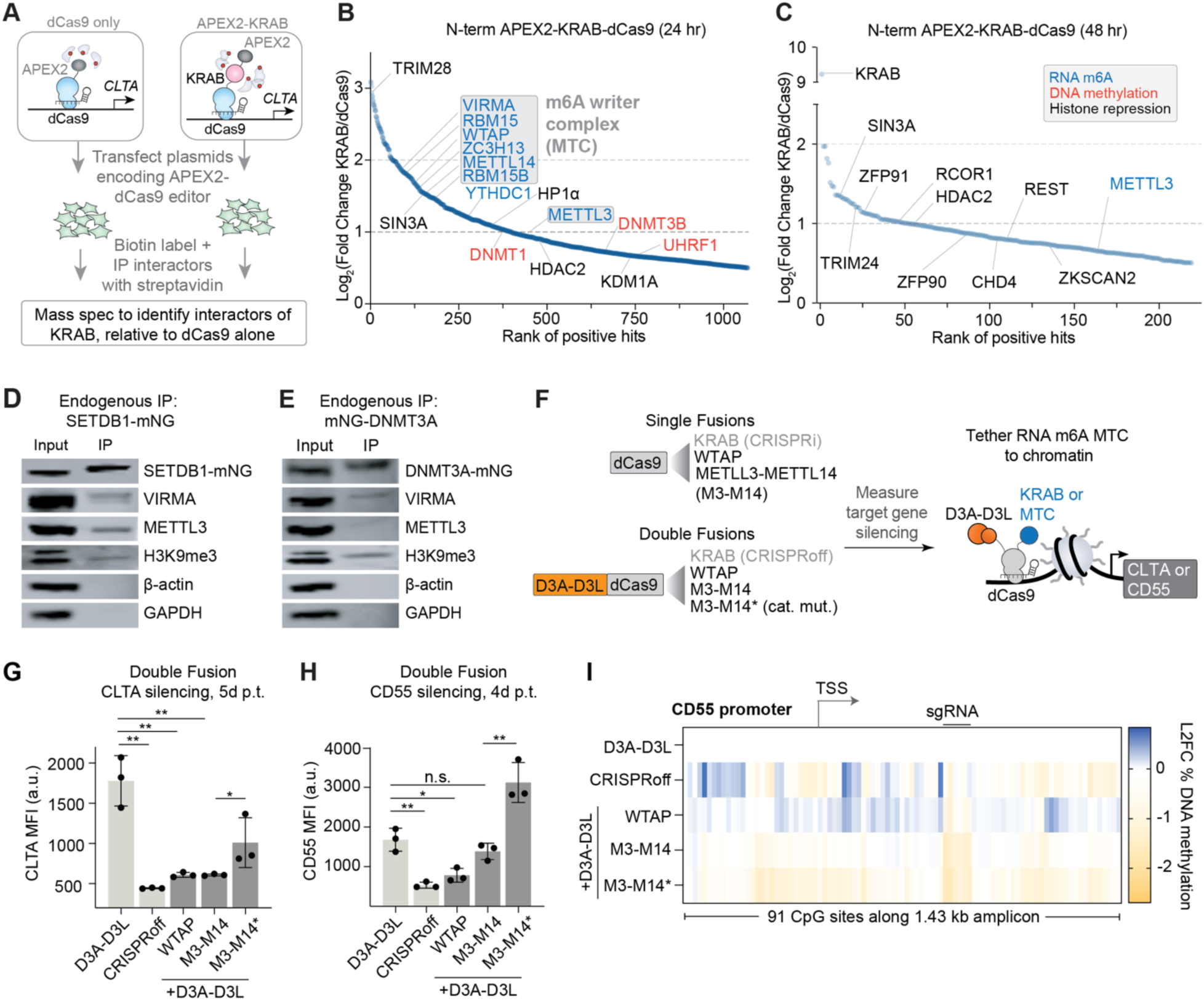
Co-localization of RNA m6A factors with chromatin repressors during the establishment phase of *de novo* epigenetic silencing. (A) Schematic illustrating APEX2 proximity-dependent labeling approach to identify protein interactors of KRAB in comparison to a negative control dCas9-only condition. (**B** and **C**) Rank-based plot of KRAB interactors measured at 24 hours (**B**) and 48 hours (**C**) post-transfection of plasmids encoding APEX2-KRAB-dCas9. Proteins in the RNA m6A pathway are highlighted in blue, including members of the RNA m6A writer complex, pink for DNA methylation proteins, and black for histone repression. (n = 2 biological replicates) (**D** and **E**) Immunoprecipitation of SETDB1 (**D**) and DNMT3A (**E)** tagged endogenously with mNeonGreen in HEK293T cells. Western blots are shown for SETDB1, TRIM28, and H3K9me3, VIRMA, METTL3, and negative controls β-actin and GAPDH. (F) Schematic of single and double fusion dCas9 constructs designed to target endogenous reporters (CLTA or CD55). D3A and D3L represents DNMT3A and DNMT3L, respectively. M3-M14 represents catalytic domains of METTL3 and METTL14; M3-M14* represents catalytic-dead METTL3 catalytic domain fused with METTL14 catalytic domain. (**G** and **H)** Bar plots showing median fluorescence intensity (MFI) of CLTA-GFP 5 days post-transfection (**G**) and CD55 4 days post-transfection (**H**) with double fusion constructs. n.s. non-significant, \**p* < 0.05, \*\**p* < 0.01, unpaired t-test. n = 3 biological replicates, error bars ± 1 s.d. (I) DNA methylation profiling showing relative gains or losses of DNA methylation across the *CD55* promoter and transcription start site (TSS) upon targeting with dCas9 double fusions, normalized to D3A-3L-dCas9 only construct.

A time course analysis of CLTA silencing assayed by flow cytometry after transfecting APEX2-dCas9 alone or APEX2-KRAB-dCas9 shows that CLTA protein levels remain high at 24 hours post-transfection and become fully depleted by 72 hours (**Figure S4D**). To gain insights into the recruitment dynamics at the establishment site of *de novo* epigenetic silencing, we performed APEX2 labeling at 48 hours post-delivery of APEX2-KRAB-dCas9 or APEX2-dCas9 and downstream biotinylation and mass spectrometry to determine the proteins that are at the *CLTA* promoter. GO term analysis shows a clear reduction in proteins localized to the establishment site by 48 hours compared to 24 hours (**Figure S4E**). In particular, we no longer detect significant enrichment of TRIM28, DNMTs or the RNA m6A factors at 48 hours despite the strong labeling of the KRAB fused to APEX2, suggesting that the *de novo* establishment phase has completed (**Figures 3C** and **S4F**; **Table S5**). These data suggest that the co-localization of the RNA m6A writer complex, *de novo* DNMTs and histone repressors occurs during the establishment phase of *de novo* epigenetic silencing.

Finally, we performed immunoprecipitation experiments to further validate the co-localization of the RNA m6A factors with histone repressors and *de novo* DNA methyltransferases. First, we generated HEK293T cell lines with *SETDB1* or *DNMT3A* tagged endogenously with mNeonGreen (SETDB1-mNeonGreen and DNMT3A-mNeonGreen), followed by mNeonGreen immunoprecipitation (F**igures S4G** and **S4H**) ^67,68^. Immunoprecipitation of SETDB1 followed by Western blot analysis confirms the presence of VIRMA, METTL3, YTHDF2, along with positive controls TRIM28 and H3K9me3 (**Figures 3D** and **S4I**). Immunoprecipitation of DNMT3A detects positive co-immunoprecipitation with VIRMA and H3K9me3 (**Figures 3E** and **S4J**). Altogether, our data support the RNA m6A factors coordinate with *de novo* DNA methyltransferases and histone repressors that together remodel the local chromatin environment to establish epigenetic silencing.

### Direct recruitment of RNA m6A MTC to chromatin induces epigenetic silencing

Our data show the RNA m6A readers and writers co-localize with *de novo* DNA methyltransferases to establish epigenetic silencing. We posit that direct tethering of the RNA m6A MTC and *de novo* DNA methyltransferases to gene promoters will result in epigenetic silencing, similar to direct co-localization of KRAB and DNMT3A-DNMT3L (D3A-D3L) in CRISPRoff. We constructed two classes of dCas9 fusions to MTC subunits, first as single fusions to WTAP and the METTL3-METTL14 catalytic domains, and second with WTAP or METTL3-METTL14 catalytic domains fused to D3A-D3L (**Figure 3F**). We transfected plasmids expressing each dCas9 fusion in HEK293T cells to silence *CLTA* or *CD55*, sorted transfected cells, and measured CLTA or CD55 protein levels in single cells to monitor silencing. As controls, we included dCas9 only, D3A-D3L-dCas9, dCas9-KRAB (CRISPRi), and a catalytically dead mutant version of METTL3 in the METTL3-METTL14 catalytic domain fusion.

Of the single dCas9 fusions, only CRISPRi results in near-complete CLTA silencing. The WTAP and METTL3-METTL14 single fusions performed similar to dCas9 only, suggesting that recruitment of the MTC alone does not signal downstream epigenetic silencing (**Figure S5A**). In contrast, the combinatorial fusions of D3A-D3L to WTAP or METTL3-METTL14 induce robust CLTA silencing at levels comparable to CRISPRoff. At 5 days post editor transfection, D3A-D3L-dCas9 silencing remains incomplete, whereas the D3A-D3L fusions to the MTC components induce near-penetrant CLTA silencing (**Figures 3G** and **S5B**). In experiments targeting CD55, we detect similar robust epigenetic silencing with the D3A-D3L fusions to WTAP or METTL3-METTL14 at levels similar to CRISPRoff (**Figures 3H**, **S5C,** and **S5D**). Of note, the catalytic mutant METTL3-METTL14 fusion to D3A-D3L results in weaker silencing efficiency of CLTA and CD55.

To determine if the measured epigenetic silencing correlates with *de novo* DNA methylation rather than steric hindrance of the transcriptional machinery, we performed DNA methylation profiling of the *CD55* and *CLTA* promoters at an early (day 3) and late (day 12) time point post-transfection of the editors to capture early establishment and maintenance of DNA methylation. As expected, all D3A-D3L fusions induce DNA methylation at the promoters at day 3 (**Figures S5E** and **S5F**). We normalized to DNA methylation levels to D3A-D3L-dCas9 to determine if adding the MTC enhances DNA methylation levels. Indeed, the D3A-D3L and WTAP combination induces higher levels of DNA methylation at both target promoters compared to D3A-D3L alone, which is heritable over subsequent cell divisions (day 12 post-transfection), especially at *CD55* (**Figures 3I, S5G,** and **S5H**). In contrast, the METTL3-METTL14 fusions to D3A-D3L do not enhance DNA methylation at the *CLTA* promoter and only at CpG sites near the sgRNA binding site at the *CD55* promoter. These data together support that direct co-recruitment of the RNA m6A writer complex with DNMT3A-DNMT3L results in enhanced *de novo* DNA methylation and ultimate establishment of heritable epigenetic silencing.

### Global interplay between RNA m6A and *de novo* DNA methylation during pluripotent stem cell transition

Our experiments thus far have relied on establishing epigenetic silencing with synthetic CRISPR epigenome editors. We next sought to determine if the interplay between RNA m6A and *de novo* DNA methylation occurs during a natural process of global establishment of *de novo* DNA methylation. Thus, we use a human pluripotent stem cell (hPSC) platform that mimics the global erasure and establishment of *de novo* DNA methylation, enabling us to dissect the functional interplay between RNA m6A and *de novo* DNA methylation in a dynamic process of global epigenetic resetting.

First, we transitioned human H9 hPSCs from primed to naive states, which leads to global erasure of DNA methylation (**Figures 4A** and **S6A-S6C**). Then, we differentiated naive hPSCs to primed hPSCs to initiate global *de novo* DNA methylation. We profiled DNA methylation at genomic loci encoding highly active TEs, LINE-1/L1Hs and HERV-K/LTR5_Hs, to determine the temporal dynamics of *de novo* DNA methylation establishment. A 24-day time course during repriming leads to gradual gain of *de novo* DNA methylation at the 5’ UTR region of LINE-1/ L1Hs and 5’LTR region HERV-K/LTR5_Hs genomic sites, as detected as early as day 4 post initiation of priming (**Figures 4B, 4C, S6B,** and **S6C**). To dissect the coordination of DNA methylation and RNA m6A, we treated naive hPSCs undergoing repriming with increasing doses of the RNA m6A inhibitor STM2457 for 4 days (**Figure S6D)**. We confirmed that STM2457 treatment during differentiation does not impair the transition from naive (CD7⁺) to primed (CD57⁺) hPSC state (**Figures S6E** and **S6F**).

**Figure 4.**
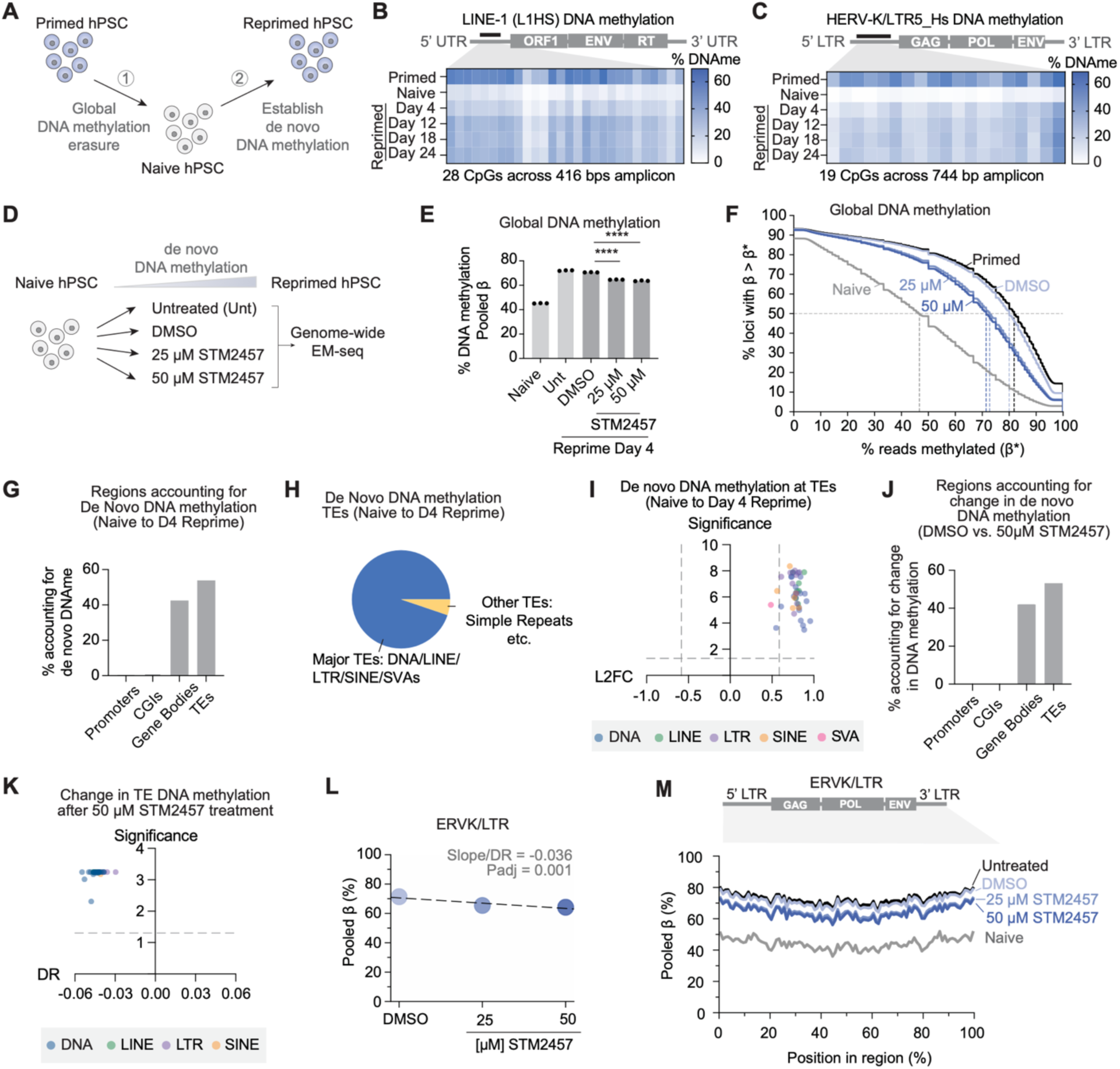
Coordination of RNA m6A and *de novo* DNA methylation for transposable element silencing during pluripotent stem cell transition. (**A**) Schematic illustrates the workflow of the human stem cells (H9) used to model the pre-implantation (naive) and post-implantation (prime) stages of development. (**B** and **C**) DNA methylation profiling/heatmaps showing percent methylation at each CpG at (**B**) L1Hs and (**C**) HERV-K/LTR5_Hs elements from naive H9 cells through 24 days of repriming. (D) Schematic of whole genome EM-seq treatment conditions during the transition from naive to primed hPSCs. (E) Bar graph showing pooled DNA methylation (β-values) in naive cells versus day 4 reprimed cells across the designated treatment conditions. \*\*\*\**p* < 0.0001, unpaired t-test. n = 3 biological replicates, error bars ± 1 s.d. (F) Survival plot indicating the percentage of genomic loci exceeding a specific methylation threshold (β*). Dashed vertical lines track the median methylation level (the 50% intercept on the y-axis) for naive, primed, DMSO-treated, and STM2457-treated populations. (G) Bar graph showing the relative percentage of *de novo* DNA methylation localized to major genomic features (Promoters, CpG Islands [CGIs], Gene Bodies, and TEs) during the naive to day 4 reprimed transition. (H) Pie chart representing the proportion of *de novo* methylation changes mapped specifically to major TE families (DNA transposons, LINEs, LTRs, SINEs, and SVAs) versus minor elements or simple repeats. (I) Volcano plot showing the statistical significance (-log10 of adjusted *p*-value) versus the log2 fold change (L2FC) of DNA methylation across individual TE families during the naive to day 4 reprimed transition. Points are color-coded by major TE class. (J) Bar graph illustrating the relative proportion of altered methylation localized across promoters, CGIs, gene bodies, and TEs when comparing the DMSO control and 50 µM STM2457 treatment conditions. (K) Volcano plot showing statistical significance (-log10 of adjusted *p*-value) versus the STM2457 dose response slope of TE classes. Points are color-coded by TE category. (L) Scatter plot showing the pooled β-values for ERV-K/LTR regions across increasing doses of STM2457. The regression slope/dose-response (DR) value (regression slope) and adjusted p-value are displayed. Repeatmasker annotates HERV as ERV. (M) Metagene plot highlighting DNA methylation levels (pooled β) mapped across the normalized length of the ERV-K/LTR consensus sequence in naive, untreated, DMSO, and dose-dependent STM2457 treatment groups. Repeatmasker annotates HERV as ERV.

We next performed whole-genome EM-seq to determine the global changes in *de novo* DNA methylation in hPSCs transitioning from naive to day 4 reprimed states (**Figure 4D**). Calculation of the pooled β-values (or average methylation level at cytosines weighted by read count) demonstrates that naive cells exhibit about 45% DNA methylation genome-wide, which increases to an average of 72% in the untreated reprimed states (**Figures 4E** and **4F**), thus confirming the global accumulation of *de novo* DNA methylation during the repriming process. Notably, STM2457 treatment disrupts the global gain of *de novo* DNA methylation. DMSO-treated cells increase to 70% methylation, while 50 µM STM2457-treated cells increase to only 63% methylation on day 4 of repriming (**Figures 4E** and **4F**).

We then identified statistically significant genomic regions that gained more than 1.5-fold methylation in the reprimed state relative to the naive state and mapped where these *de novo* events occurred in the genome. Gene bodies and transposable elements (TEs) account for most of the newly acquired DNA methylation (**Figure 4G**). Specifically, among the TEs targeted for silencing, major transposon and retrotransposon families, DNA transposons, LINEs, LTRs, and SINEs, constituted over 90% of the TE loci undergoing significant *de novo* methylation (**Figure 4H**). Further analysis confirms a statistically significant, positive fold change in methylation across different TE families during the transition from naive state to day 4 of repriming (**Figure 4I** and **S7A**).

Next, we sought to determine the effect on *de novo* DNA methylation in hPSCs transitioning from naive to reprimed in the presence of increasing doses of the STM2457 RNA m6A chemical inhibitor compared to cells treated with a vehicle control (DMSO) (**Figure 4D**). We determined a dose response (DR) to quantify the magnitude of DNA methylation change, calculated as the slope of pooled β-values across the DMSO, 25 μM, and 50 μM conditions (**Figure S7B**). Most of the dose-dependent reduction was concentrated within gene bodies and TE genomic sites (**Figure 4J**). While gene body dose responses showed some variance, repetitive elements showed uniform global loss in *de novo* DNA methylation establishment (**Figures 4K** and **S7C**). Strikingly, every examined sub-family of TEs failed to establish wild-type levels of *de novo* DNA methylation under increasing concentrations of STM2457. Specifically, HERV-K/LTR5 and LINE-1/L1 TE families show a dose response of -0.036 and -0.043, respectively (**Figures 4L, 4M, S7D,** and **S7E**). Beyond the LINE-1 and HERV-K families, we observed dose-dependent methylation deficits across a wide array of transposon subfamilies upon STM2457 treatment. DNA transposons (e.g., TcMar, hAT), various LINE classes (L1, L2, RTE), multiple LTR variants (HERV1, HERV-L, HERV-K, Gypsy), and SINE elements (tRNA, MIR) all exhibited a failure to gain methylation (**Figure S7F**). Collectively, this uniform failure across evolutionarily distinct repetitive sequences strongly suggests a critical global interplay wherein RNA m6A acts as a prerequisite for the establishment of *de novo* DNA methylation across the genome during pluripotent stem cell transition.

### RNA m6A mediates *de novo* methylation to epigenetically silence transposable elements

Our genomics data show that RNA m6A inhibition reduces *de novo* DNA methylation during embryonic differentiation, particularly at TE genomic loci. We next wondered whether these observations have a direct impact on transcriptional repression of TEs. Using the same naive to primed hPSC model to establish *de novo* DNA methylation, we profiled transcript levels, m6A enrichment on transcripts, and *de novo* DNA methylation of TEs in the presence or absence of the STM2457 RNA m6A inhibitor (**Figure 5A**). We focus on LINE and HERV-K elements as these TEs are transcriptionally active in pre-implantation embryos during global DNA methylation erasure.

**Figure 5.**
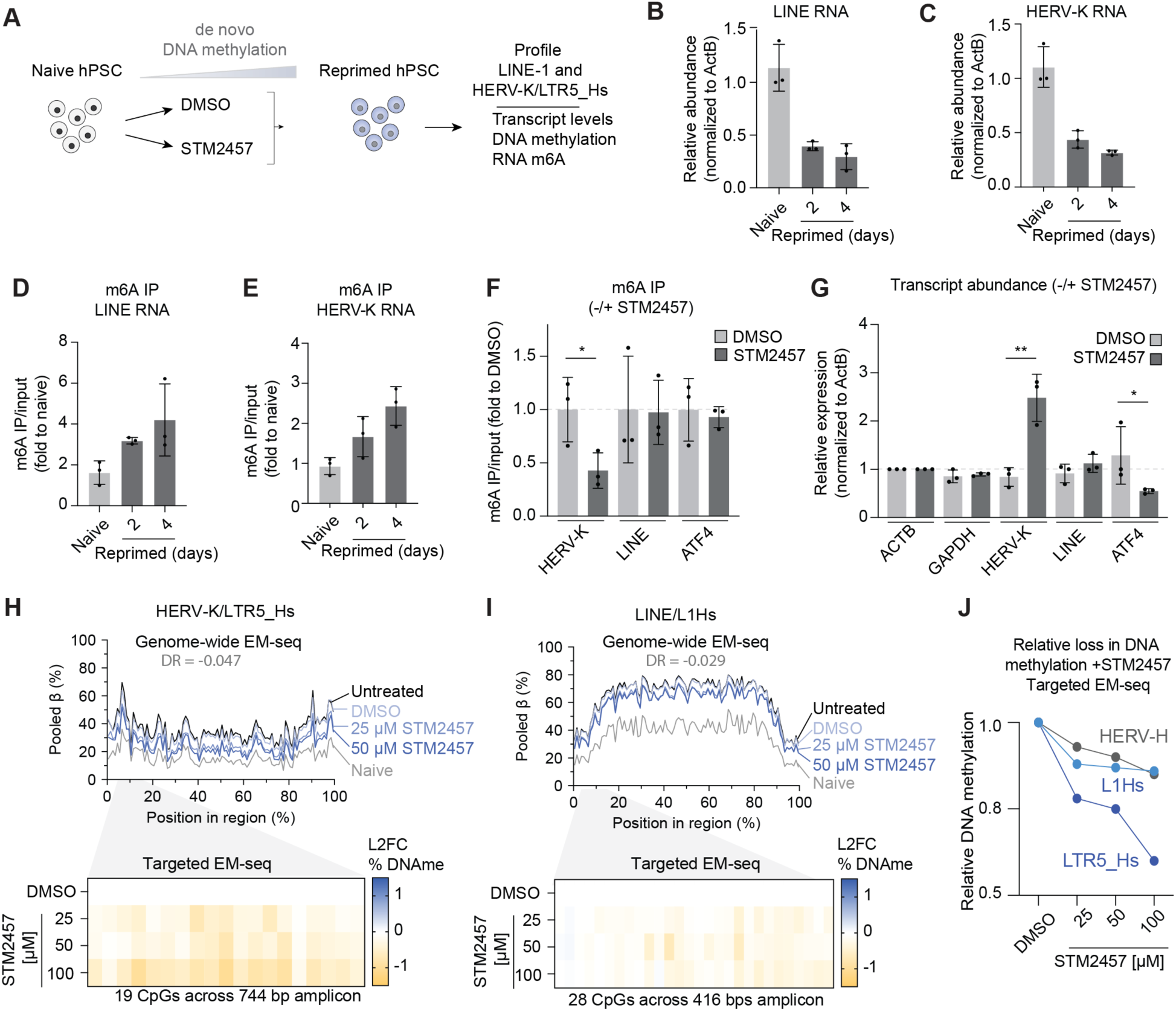
RNA m6A mediates *de novo* methylation to epigenetically silence transposable elements. (A) Schematic of experimental workflow where naive hPSCs are transitioned to reprimed hPSCs in the presence of either DMSO (vehicle control) or METTL3 inhibitor STM2457. Downstream analysis profiled LINE-1 and HERV-K dynamics across three regulatory layers: transcript abundance, DNA methylation and RNA-m6A levels. (**B** and **C**) Reverse transcription-quantitative PCR (RT-qPCR) analysis of LINE-1 (**B**) and HERV-K (**C**) RNA levels in H9 hPSC naive, D2 reprimed, and D4 reprimed cells (n = 3 biological replicates). (**D** and **E**) Level of m6A in LINE1 (**D**) and HERV-K (**E**) RNA measured by m6A RNA immunoprecipitation (m6A IP) and RT-qPCR in H9 hPSC naive, D2 and D4 reprimed cells (n = 3 biological replicates). (**F**) Level of m6A on LINE-1 and HERV-K RNAs, alongside a known m6A-modified ATF4 RNA measured by m6A IP and RT-qPCR in H9 hPSC D4 reprimed cells treated with DMSO and 50 μM STM2457 for 72 hours. \**p*<0.05, one-sided t-test; n = 3 biological replicates. (**G**) RT-qPCR analysis of LINE-1 and HERV-K RNA levels, alongside a known m6A-modified ATF4 RNA in H9 hPSC D4 reprimed cells treated with DMSO and 50 μM STM2457 for 72 hours. \**p*<0.05, **p<0.01, one-sided t-test; n = 3 biological replicates. (**H** and **I**) Metagene profiles from genome-wide EM-seq tracking pooled β (%) across the normalized length of the HERV-K/LTR5_Hs (**H**) and LINE/L1Hs (**I**) elements, and heatmaps depicting targeted EM-seq under naive, untreated, DMSO, and dose-dependent STM2457 conditions. Dose-response (DR) shifts are shown. (J) Line plot summarizing the relative loss in DNA methylation calculated via targeted EM-seq for HERV-H, L1Hs, and LTR5_Hs across the concentration gradient of STM2457. Values are normalized relative to the DMSO control baseline.

First, we measured the relative RNA levels of LINE and HERV-K transcripts by reverse transcription quantitative PCR (RT-qPCR) during the repriming process. Indeed, the naive to reprimed transition coincides with reduced levels of LINE and HERV-K transcripts on days 2 and 4 of priming, which confirms transcriptional silencing of these TEs during global DNA methylation establishment. (**Figures 5B** and **5C)**. We then performed immunoprecipitation of transcripts marked by m6A (m6A-IP), followed by RT-qPCR to determine the m6A enrichment on LINE and HERV-K transcripts. We detect increased m6A on LINE and HERV-K transcripts during the naive to primed transition, in line with RNA m6A being associated with reduced transcript levels of these TE classes (**Figures 5D** and **5E**).

We next treated naive hPSCs with STM2457 or DMSO during differentiation to determine the levels of LINE and HERV-K transcripts upon RNA m6A inhibition. RNA m6A-IP confirms marked reduction in RNA m6A on HERV-K upon STM2457 treatment (**Figures 5F** and **S8A**). Concurrent with depletion of RNA m6A, we detect significantly high abundance of HERV-K transcripts upon RNA m6A inhibition whereas LINE transcript levels are less sensitive to STM2457 treatment (**Figure 5G**). Lastly, we performed targeted DNA methylation profiling at the promoter region of the most active subset of HERV-K and LINE elements, LTR5_Hs and L1Hs, respectively, during repriming in the presence of increasing doses of STM2457. We observe a significant defect in establishing *de novo* DNA methylation at HERV-K/LTR5_Hs upon RNA m6A inhibition (**Figures 5H** and **S8B**). In comparison, *de novo* DNA methylation at L1Hs and another HERV element, HERV-H, are less affected upon increasing doses of STM2457 (**Figures 5I, S8D,** and **S8E**). Notably, we compared our targeted DNA methylation profiling at the LTR5_Hs and L1Hs promoter regions with our genome-wide EM-seq data of the full-length copies of HERV-K and L1Hs, which confirms more marked reduction in DNA methylation at LTR5_Hs compared to L1Hs (**Figures 5H-5J**).

Finally, we sought to determine whether the interplay between RNA m6A and *de novo* DNA methylation in hPSCs occurs in the context of statically dividing cells (epigenetic maintenance) instead of *de novo* epigenetic establishment. Thus, we treated primed H9 hPSCs with increasing doses of STM2457 (**Figure S8F**). DNA methylation profiling at TE genomic sites shows non-significant changes at L1Hs, HERV-K/LTR5_Hs, and HERV-H loci with increasing STM2457 concentrations (**Figures S8G-S8I**). Thus, we posit that the interplay between RNA m6A and *de novo* DNA methylation occurs in the context of establishing epigenetic silencing rather than maintaining a pre-established epigenetically silenced state.

## DISCUSSION

Here, we employ two independent approaches to study *de novo* epigenetic silencing (CRISPR epigenome editing and pluripotent stem cell state transition) to uncover a novel mechanism that bridges RNA m6A and *de novo* DNA methylation to program heritable epigenetic silencing in human cells. We posit that this mechanism serves as a critical genomic surveillance system to repress evolutionarily TEs during global establishment of DNA methylation following a period of low DNA methylation in pre-implantation embryonic cells, thereby ensuring proper heritable silencing of TEs that pose a threat to genomic stability (**Figure 6**).

**Figure 6.**
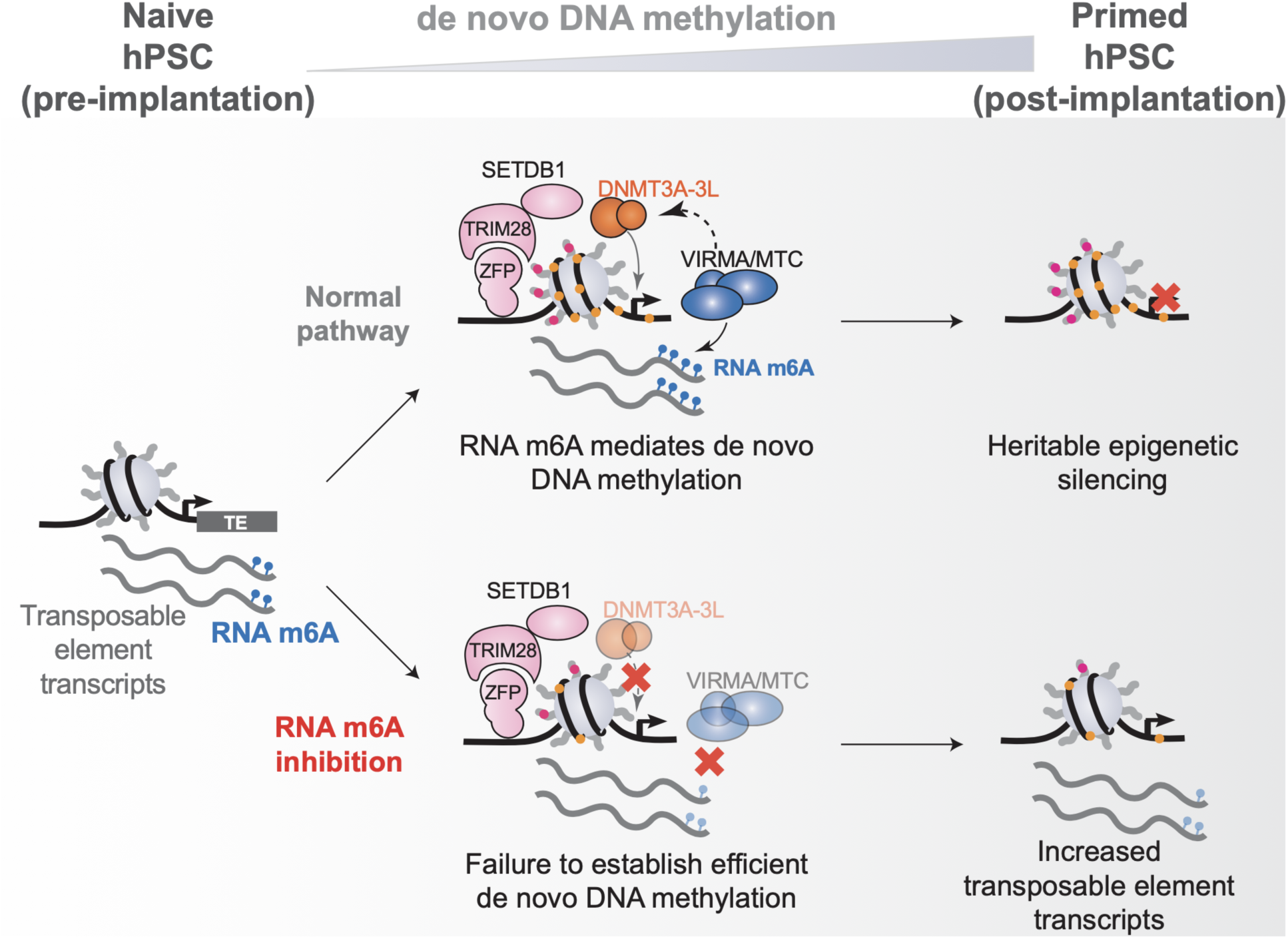
Model of the interplay between RNA m6A and *de novo* DNA methylation to repress TEs during embryonic differentiation. The RNA m6A writer complex adds RNA m6A to TEs and coordinates with *de novo* DNA methyltransferases and chromatin repressors to establish epigenetic silencing during the dynamic phase of epigenetic resetting in early embryonic development. In the absence of MTC/VIRMA or RNA m6A, efficient *de novo* DNA methylation establishment is affected negatively, leading to upregulation of TE transcripts.

Recent work has demonstrated crosstalk between the RNA m6A pathway and DNA methylation via a signaling axis between the RNA m6A catalytic subunits METTL3-METTL14 and DNMT1 to regulate gene body DNA methylation and gene expression ^62^. DNMT1 functions primarily in faithful maintenance of DNA methylation during cell division, whereas our work focuses on *de novo* DNA methylation catalyzed by DNMT3A and DNMT3B. As such, RNA m6A chemical inhibition did not affect pre-existing DNA methylation at HERV-K and L1Hs genomic loci in statically dividing primed hPSCs or at the promoters of genes that were synthetically DNA methylated by CRISPRoff. Instead, we show that the cooperation between RNA m6A and *de novo* DNA methylation is selectively required for the dynamic establishment phase of silencing. Our data highlight the broad coordination of RNA m6A and DNA methylation to regulate gene expression and the *de novo* establishment of epigenetic silencing.

TEs are among the most enriched transcripts marked by RNA m6A modifications in mammalian cells, which can function as key signals for controlling TE transcript levels through crosstalk with epigenetic modulators ^54–56,59–61,69^. Moreover, the RNA m6A writers and erasers co-localize with chromatin repressors at human endogenous retrovirus genomic loci ^65^. Our results open a new paradigm by which the RNA m6A writer complex interacts with *de novo* DNA methyltransferases and chromatin repressors during the dynamic establishment phase of epigenetic silencing, ultimately leading to heritable repression of TE expression. Though we capture highly dynamic changes in the proteome at the establishment site of epigenetic silencing, future work can help determine the order of events and direct protein-protein and/or RNA-protein interactions to establish a repressive chromatin environment.

RNA-based surveillance is critical for genome defense in human cells. While small RNAs (e.g. PIWI-interacting RNAs) and the RNA-sensing HUSH complex play key roles in surveying transposons, our study opens intriguing questions that link RNA and DNA modifications in controlling TE levels ^70,71^. We show that m6A reduces levels of HERV-K transcripts during pluripotent cell state transition, suggesting that m6A leads to transcript destabilization or degradation ^61,72^. How RNA is used as a signal for *de novo* DNA methylation remains largely unknown in human cells, although RNA-directed *de novo* DNA methylation is fundamental in plant epigenetic regulation ^73^. Finally, aberrant patterning in DNA methylation, RNA m6A, and transposable element expression is associated with diseases such as cancer, developmental disorders, and neurological diseases. Future work will shed light on how defects in the RNA m6A - *de novo* DNA methylation signaling axis are misregulated in human diseases.

## RESOURCE AVAILABILITY

### Lead contact

Requests should be directed to the lead contact, James K. Nuñez.

### Materials availability

All reagents and resources generated in this study will be made available upon request.

### Data and code availability

The CRISPR screen phenotype scores and proteomics enrichment calculations are available in Tables S1-S5. Custom computational scripts will be made publicly available on GitHub (https://github.com/Nunez-Lab). EM-seq data will be made publicly available on the NCBI SRA (reviewer access link: https://dataview.ncbi.nlm.nih.gov/object/PRJNA1477060?reviewer=c9qftgg45ot3tbd4r5g798kvih

## ACKNOWLEDGMENTS

We thank members of the Nuñez lab for technical assistance and feedback on this work. We thank Norma Neff and the Genomics Platform at Biohub San Francisco and the UCSF CAT (supported by UCSF PBBR, RRP IMIA, and NIH 1S10OD028511-01 grants) for DNA sequencing. We thank Vincent Phan for the bleomycin resistance gene used in this study. The human pluripotent stem cell work was funded by a Hanna Gray Fellowship, a Pew Biomedical Scholars award, and a Shurl and Kay Curci Foundation grant to J.K.N. and the California Institute for Regenerative Medicine EDUC4-12790 grant to N.S.D. and L.G.P. We thank the following funding sources for supporting this work: National Institutes of Health grants R35GM155044 (J.K.N), R35GM142527 (A.S.Y.L.), 5T32GM007232 (N.T.P., A.E.C.), Howard Hughes Medical Institute Hanna Gray Fellowship (J.K.N.), The Shurl and Kay Curci Foundation (J.K.N.), The Pew Charitable Trusts (J.K.N.), Biohub, San Francisco (J.K.N. and S.E.C.), California Institute for Regenerative Medicine EDUC4-12790 (N.S.D. and L.G.P.), University of California, Berkeley Mentored Research Award (N.T.P.), National Science Foundation Graduate Research Fellowship Program (J.P.L.), Weill Neurohub Postdoctoral Fellowship (D.X.), Claudia Adams Barr Program In Cancer Research - Innovative Basic Research Award (S.M.).

## AUTHOR CONTRIBUTIONS

Conceptualization: NS.D., J.K.N.

Methodology: N.S.D., J.K.N., N.T.P., L.G.P., A.E.C., S.M.

Investigation: N.S.D., N.T.P., L.G.P., A.E.C., S.C.U., P.J.C., D.X., B.D.M.

Visualization: N.S.D., N.T.P., L.G.P.

Software: J.P.L., J.L., P.J.C.

Supervision: J.K.N., S.E.C., A.S.Y.L.

Writing – original draft: N.S.D., J.K.N., N.T.P.

Writing – review & editing: N.S.D., J.K.N., N.T.P., A.E.C., L.G.P.

## DECLARATION OF INTERESTS

N.S.D. and J.K.N. have filed a patent application related to the CRISPR technologies resulting from the study. J.K.N. is an inventor of patents related to CRISPRoff/on technologies, filed by The Regents of the University of California. J.K.N. and D.X. have filed a patent application related to the delivery of CRISPR epigenome editors.

## SUPPLEMENTARY INFORMATION

**Table S1.** CRISPR Screen data Humagne Library A (L2FC Score for CRISPRi and CRISPRoff).

**Table S2.** CRISPR Screen data Humagne Library B (L2FC Score for CRISPRi and CRISPRoff).

**Table S3.** Positive hits identified by APEX2-proximity labeling (N-term APEX2-KRAB-dCas9 24 hours).

**Table S4.** Positive hits identified by APEX2-proximity labeling (C-term dCas9-KRAB-APEX2 24 hours).

**Table S5.** Positive hits identified by APEX2-proximity labeling (N-term APEX2-DNMT3A-3L-KRAB).

**Table S6.** List of sgRNA sequences.

**Table S7.** sgRNA and donor sequences for generation of tagged HEK293T cell lines for immunoprecipitation experiments.

**Table S8.** List of antibodies used in this study.

**Table S9.** DNA methylation profiling primers.

**Table S10.** Sequences of Quantitative PCR primers used in this study.

## METHODS

### Plasmids and cloning

All DNA was PCR amplified with KAPA HiFi HotStart ReadyMix PCR Kit (Roche 09420398001). Plasmids were cloned using Gibson assembly (Vazyme C117) and Stellar™ competent cells (Takara 636763) were used for transformation, unless noted. Inducible CRISPRi and inducible CRISPRoff expression plasmids were utilized as previously described ^34^. To clone Cas12-expressing plasmids, mCherry or bleomycin resistance was cloned into the C-terminus of EnAsCas12a lentiviral vector pRDA_174 (Addgene #136476) separated by T2A using Gibson assembly ^37^.

To generate the Cas12 knockout crRNAs, crRNA sequences were cloned from Humagne library A and B for the respective genes. First, crRNA sequences were ordered as complementary sequences from IDT with compatible AgeI and EcoRI overhangs, annealed, and then ligated into PRDA_052 EnAsCas12a sgRNA vector (Addgene # 136474) using AgeI and EcoRI restriction sites ^37^.

To target *CD55* with *S. pyogenes* dCas9 epigenome editors, protospacer sequences were synthesized and ordered as complementary sequences from IDT with BstX1 and BlpI overhangs. For targeting *CD81*, the CD81 single guide (sgRNA)-a protospacer sequence was used ^34^. The protospacer sequences were ligated and cloned into Addgene Plasmid #240637 modified with EGFP selection marker ^63^. For *CLTA* targeting, the sgRNA was expressed using Addgene Plasmid #240637 ^63^. The sgRNA sequences are detailed in table S6.

For MTC chromatin tethering experiments, WTAP and the catalytic domains of METTL3 and METTL14 were amplified from Addgene plasmids 53741 and 155367, respectively, with overhangs to the destination vector backbone ^50,74^. To generate dCas9 fusions, vectors encoding dCas9 only, CRISPRi, and DNMT3A-3L were digested with a restriction enzyme (AgeI for N-terminal fusions and NotI for C-terminal fusion (NEB)) and phosphatase-treated (Quick CIP NEB). Plasmids were assembled using Gibson assembly and transformed into competent cells. Full plasmid sequencing was performed to validate successful construct generation.

For cloning of the APEX2 plasmids for proximity labeling experiments, APEX2 fragments with overlapping ends were amplified from Addgene Plasmid #129640 plasmid by PCR and purified from the gel using Zymogen^TM^ Gel Cleanup Kit ^75^. The purified amplicons were cloned into the N- or C-terminus of dCas9 in a vector encoding KRAB-dCas9-BFP or dCas9-BFP-KRAB under a CAG promoter for transient expression according to manufacturer instructions using the ClonExpress Ultra One Step Cloning Kit (Vazyme, C115). Cloning products were transformed into competent cells, plated on LB Agar plates with carbenicillin resistance, and the resulting plasmids were sequence confirmed.

### Lentivirus generation

Lentiviral particles were produced by transfecting standard packaging vectors into HEK293T along with TransIT^®^ -LT1 Transfection Reagent (Mirus Bio, MIR2306). Cells were seeded at 400,000 cells per well in a 6-well plate and transfected the next day with 1.5 µg lentiviral DNA, 0.1 µg pGag/Pol, 0.1 µg pREV, 0.1 µg pTAT, 0.2 µg pVSVG, 6 µl TransIT^®^ -LT1 and 250 µl Opti-MEM (Gibco) according to the manufacturer’s instructions. The media was changed 24 hours post-transfection with complete DMEM. Viral supernatants were harvested 60 hours post-transfection and filtered using a 0.45 mm PES syringe filter (VWR). The lentiviral particles were stored at -80°C until use.

### Cell culture and cell line generation

HEK293T (ATCC CRL-3216) cells were cultured in DMEM (Gibco 11965092). K-562 (ATCC CCL-243) cells were cultured in RPMI-1640 Medium (Gibco 11875093). Both media were supplemented with 10% (v/v) FBS (VWR) and 100 U/ml streptomycin, 100 mg/ml penicillin, and glutamine (Gibco 10378016). Cell lines were cultured at 37°C with 5% CO_2_ in tissue culture incubators and routinely confirmed to be negative for Mycoplasma.

Human pluripotent stem cells (H9, WA09 Lot #WB68075, WiCell) were cultured in mTeSR Plus (Stem Cell Technologies, 100-0276) and hESC-qualified Matrigel (Corning, 354277)-coated plates following manufacturer’s instructions. Briefly, hESCs-qualified Matrigel was initially pre-diluted in ice-cold DMEM/F12 (Gibco, 10565018) supplemented with 15 mM HEPES (Gibco, 15630080), and plates were incubated at room temperature for an hour before hPSCs were seeded. hPSCs were split every 4-5 days at 70-80% confluency. hPSCs were harvested into cell clumps by incubation for 3 min at 37°C with Gentle Cell Dissociation Reagent (Stem Cell Technologies, 100-0485). Media was refreshed every other day, and cells were cultured at 37°C under normoxia conditions (20% O_2_ and 5% CO_2_).

To generate doxycycline-inducible K-562 cell lines for CRISPR screening, doxycycline-inducible CRISPRi and CRISPRoff v2.1 plasmids were packaged into lentiviral particles ^34^. The harvested lentivirus was transduced by spinoculation into K-562 cells at 1000 xg for 2 hours. Two days post-transduction, cells were treated with 150 ng/ml doxycycline for 72 hours to induce editor expression, followed by fluorescence-activated cell sorting (FACS) to enrich for BFP-positive cells. Following selection, cells were transduced with lentivirus expressing AsCas12f (pRDA_174; Addgene #136476) fused to mCherry via a T2A peptide, together with Cas9 sgRNAs targeting *CD55* or *CD81* ^34,37^. Cells were subsequently sorted for GFP⁺ and mCherry⁺ populations and expanded for downstream CRISPR screening experiments.

To generate HEK293T cell lines with SETDB1 and DNMT3A endogenously tagged with mNeonGreen, HEK293T cells were first engineered to constitutively express mNeonGreen1-10 as described previously ^67,68^. For homology directed repair knockin of mNeonGreen11, the sgRNA and single stranded oligodeoxynucleotide (ssODN) donor sequences were designed using protoSpace.Jam ^76^. Synthetic sgRNA from IDT were resuspended to 260 μM in IDTE buffer (10mM Tris, 0.1mM EDTA, pH 8), combined in equimolar amounts, annealed at 95°C for 5 min, and immediately placed on ice. To assemble ribonucleoprotein (RNP) complexes, 65 pmol of gRNA in 5 µl of Cas9 buffer (20 mM HEPES pH 7.5, 150 mM KCl, 10% v/v glycerol, 1 mM 2-mercaptoethanol) was incubated with Cas9 enzyme (1.25 µl, diluted to a 9 µl final volume in Cas9 buffer) on ice for about 10 min. Following incubation, 1 µl of 50 μM ssODN was added to the RNP mixture. HEK293T cells expressing mNeonGreen10 were harvested, and 20 µl of the cell suspension was mixed with 10 µl of the RNP-ssODN complex for electroporation using a Lonza 4D-Nucleofector (program CM-130). Post-nucleofection, cells were recovered in Dulbecco’s modified Eagle’s medium (DMEM) supplemented with a homology-directed repair (HDR) enhancer. Cells stably expressing mNeonGreen were sorted to purity and the knockin was confirmed by DNA sequencing. The sgRNA and ssODN sequences are detailed in table S7.

### Antibody staining and flow cytometry

Protein expression was assessed by cell surface antibody staining of live cells. Cells were washed with PBS, then incubated in 1:50 diluted antibody for 30 min in the dark at room temperature, washed once with PBS and measured on the BD FACSymphony^TM^ A1 Cell Analyzer (BD Biosciences). For CLTA-GFP HEK293T reporter lines, CLTA-GFP protein expression was assessed directly by flow cytometry. Data analysis was performed using FlowJo^TM^ (v10.10). The antibodies used in this study are listed in table S8.

### CRISPR screens

iCRISPRi and iCRISPRoff K-562 cells were spinoculated with Humagne Library A and B as described below, targeting 20,081 genes each and 1,740 negative control sgRNAs at an multiplicity of infection (MOI) of 0.3-0.4 ^37^. MOI was measured by taking increasing volumes of library lentivirus ranging from 50 µl to 500 µl and performing test spinoculations with 10 x 10^6^ cells in each well of a 6-well plate. Cells were then treated with puromycin for 2 days, and cell survival was compared to control cells that were not treated with puromycin.

For the screen, transductions were performed by spinoculation, wherein 100 x 10^6^ cells were spinoculated with each library. ∼10 x 10^6^ cells were plated in each well of 6-well plates with polybrene and spun at 1000 xg for 2 hours at 37°C with lentivirus expressing guides. Cells were pooled and cultured in T-175 flasks, ensuring their confluency is maintained under 1 x 10^6^/ml and viability greater than 95%. Two days after spinoculation, cells were selected with 2 μg/ml of puromycin for two days and then replenished with 0.8 μg/ml on the third day. Four days after spinoculation, cells were induced with 150 μg/ml of doxycycline for three consecutive days to induce CRISPRi or CRISPRoff editor expression. Seven days after guide infection, all cells were stained with APC CD55 antibody (BioLegend 311312), washed with PBS, and prepared for sorting on the BD Aria Fusion II sorter.

To select for cells that have failed to repress the CD55 reporter, ∼10 x 10^6^ cells that were positive for CD55 protein were sorted to purity (fig. S1F). During sorting, the cells were pre-gated for expressing Cas12 (mCherry), CD55 sgRNA (lentivirus expressing GFP), and CRISPRoff or CRISPRi (BFP). The sorted CD55-positive cells were centrifuged, and the pellet was collected for sample preparation. Additionally, the total unsorted population was collected and prepared for sample preparation to determine the enrichment of failed to silence cells (CD55-positive) compared to the total unsorted population.

For sample preparation to identify the CRISPR RNA (crRNAs), genomic DNA was extracted using NucleoSpin Blood L kit, and crRNAs were amplified using a common P5’ primer Argon and unique P7’ Beaker primer sets for each sample. crRNA libraries were prepared using NEBNext Ultra II DNA Library master mix, analyzed on TapeStation before being sequenced using SE65 on HiSeq4000 ^34^. The resulting fastq files were aligned against the Humagne Library A and B sequences and population-specific enrichment rankings were determined using MAGeCK ^77^.

### Arrayed doxycycline-inducible CRISPRi and CRISPRoff assays in K-562 cells

Doxycycline-inducible CRISPRi and CRISPRoff K-562 cells were seeded in 24-well tissue culture plates and transduced with lentivirus encoding Cas12 crRNAs in the presence of polybrene. Transductions were performed by spinoculation at 1000 xg for 2 hours at 37°C. Two days post-transduction, cells were selected with 2 μg/ml puromycin. Four days after lentiviral transduction, cells were split equally into induced and uninduced conditions. For editor induction, cells were treated with 150 ng/ml doxycycline and 0.8 μg/ml puromycin on the first day, followed by treatment with 150 ng/ml doxycycline alone on the subsequent two days. For uninduced controls, cells were treated with 0.8 μg/ml puromycin on the first day and passaged for 2 days.

Seven days post-lentiviral transduction, cells were stained with the indicated antibodies and analyzed by flow cytometry. Failure-to-silence scores were calculated by dividing the median fluorescence intensity (MFI) of editor-induced (BFP⁺) cells by the MFI of uninduced (BFP⁻) cells for each knockout condition.

### Target locus DNA methylation profiling

Genomic DNA was isolated from HEK293T cells using isopropanol precipitation. For *CD55* profiling, DNA was digested with methylation-insensitive restriction enzymes that cut outside the amplicon site to enrich for the site of interest. Digested DNA was purified using the Qiagen PCR purification kit (Qiagen 28106). Enzymatic methyl conversion was performed on 200 ng of DNA using the NEB Enzymatic Methyl Conversion Module (NEB E7125S) according to the manufacturer’s instructions. Following conversion, half of the converted DNA was PCR amplified using primers designed with MethPrimer. PCR was performed using the Keypo Se polymerase mix (PK512) for 35-37 cycles. Amplified DNA was purified using the Qiagen PCR purification kit (Qiagen 28106). DNA was prepared using the rapid barcoding system for Nanopore sequencing. DNA was sequenced on a Nanopore sequencer by the Barker DNA Sequencing Facility. Specific primer sequences for DNA methylation profiling can be found in table S9.

Methylation was called using a script created with the help of the Honeybee programming system ^78^. FastQ files generated from sequencing were aligned to a C-to-T converted reference genome using Minimap2 (version 2.26-r1175) with the arguments -ax map-ont --sam-hit-only ^79^. Alignment files were converted to bam files, sorted, and indexed using SAMtools ^80^. Methylation calling from the resulting files was performed using a custom Python script (LEMONmC.py, available at https://github.com/Nunez-Lab/LEMONmethyl-seq/tree/main) using pysam version 0.24.0 ^80,81^. For all loci, only CpG sites annotated from the reference were considered. Percent methylation at each CpG site was plotted using Prism (GraphPad).

### APEX2 proteomic labeling

HEK293T cells with stable expression of sgRNAs targeting the endogenous *CLTA-GFP* promoter were cultured in 15 cm dish and transfected with plasmids expressing dCas9-KRAB-APEX2, APEX2-KRAB-dCas9, dCas9-APEX2 or APEX2-dCas9 only ^34^. 24 hours or 48 hours post-transfection, cells were treated with 500 µM biotin-tyramide (MilliporeSigma -SML2135) for 30 min. 1 mM final H_2_O_2_ was added to each dish for a total labeling time of 1 minute. Cells were quenched with 2X quenching buffer (10 mM Trolox and 20 mM sodium ascorbate in DPBS), collected and washed twice with cold DPBS. Cell pellets were lysed with 500 µl cytoplasmic lysis buffer (10 mM Tris pH 8, 1.5 mM MgCl_2_, 10 mM KCl, 0.2% NP-40, and 1 mM DTT) for 10 min on ice. After centrifugation at 2500 xg for 5 min at 4°C, the pellets were collected and washed once with cold PBS. Cell pellets were lysed in 200 µl nucleus lysis buffer (20 mM Tris pH 8, 25% glycerol, 400 mM NaCl, 1.5 mM MgCl_2_, 10 mM EDTA, 1 mM DTT, and 0.2 µl benzonase (Sigma-Aldrich, E1014) and incubated on ice for 30 min. After centrifugation at 20,000 xg for 10 min at 4°C, the supernatant was collected, combined with 800 µl of cold RIPA lysis buffer (10% glycerol, 25 mM Tris-HCl pH 8, 150 mM NaCl, 2 mM EDTA, 1% NP-40, 0.2% sodium deoxycholate, 5 mM Trolox, 10 mM sodium ascorbate, and 1x Protease Inhibitor), and incubated with Streptavidin beads (Pierce) at RT for 1 hour. Beads were washed twice with RIPA lysis buffer, once with 1 M KCl solution, once with 2 M Urea, pH 8, and twice with RIPA lysis buffer (without detergent or quenching reagents). Beads were washed three times with ABC buffer (50 mM ammonium bicarbonate, pH 8), rotated head-over-head at 4°C for 20 min. Beads were resuspended in 25 µl ABC buffer with 2.5 µl of trypsin (1 µg/µl) and incubated for 5 hours at 37°C. Peptides were collected, quantified with Fluorescent Peptide Analysis, and 0.6 µg of each sample was subjected to mass-spectrometry (MS)-based analysis. DIA data was analyzed using Spectronaut 18.4 (Biognosys Schlieren, Switzerland) using the direct DIA workflow with PTM localization selected. We identified 58 hits showing the most significant enrichment in dCas9-KRAB-APEX2 at 24 hours, 423 hits showing the most significant enrichment in APEX2-KRAB-dCas9 at 24 hours, and 49 hits showing the most significant enrichment in APEX2-KRAB-dCas9 at 48 hours (Table S3, S4, and S5).

### Human naive hPSC resetting and repriming from naive pluripotency

Primed hPSCs were cultured at least two passages in mTeSR Plus media before starting the transition. hPSCs were reset to naive pluripotency following the protocol described previously ^82^. In detail, a day before starting the conversion, mitomycin C-inactivated mouse embryonic fibroblast (iMEFs) were seeded in a 0.1% Gelatin (Stem Cell Technologies, 07903)-coated 6-well plates at a density of 1 x 10^5^ iMEFs/cm^2^. A day later, primed hPSCs were harvested in single-cell suspension by using Accutase (Gibco, A1110501), and 2 x 10^5^ hPSCs were seeded in each well with mTeSR Plus supplemented with 10 μM Y-27632 inhibitor (MedChemExpress, HY-10071). Cells were maintained under normoxia conditions (20% O_2_ and 5% CO_2_) during the first 24 hours. The next day, the media was switched to PXGGA media and cells were cultured under hypoxia (5% O_2_ and 5% CO_2_) from that day onwards.

PXGGA media is composed of N2B27 media (50% DMEM/F12, 50% Neurobasal (Gibco, 21103149), 1X N2 supplement (Gibco, 17502048), 1X B27 supplement (Gibco, 17504044), 1X MEM NEEAA (Gibco, 11140050), 1X GlutaMAX (Gibco, 35050061), 100 units/ml penicillin/100 μg/ml Streptomycin (Gibco, 15140122), 0.0053% BSA Fraction V (Gibco, 15260037), 0.1 mM β-Mercaptoethanol (Gibco, 21985023)) supplemented with 1 μM PD0325901 (SelleckChem, S1036), 2 μM XAV939 (Sigma-Aldrich, X3004), 2 μM Gö6983 (Tocris, 2285), 2.5 μM GDC-0994 (SelleckChem, S7554) and 10 ng/ml Activin A (Gibco, 120-14P). PXGGA media was changed every other day, and cells were split in a 1:6 dilution for the first time after 12 days by incubating them at 37°C with Accutase for 3 min. After the first passage, naive hPSCs were split every 5-6 days, changing the PXGGA media every other day and supplementing it with 10 μM Y-27632 during the first 24 hours. Experiments were performed in naive hPSCs cultured for at least 5 passages in PXGGA media, and the quality of naive conversion was quantified by flow cytometry with the loss of CD57 (primed pluripotency) and the acquisition of CD7, SUSD2 and CD75 ^83^.

Approximately 5 x 10^5^ naive hPSCs were seeded per each Matrigel-coated 9.6 cm^2^ well (6-well plate) in mTeSR Plus media supplemented with 10 μM Y-27632 during the first 24 hours. The next day, Y-27632 was removed from the media and mTeSR Plus was refreshed every other day. For m6A inhibition experiments, naive hPSCs were seeded during the first 24 hours without inhibitor. One day later, either DMSO, 25 μM, 50 μM, or 100 μM m6A inhibitor (STM2457, SML3360 Sigma-Aldrich) was added into the media, refreshing the media every day until day 4. Day 4 repriming cells were dissociated and collected for further analysis by using Accutase as previously described.

### Generation and delivery of RENDER CRISPRoff

CRISPRoff enveloped virus-like particles (eVLPs) were generated as previously described ^63^. Briefly, approximately 4 x 10^6^ Lenti-X 293T cells were seeded in 10-cm BioLite cell culture dishes (Thermo Fisher Scientific, USA). About 24 hours after seeding, cells were transfected with plasmids encoding 0.5 μg VSV-G, 0.45 μg gag-pol, 4.05 μg gag-CRISPRoff (KOX1), and 6.0 μg mU6-CLTA sgRNA using TransIT-LT1 transfection reagent (Mirus Bio) diluted in Opti-MEM (Gibco), according to the manufacturer’s instructions.

At day 3 post-transfection, culture supernatants were collected and filtered through a 0.45-μm PES syringe filter (VWR). eVLP particles were precipitated overnight using Lenti-X Concentrator (Takara Bio) following the manufacturer’s protocol. Concentrated eVLPs were resuspended in 100 µl Opti-MEM to achieve a 100-fold concentration, frozen overnight using a Corning CoolCell LX freezing container, and stored at −80°C. Frozen eVLPs were thawed on ice prior to use.

To generate Cas12 expressing CLTA lines, we used a previously engineered HEK293T cell line with a CLTA-targeting sgRNA stably expressed ^34^. The cells were transduced with lentivirus encoding EnAsCas12a with bleomycin cassette and selected with 500 μg/ml zeocin (Thermo Fisher Scientific, USA, R25001) for 10 days. The Cas12 expressing cells were subsequently transduced with lentiviruses expressing Cas12 crRNAs targeting CD55 (non-targeting control), METTL3, VIRMA, WTAP, or SETDB1. About 48 hours after transduction, cells were selected with 2 μg/ml puromycin for another 48 hours. Following puromycin selection, cells were plated in 96-well flat-bottom tissue culture plates at a density of 15,000 cells per well. About 12 hours after plating, eVLPs encapsulating the indicated epigenome editors were added to each well. Cells were cultured for an additional 3 days post-eVLP treatment, after which they were harvested and processed for DNA methylation analysis.

### DNA methylation maintenance assay

To perform the CRISPRoff maintenance screen, CLTA-GFP HEK293T cells previously silenced by CRISPRoff RENDER, were sorted by fluorescence-activated cell sorting (FACS) to isolate GFP-negative cells and subsequently expanded. Cells were then transduced with lentivirus expressing Cas12 with a bleomycin resistance cassette. Two days post-transduction, cells were selected for Cas12 expression using 500 μg/ml zeocin (Thermo Fisher Scientific, R25001) for 10 days.

Following Cas12 selection, cells were plated in 24-well tissue culture plates and transduced with lentiviruses expressing Cas12 crRNAs targeting RNA m6A regulatory factors (one gene knockout per well in an arrayed format). Two days post-transduction, cells were selected with 2 μg/ml puromycin. Cells were passaged every two days, and GFP expression was monitored on days 6 and 8 post-guide delivery by flow cytometry using a BD FACSymphony A1 Cell Analyzer (BD Biosciences). Data analysis was performed using FlowJo^TM^ (v10.10).

### MTC chromatin tethering experiments

Epigenome editors were transfected using previously established methods. Two HEK293T cell lines were utilized: one encoding *CLTA* endogenously tagged with GFP along with a transgene expressing a sgRNA targeting the *CLTA* promoter, and HEK293Twith a sgRNA expressed targeting the *CD55* promoter.

In short, the HEK293T cell lines were plated in 24-well plates at a seeding density of ∼70,000 per well. A day after plating, cells were transfected with 500 ng of plasmid DNA per well with Mirus TransIT® -LT1 transfection reagent according to the manufacturer’s specifications. To test the editors, four wells of a 24-well plate were transfected with plasmid DNA. Two days following transfection, cells were sorted for BFP⁺ cells that indicate successful delivery of the epigenome editor. Following sorting, cells were plated into three wells of a 96-well plate for technical replicates. CLTA expression was assayed using GFP expression, and CD55 protein expression was assayed using cell-surface staining and flow cytometry to evaluate target knockdown.

For DNA methylation profiling experiments, the same transfection as described above was performed, however BFP⁺ cells were sorted on day 3 post-transfection either and immediately harvested for gDNA extraction and DNA methylation analysis or cultured for 12 days before DNA methylation profiling.

### Immunoprecipitation assay on cell lysates

One 6 well plate of HEK293T cells with SETDB1-mNeonGreen tagged or mNeonGreen-DNMT3A tagged were harvested, washed with ice-cold PBS, and lysed in three volumes of NP-40 lysis buffer (50 mM HEPES-KOH pH 7.5, 150 mM KCl, 5 mM MgOAc, 0.5% Nonidet P-40 alternative, and 0.5 mM DTT supplemented with 1x protease inhibitor) for 10 min. After centrifuging at 4°C at 21,000 xg for 10 min, the total proteins in the supernatant were collected. 50 μg of the protein lysate was used as input, and the remainder was incubated with mNeonGreen-Trap magnetic agarose (ChromoTek) overnight at 4°C. mNeonGreen-associated proteins were washed twice in wash buffer (50 mM HEPES-KOH pH 7.5, 200 mM KCl, 5 mM MgOAc_2_, 1% Nonidet P-40 alternative, and 0.5 mM DTT) and subjected to western blot analysis.

### Western blot

Protein samples were mixed with 1x loading buffer (containing SDS and β-mercaptoethanol) and heated at 95°C for 5 min. The denatured proteins were separated by size in 4-12% Bis-Tris Protein Gels (Thermo Fisher Scientific, USA NW04122BOX) and transferred to PVDF membranes (Bio-Rad). Membranes were blocked with 5% (w/v) non-fat dried milk (VWR) in TBST (0.05% Tween-20) for 1 hour at room temperature and incubated overnight with primary antibody in 1% milk in TBST (0.05% Tween-20) at 4°C. Membranes were washed three times, 5 min each, with TBST and incubated with a 1:5000 dilution of goat anti-Mouse IgG or goat anti-Rabbit IgG secondary antibody (Li-Cor 926-68070 or 926-32211) for 1 hour. Membranes were washed three times with TBST, and the protein bands on the membranes were detected and analyzed using the Image Studio^TM^ Software from the LICORbio Odyssey CLx Imaging System. The following primary antibodies were used: β-actin (Cell Signaling, 3700S), GAPDH (Abcam, Ab245355), SETDB1 (Proteintech, BC009362), DNMT3A (Proteintech, 20954-1-AP), KAP1 (Cell Signaling, Q13263), H3K9me3 (Abcam, Ab8898), VIRMA (Proteintech, 25712-1-AP), WTAP (Proteintech, 60188-1-Ig), METTL3 (Proteintech, 15072-1-AP), YTHDF2 (Proteintech, 24744-1-AP), YTHDC1 (Proteintech, 29441-1-AP). A list of antibodies is detailed in table S8.

### RNA m6A immunoprecipitation

One 6 well plate of hPSCs in each condition (naive, Day 2 reprimed, Day 4 reprimed untreated, Day 4 reprimed treated with DMSO and Day 4 reprimed treated with 50µM STM2457 for 72 hours) were lysed in three volumes of NP-40 lysis buffer (50 mM HEPES-KOH pH 7.5, 150 mM KCl, 5 mM MgOAc, 0.5% Nonidet P-40 alternative, 0.5 mM DTT). Total RNA was extracted from the lysate using phenol-chloroform extraction with acid phenol:chloroform:isoamyl alcohol (25:24:1, pH 5.2, Fisher Scientific) and ethanol precipitation. The total RNA samples were precipitated overnight at −20°C, then centrifuged for 30 min at 21,000 xg at 4°C. RNA pellets were washed with 70% ethanol, air-dried for 2 min, and resuspended in nuclease-free water. One microgram of total RNA was treated with RQ1 DNase (Promega) for 30 min at 37°C according to the manufacturer’s instructions. The total RNA (DNA-free) was purified with standard phenol:chloroform extraction and ethanol precipitation. The total RNA samples were precipitated overnight at −20°C, then centrifuged for 30 min at 21000 xg at 4°C. RNA pellets were washed with 70% ethanol, air-dried for 2 min, and resuspended in 12 µl nuclease-free water. rRNAs were depleted from each RNA sample using the NEBNext rRNA Depletion Kit (NEB). Ten percent of the rRNA-depleted RNA was collected and proceeded to RT-qPCR as the input. The rest of the rRNA-depleted RNA was used to perform m6A immunoprecipitation (NEB) according to the manufacturer’s instructions using Pierce^TM^ Protein A/G magnetic beads (Thermo Fisher Scientific, USA). m6A RNAs were isolated from the beads using phenol:chloroform and ethanol precipitation. The purified m6A RNA was subjected to RT-qPCR.

### Reverse transcription and quantitative PCR (RT-qPCR)

cDNA was generated from the input and immunoprecipitated RNA samples from m6A immunoprecipitation experiments by reverse transcription using random hexamers and the SuperScript^TM^ II Reverse Transcriptase kit (Thermo Fisher Scientific, USA) according to the manufacturer’s instructions. Quantitative PCR was performed using iTaq^TM^ Universal SYBR^R^ Green Supermix (Bio-Rad) on the CFX Opus 96 Real-time PCR system. Gene expression levels of HERV-K, LINE1, ATF4, and GAPDH were normalized to ACTB using the 2-ΔΔCq method. Primer sequences are listed in table S10.

### Chromatin immunoprecipitation quantitative polymerase chain reaction

About 5 x 10^6^ cells of each condition were collected in a tube and crosslinked in 1% formaldehyde in DMEM for 10 min at room temperature, then quenched with 200 mM glycine for 10 min at room temperature. Cells were pelleted by spinning down at 500 xg for 5 min at room temperature. The media was aspirated, and cells were resuspended in 10 ml cold PBS with 1% Protease Inhibitors. The cells were spun down again at 500 xg for 5 min at 4°C, media was aspirated and resuspended in 1 ml cold PBS with 1% Protease inhibitor. Cells were then transferred to a microcentrifuge tube, pelleted at 300 xg for 5 min at 4°C. Cells were lysed in 1 ml lysis buffer (5 mM PIPES pH 8.0, 85 mM KCl, 1% Igepal, and 1X Protease Inhibitor) on ice for 10 min. Samples were spun down at 2000 rpm at 4°C for 5 min, the supernatant was discarded, and nuclei were lysed in 300 µl of nuclei lysis buffer (50 mM Tris pH 8.0, 10 mM EDTA, 1% SDS, 1X protease inhibitor) for 10 min on ice. The nuclei were transferred into Bioruptor® Pico Microtubes 1.5 ml Diagenode Bioruptor tubes in a Diagenode Bioruptor® Pico sonication device and sonicated (10 cycles 30 sec on 30 sec off). The samples were spun down at 13,000 rpm at 4°C for 10 min. The supernatant was collected and diluted to 1.5 ml with IP dilution buffer (50 mM Tris pH 7.5, 150 mM NaCl, 1 mM EDTA, 1% Igepal, 0.25% Na-DOC, 1X protease inhibitor). About 4% of the sample was saved for input, and 5 μg of H3K9me3 antibody (Active Motif, AB_2687870) was added to the sample and rotated overnight at 4°C. Pierce™ Protein A/G Magnetic Beads (Thermo Fisher Scientific, USA) were added to the sample and cleaned using a magnetic rack. For cleaning, the samples were washed with 1 ml of the following: one round of ice-cold IP wash buffer 1 (20 mM Tris pH 8.0, 2 mM EDTA, 50 mM KCl, 1% Triton X-100, 0.1% SDS), followed by two rounds of ice-cold high salt buffer (20 mM Tris pH 8.0, 2 mM EDTA, 500 mM NaCl, 1% Triton X-100, 0.01% SDS), one round of ice-cold IP wash buffer 2 (10 mM Tris pH 8.0, 1 mM EDTA, 0.25 M LiCl, 1% Igepal, 1% Na-DOC), and finally with two rounds of ice-cold TE (10 mM Tris pH 8.0, 1 mM EDTA) buffer. After washing, samples were eluted using 100 µl of elution buffer (50 mM NaHCO_3_, 1% SDS), by shaking the samples for 1 hour at 65°C on a thermomixer.

Samples and input were reverse crosslinked by adding 0.3 M NaCl, then treated with 1µl of RNaseA at 37°C for 1 hour, and 2.25 µl proteinase K at 63°C overnight. DNA was cleaned using Zymo Clean and Concentrator Kit. Quantitative PCR was performed using iTaq^TM^ Universal SYBR^R^ Green Supermix (Bio-Rad) on the CFX Opus 96 Real-time PCR system. Sequences for ChIP-qPCR primers are listed in table S10.

### Whole genome DNA methylation sequencing and analysis

Whole genome DNA methylation profiling was performed as described previously ^63^. Twenty-four libraries were sequenced with paired-end 150 bp reads on an Illumina NovaSeq X 25B 8 Lanes (25 billion clusters), fifteen of which were used for analysis (three biological replicates each of naive hPSCs, reprimed hPSCs, DMSO-treated reprimed hPSCs, and 25 and 50 µM STM2457 RNA m6A inhibitor-treated reprimed hPSCs). We obtained between 804.2 million and 1456.1 million reads per library across the fifteen libraries, resulting in median CpG coverage between 11x and 19x.

Reads were preprocessed, and methylation was called using a script created with the help of the Honeybee programming system ^78^ using steps as previously described ^63^ but with no locus filtering and with the addition of a read deduplication step (deduplicate_bismark --bam -- multiple). Quality control was assessed via FastQC version 0.12.1 ^84^ with MultiQC version 1.18 ^85^, adapter trimming was performed with cutadapt version 4.4 ^86^, and methylation calling was performed with Bismark version 0.24.2 ^87^.

Downstream methylation analysis was performed using a custom pipeline using Python version 3.14 with Matplotlib version 3.10.9 ^88^, NumPy version 2.4.4 ^89^, Polars version 1.40.0, and SciPy version 1.17.1 ^90^. Region annotations for this pipeline were obtained from the UCSC Genome Browser ^91^. Promoters were defined as 500 bp upstream and 100 bp downstream of EPDnew (version 006)-annotated transcription start sites ^92^. CGI annotations were obtained via the table hg38.cpgIslandExt following the traditional definition ^93^. Gene bodies were defined as spanning the first transcription start site to the last transcription end site as obtained via the table hg38.ccdsGene using data from the Consensus Coding Sequence (CCDS) project ^94–97^. Transposable elements annotations were obtained via the table hg38.rmsk using data from RepeatMasker ^98^. Percent attribution of the total change in pooled methylation *Δβ* to a region *R* as defined above was calculated as the change in the region’s pooled *β* value weighted by the proportion of read counts in the region divided by the total change in pooled *β* value, *i.e.*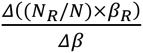, where *N* is the number of reads for the region *R*, *N* is the total number of reads, *β* is the pooled beta value for the region *R*, and *β* is the overall pooled *β* value. *p*-values for determining regions with *de novo* methylation were calculated via an unpaired *t*-test on pooled *β* values with three biological replicates in each of the naive and untreated reprimed conditions. *p*-values and slope values for the m6A inhibitor dose dependence analysis were calculated using linear regression on pooled *β* values with three biological replicates each between the DMSO-treated, 25 µM inhibitor-treated, and 50 µM inhibitor-treated conditions. All adjusted *p*-values were calculated using the Benjamini-Hochberg procedure ^99^.

## SUPPLEMENTARY FIGURES

**Figure S1.**
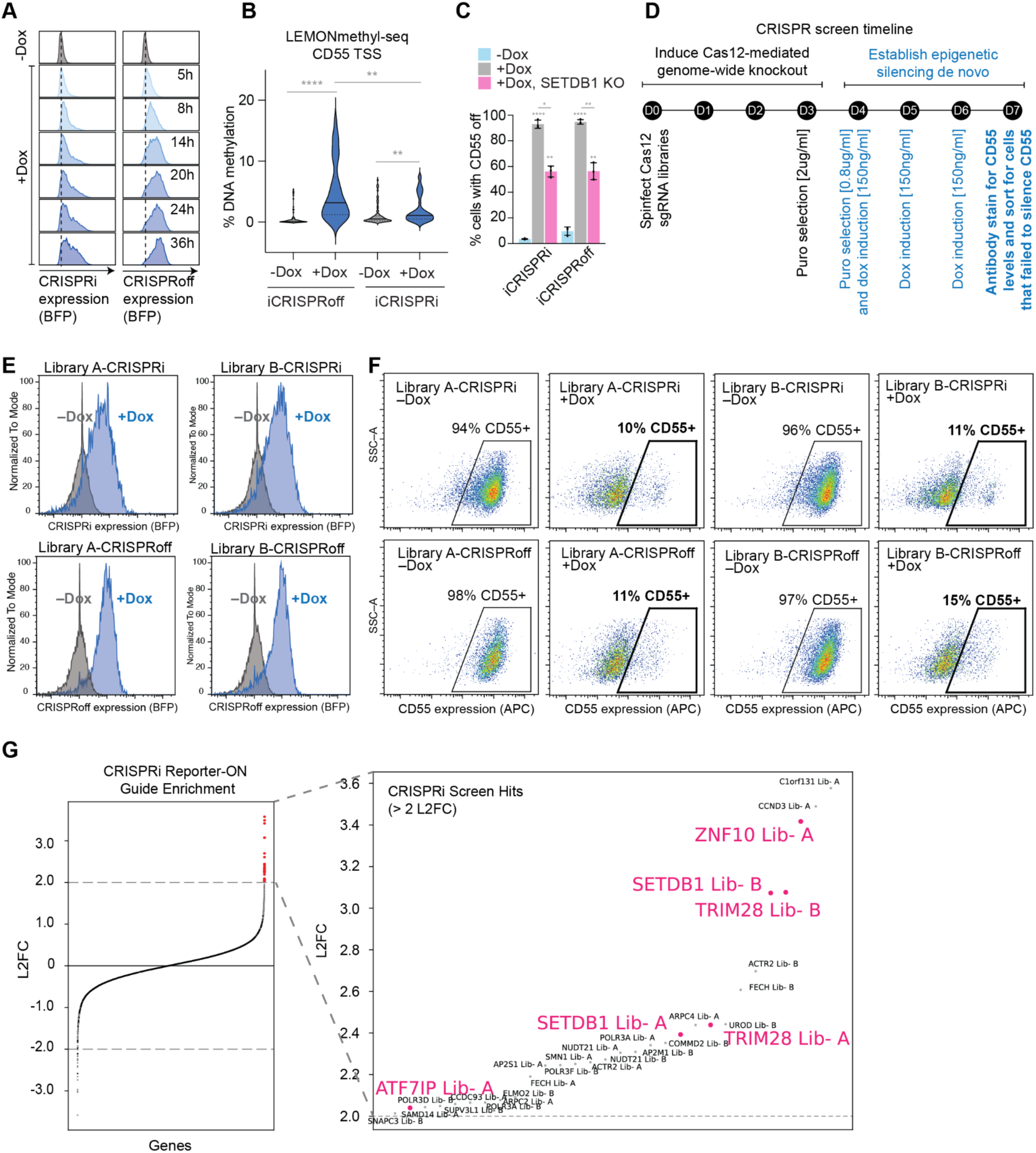
Genome-wide CRISPRoff screens identify regulators of heritable epigenetic silencing. (A) Time course of BFP expression in CRISPRi and CRISPRoff doxycycline-induced K-562 cells. (B) Violin plot showing average methylation gain at CD55 TSS in the doxycycline-induced iCRISPRi and iCRISPRoff cell lines. \**p*<0.05, ns, not-significant, Mann-Whitney U test. (C) Bar plot showing average percent of CD55 silenced cells in doxycycline induced iCRISPRi and iCRISPRoff cells, with SETDB1 knockout. \**p*<0.05, \*\**p*< 0.01, \*\*\*\**p*< 0.0001, unpaired t-test. n = 3 biological replicates, error bars ± 1 s.d. (D) Genome-wide CRISPR knockout screen timeline to identify factors required to establish *de novo* DNA methylation and heritable silencing. (E) Flow histogram showing BFP expression 72 hours post doxycycline induction in iCRISPRi and iCRISPRoff after Humagne Library A and B knockouts. (F) Single-cell flow plot shows total unsorted and sorted (gated) populations for each Humage Library knockout and editor-induced conditions. (G) Log2 fold-change (L2FC) ranking of sgRNAs enriched in the CRISPRi failure-to-silence population relative to the unsorted population. sgRNAs are colored by functional pathways, with KRAB-associated factors in pink.

**Figure S2.**
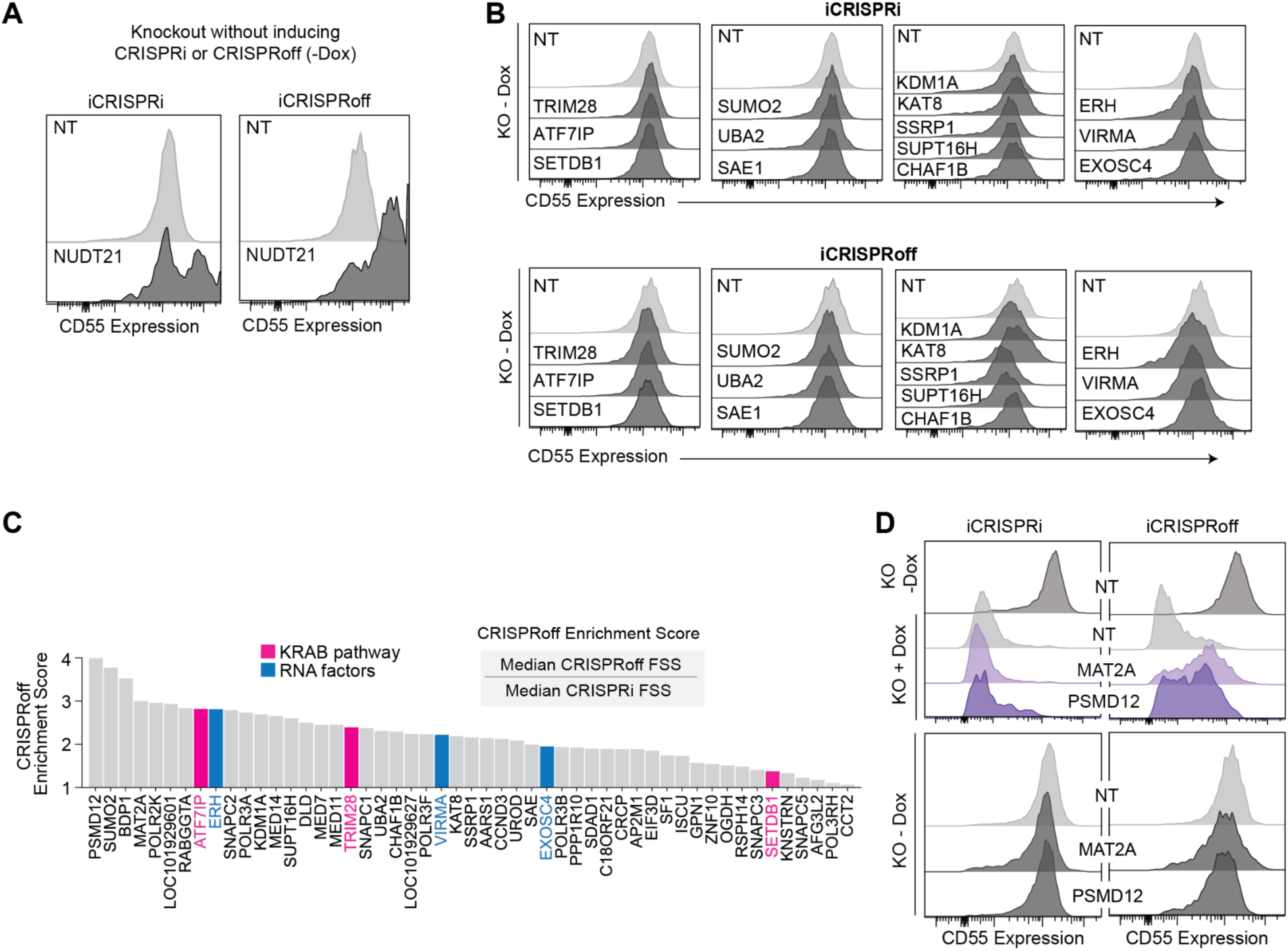
Validation of CRISPR screen hits identifies RNA m6A factors are enriched in CRISPRoff. (A) Flow histogram showing CD55 expression after *Nudt21* KO in the absence of doxycycline in iCRISPRi and iCRISPRoff lines. (B) Flow histogram showing CD55 expression after knockout of validated factors in the absence of doxycycline in iCRISPRi and iCRISPRoff lines. (C) CRISPRoff enrichment scores of top iCRISPRoff and iCRISPRi hits as calculated from the arrayed validation screen. (D) Flow histogram showing CD55 expression after knockout of *Mat2a* and *Psmd12* in iCRISPRi and iCRISPRoff in the absence and presence of doxycycline.

**Figure S3.**
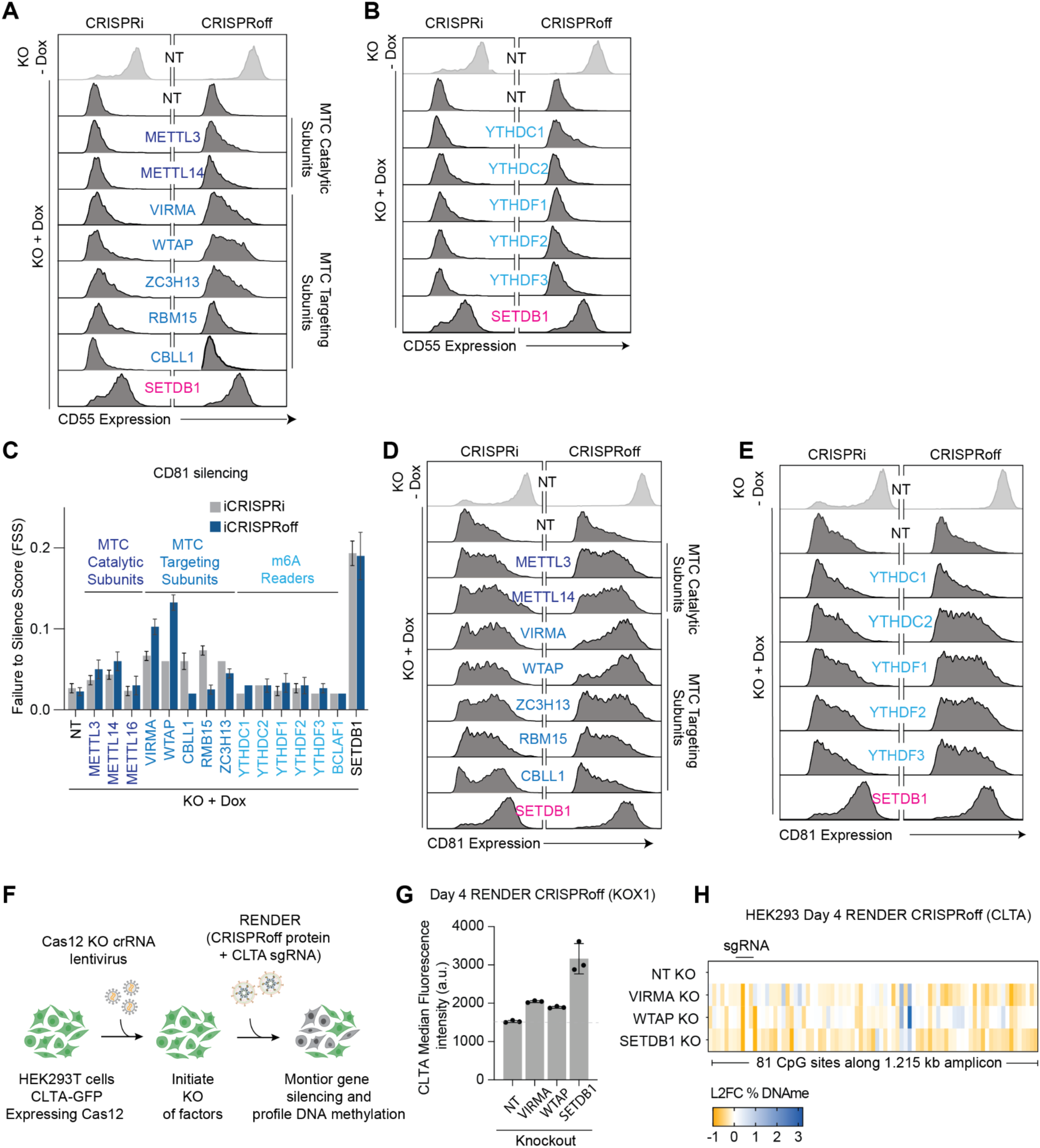
RNA m6A dependency is consistent across different gene targets and cell types. (**A** and **B**) Flow histograms showing CD55 protein levels upon induction of CRISPRi and CRISPRoff after knockout of m6A writer complex components (**A**), readers (**B**) and positive control SETDB1 (**C**) Comparison of FSS values for CD81 following knockout of MTC targeting components and RNA m6A reader proteins in CRISPRi and CRISPRoff secondary validation screens. n = 4 biological replicates, error bars ± 1 s.d. (**D** and **E**) Flow histograms showing CD81 protein levels upon induction of CRISPRi and CRISPRoff after knockout of m6A writer complex components (**D**), readers (**E**) and positive control SETDB1. (**F**) Schematic of the RENDER protein-based delivery assay used to evaluate target gene repression in CRISPR-Cas9 generated knockout (KO) HEK cell lines. (**G**) CLTA silencing measured by CLTA median fluorescence intensity after RENDER CRISPRoff delivery upon knockout of VIRMA and WTAP m6A-writer factors and SETDB1. n = 3 biological replicates, error bars ± 1 s.d. (**H**) DNA methylation profiling showing gain or loss in DNA methylation establishment in VIRMA, WTAP and SETDB1 knockout conditions using RENDER CRISPRoff, relative to non-targeting control.

**Figure S4.**
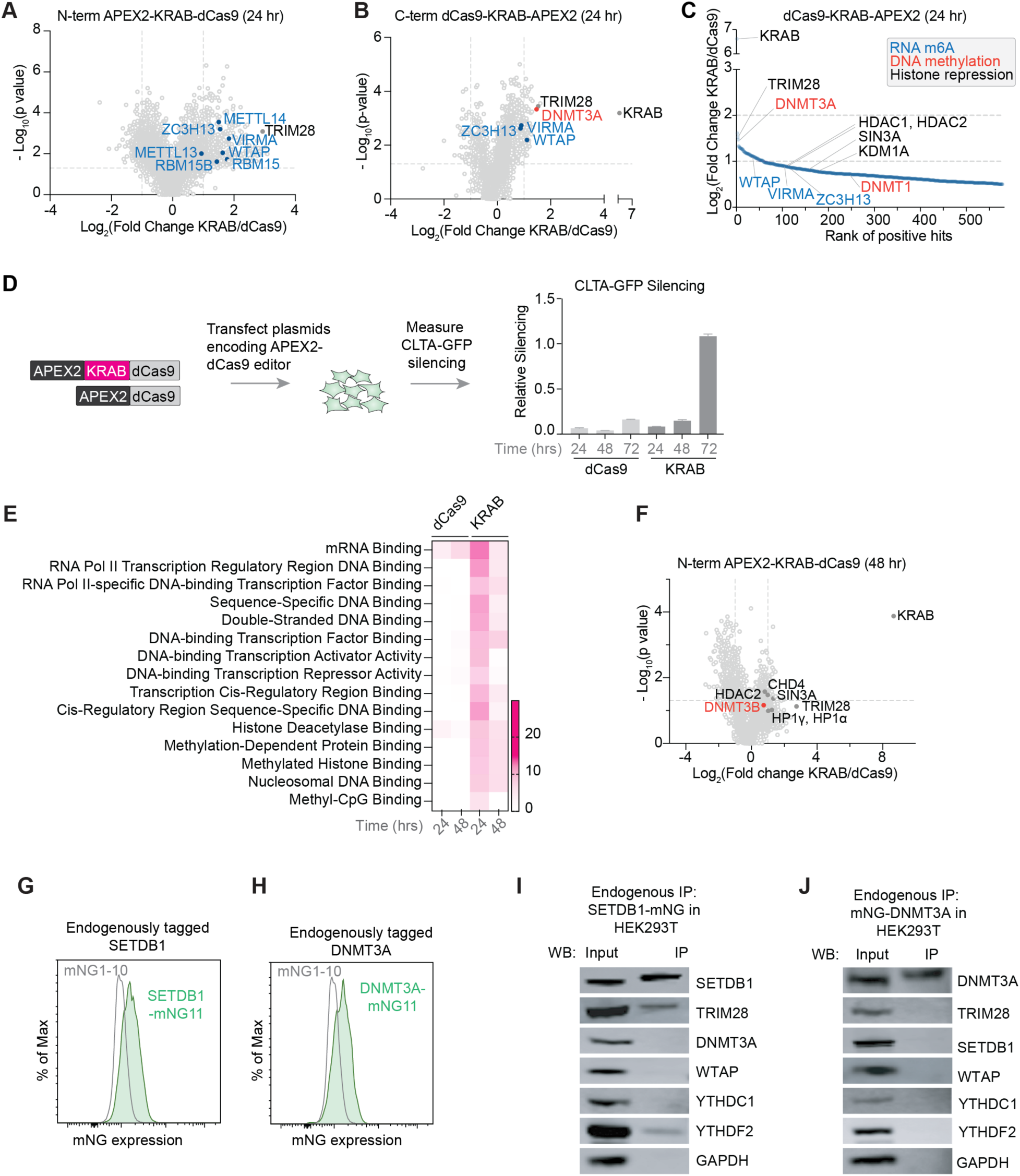
Proteomics identifies co-localization of RNA m6A factors with chromatin repressors. (**A** and **B**) Volcano plot of KRAB interactors measured at 24 hours post-transfection of plasmids encoding (**A**) APEX2-KRAB-dCas9, and (**B**) dCas9-KRAB-APEX2 (n = 2 biological replicates). Proteins in the RNA m6A pathway are highlighted in blue, orange for DNA methylation proteins, and black for histone repression. (C) Rank-based plot of KRAB interactors measured at 24 hours post-transfection of plasmids encoding dCas9-KRAB-APEX2 (n = 2 biological replicates). Proteins in the RNA m6A pathway are highlighted in blue, including members of the RNA m6A writer complex, pink for DNA methylation proteins, and black for histone repression. (D) Time course analysis of CLTA-GFP silencing after transfecting APEX2-dCas9 alone and APEX2-KRAB-dCas9 at 24, 48, and 72 hours post-transfection. (E) GO term analysis of proteins detected from APEX2 labeling at 24 hours and 48 hours post-delivery of APEX2-dCas9 alone and APEX2-KRAB-dCas9. (F) Volcano plot of KRAB interactors measured at 48 hours post-transfection of plasmids encoding APEX2-KRAB-dCas9. Proteins in the KRAB-TRIM28 pathway are highlighted in black, orange for DNA methylation proteins. (**G** and **H**) Flow histograms showing expression of (**G**) SETDB1 and (**H**) DNMT3A tagged endogenously with mNeonGreen in HEK293T cells. (**I** and **J**) Immunoprecipitation of (**I**) SETDB1 and (**J**) DNMT3A tagged endogenously with mNeonGreen in HEK293T cells. Western blots are shown for SETDB1, DNMT3A, TRIM28, WTAP, YTHDC1, YTHDF2, and negative control GAPDH. Representative of n = 3 biological replicates.

**Figure S5.**
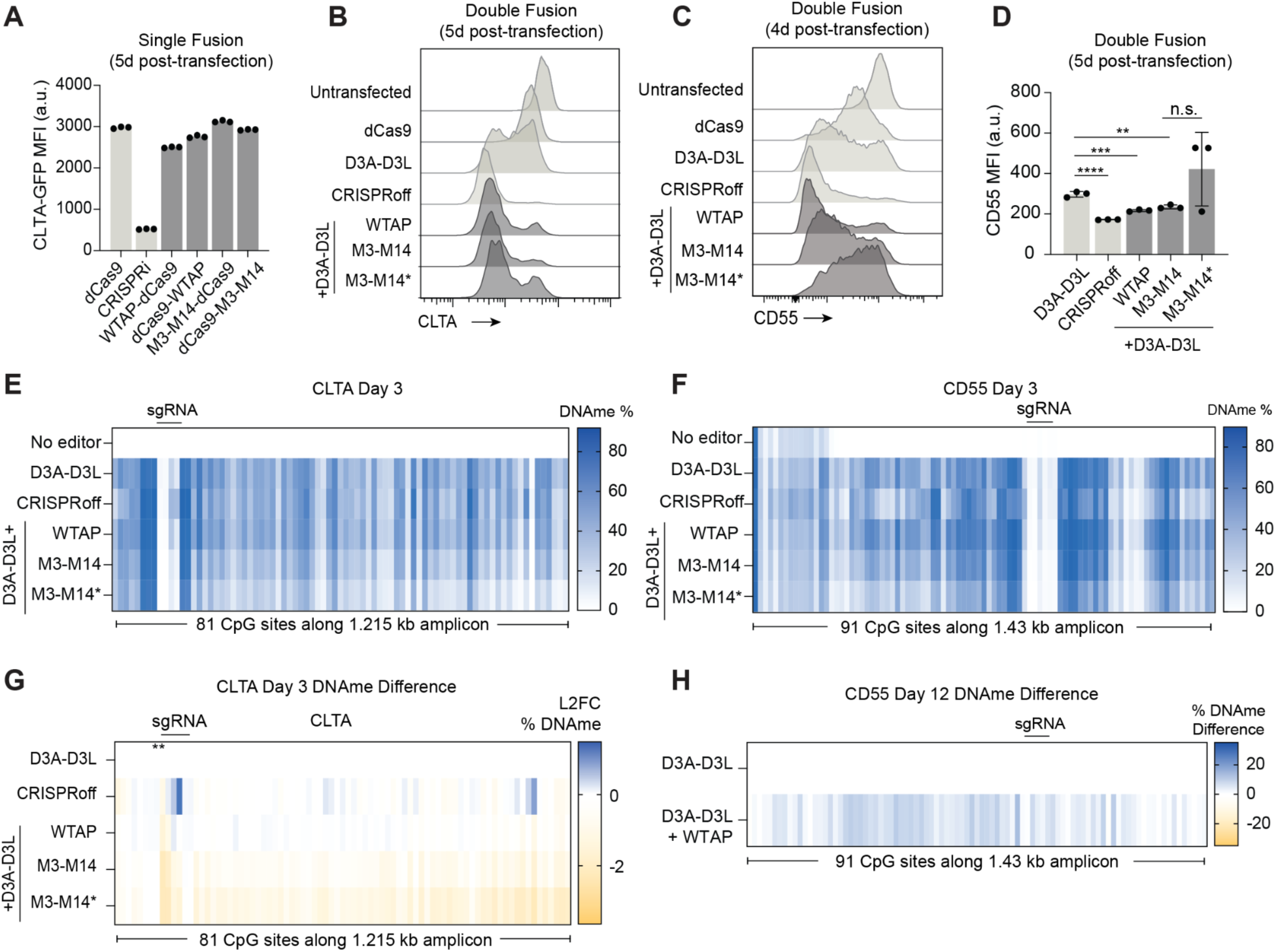
Targeted co-recruitment of MTC components and DNMT3A/L synergistically enhances *de novo* DNA methylation. (**A**) Bar plot showing median fluorescence intensity (MFI) of CLTA-GFP after transfection with single fusion dCas9 constructs. (**B** and **C**) Flow histogram showing silencing of CLTA-GFP (**B**) and CD55 (**C**) protein levels in cells transfected with double fusion constructs. (**D**) Bar plot showing median fluorescence intensity (MFI) of CD55 after transfection with double fusion constructs. n.s. non-significant, \*\**p*< 0.01, \*\*\**p*< 0.001, \*\*\*\**p*< 0.0001, unpaired t-test. n = 3 biological replicates, error bars ± 1 s.d. (**E** and **F**) DNA methylation profiling of CLTA promoter (**E**) and CD55 promoter (**F**) 3 days post-transfection with D3A-D3L-dCas9, CRISPRoff, D3A-dCas9-WTAP, D3A-dCas9-METTL3/14 (M3-M14), and D3A-dCas9-METTL3/14* (M3-M14*) catalytic mutant. (G) DNA methylation profiling showing differences in DNA methylation established by the MTC dual fusion constructs relative to D3A-D3L-dCas9 only at the *CLTA* locus. (H) DNA methylation profiling showing differences in DNA methylation established by the MTC dual fusion construct D3A-D3L-dCas9-WTAP, relative to D3A-D3L-dCas9 only at the *CD55* locus measured at 12 days post transfection.

**Figure S6.**
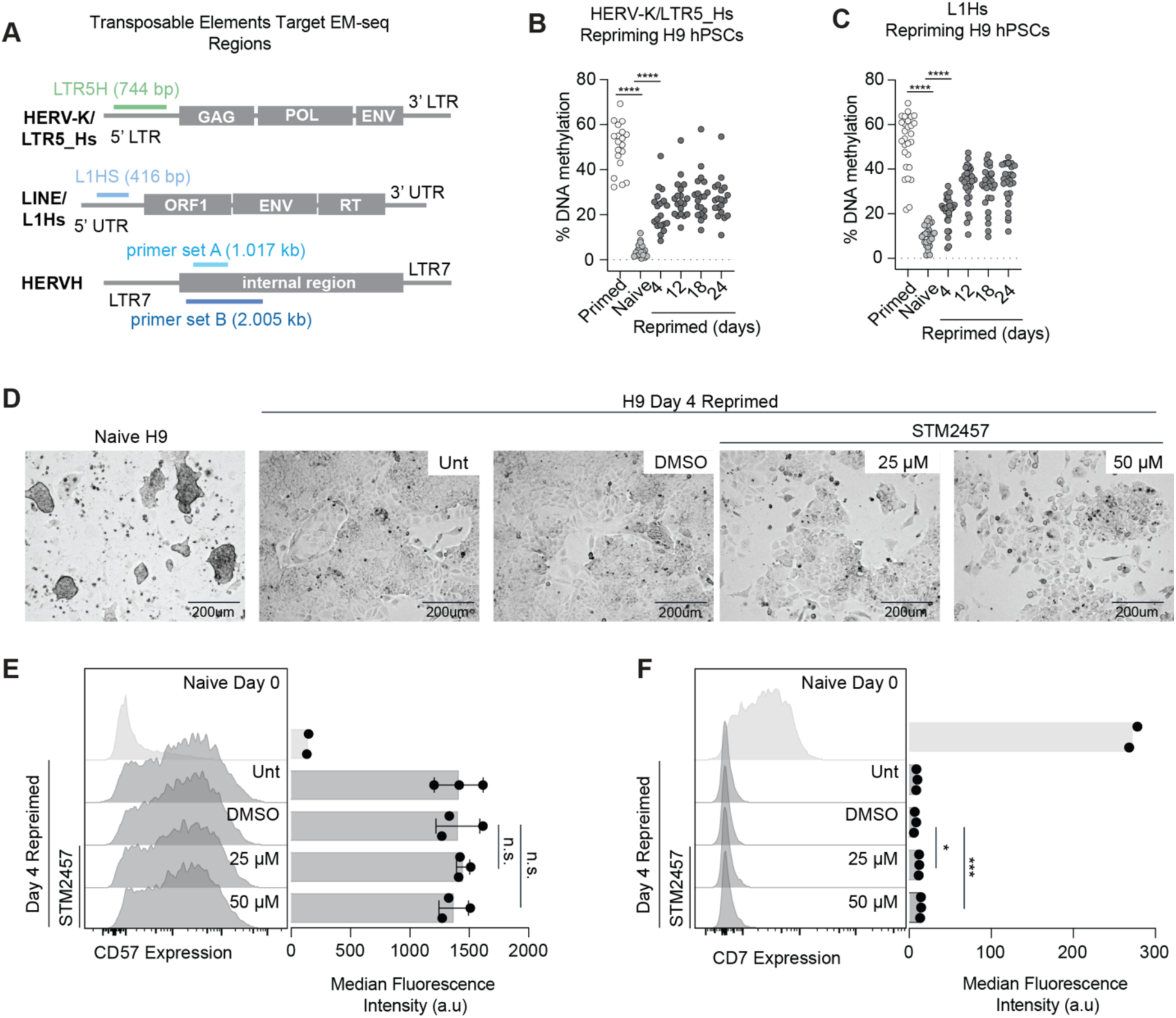
Naive to prime pluripotency transition is not blocked by STM2457 treatment. (A) Schematic of transposable elements and DNA methylation profiling amplicons. (**B** and **C**) Scatter plots showing DNA methylation profiling, where each point represents percent methylation at each CpG at HERV-K/LTR5_Hs (**B**) and L1Hs (**C**) element in the naive H9, and during 24 days of repriming. \*\*\*\**p*< 0.0001, Mann-Whitney U-test, error bars ± 1 s.d. (**D**) Microscopy images of naive H9, and day 4 reprime H9 without treatment (Unt), empty vehicle (DMSO), 25 μM, and 50 μM STM2457 m6A inhibitor. (**E** and **F**) Flow plots and median fluorescence intensity of CD57 (**E**) and CD7 (**F**) expression in naive, day 4 reprimed, day 4 reprimed untreated, day 4 reprimed DMSO-treated, day 4 reprimed 25 μM and 50 μM STM2457 inhibitor-treated cells. n.s. non-significant, \**p*< 0.05, \*\*\**p*< 0.001, unpaired t-test.

**Figure S7.**
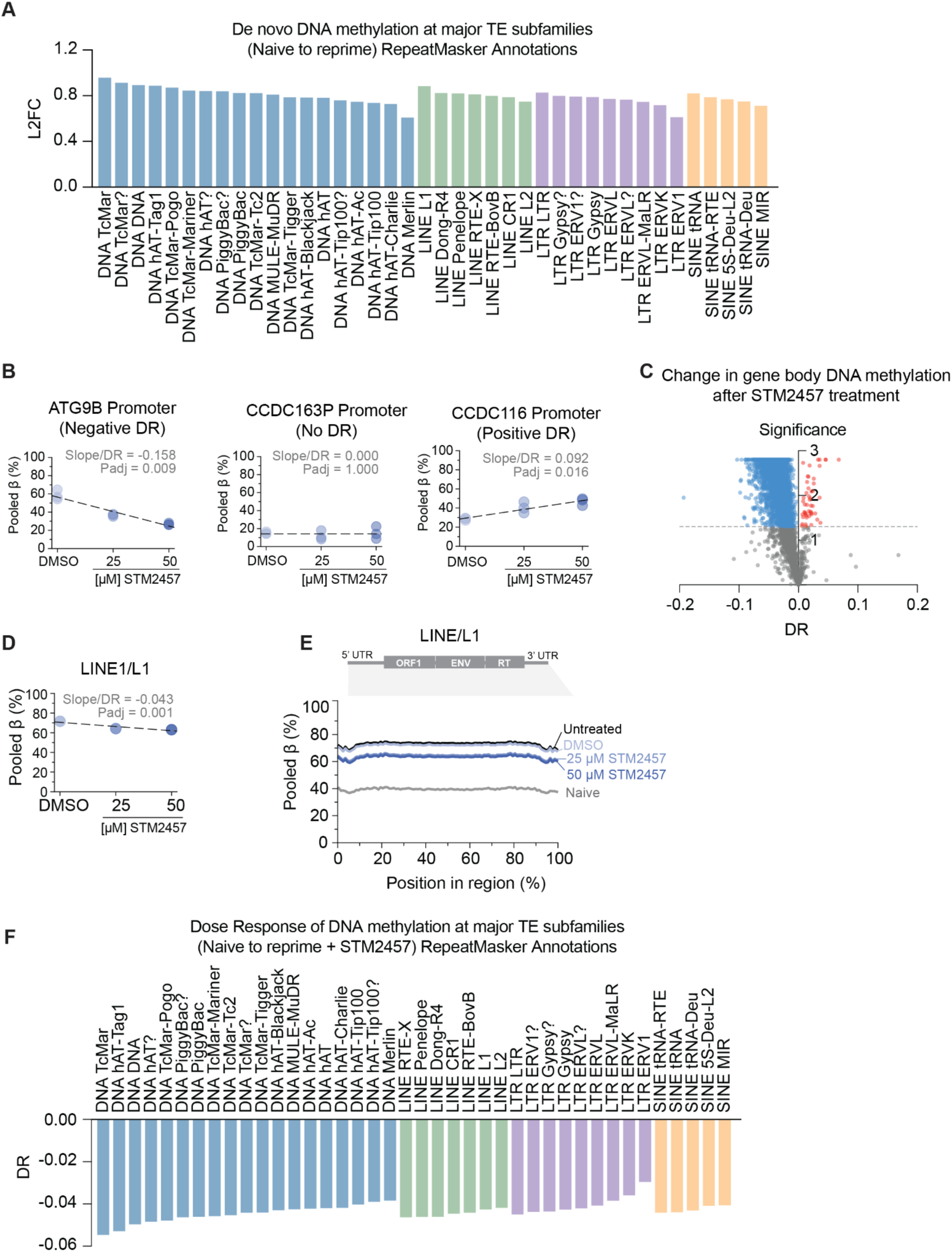
Global interplay between RNA m6A and *de novo* DNA methylation. (A) Bar plot showing the magnitude in gain of novo DNA methylation (Log2 Fold Change, L2FC) at each of the TE subfamilies (as annotated by RepeatMasker). (B) Sample calculations and dose responses at three different regions, which show loss of *de novo* DNA methylation (ATG9B promoter region), no change in *de novo* DNA methylation (CCDC163P region), and gain in *de novo* DNA methylation (CCDC116 promoter). (C) Volcano plot showing dose response of gene bodies after STM2457 treatment, where each point represents a single gene body. Grey points are gene bodies showing non-significant DR change, blue show significantly reduced, and red show significantly increased DR. (D) Scatter plot showing the pooled β-values for LINE consensus across increasing doses of STM2457. The regression slope/dose-response (DR) value and adjusted p-value are displayed. (E) Metagene plot highlighting DNA methylation levels (Pooled β mapped across the normalized length of the LINE consensus sequence in naïve cells, and untreated, DMSO, and dose-dependent STM2457 treatment day 4 reprimed cells. (F) Bar plot showing the dose response at each of the TE subfamilies upon STM2457 treatment.

**Figure S8.**
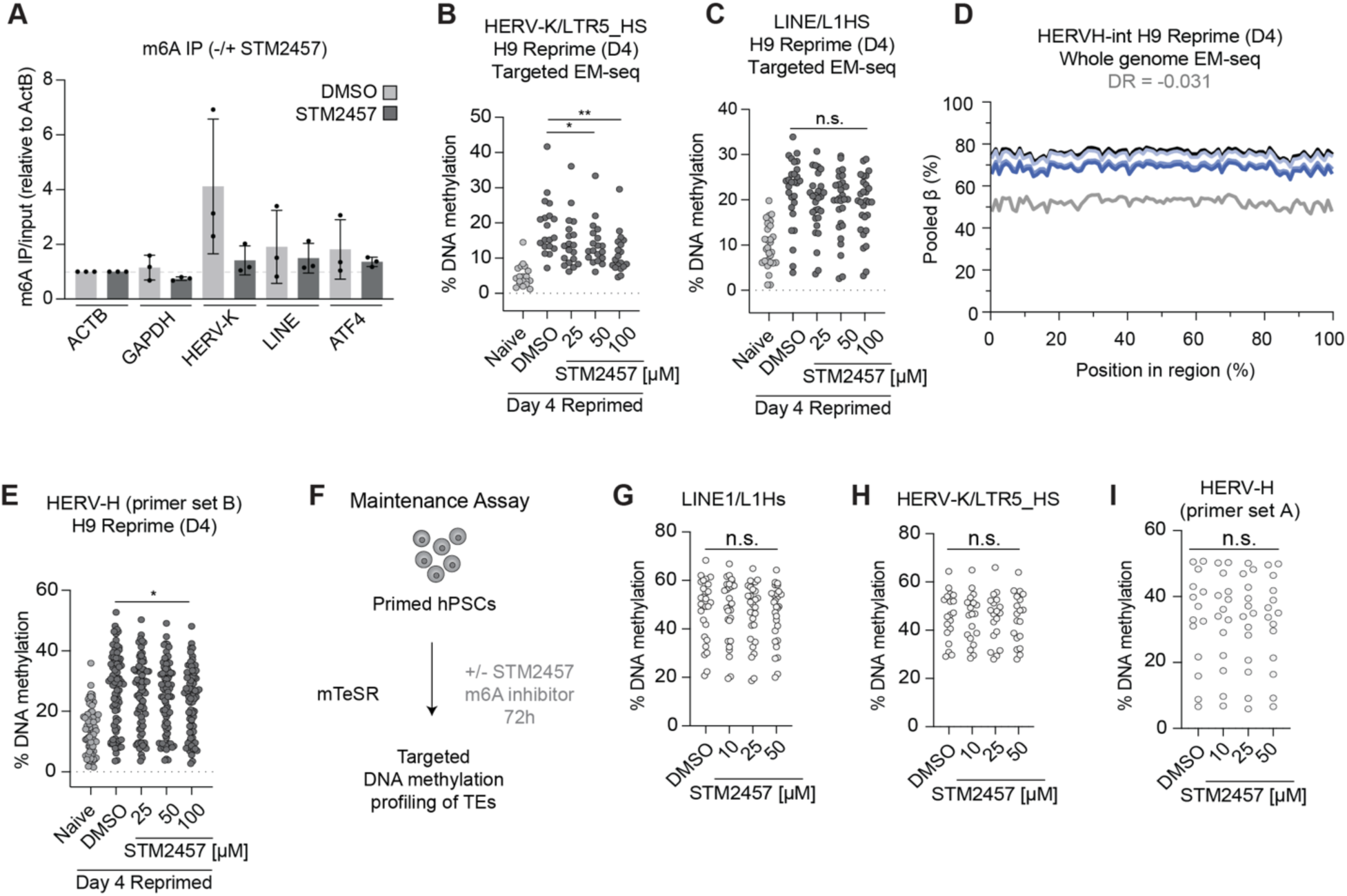
RNA m6A is required for *de novo* DNA methylation, not maintenance, of TE genomic loci. (A) Levels of m6A in LINE-1 and HERV-K RNAs, alongside ACTB, GAPDH, and a known m6A-modified ATF4 RNA in H9 hPSC D4 reprimed cells treated with DMSO and 50 μM STM2457 for 72 hours. n = 3 biological replicates. (**B** and **C**) Scatter plots showing DNA methylation profiling, where each point represents percent methylation at each CpG at HERV-K/LTR5_Hs (**B**), and L1Hs (**C**). n.s. non-significant, \**p*<0.05, \*\**p*<0.01, Mann-Whitney U-test. (**D**) Line graph showing pooled β values across HERVH-int elements in whole genome EM-seq in the DMSO, 25 µM and 50 µM of STM2457 treatment. (**E**) Scatter plots showing DNA methylation profiling, where each point represents percent methylation at each CpG at HERV-H in the naive H9, and day 4 repriming in the presence of empty vehicle (DMSO) or STM2457. \**p*<0.05, Mann-Whitney U-test. (**F**) Schematic of DNA methylation maintenance assay in primed hPSC. (**G** and **H** and **I**) Scatter plots showing DNA methylation profiling, where each point represents percent methylation at each CpG at L1Hs (**G**), HERV-K/LTR5_Hs (**H**), and HERV-H (**I**) amplicon in the primed hPSC treated with STM2457. n.s. non-significant.

## Notes

### Competing Interest Statement

N.S.D. and J.K.N. have filed a patent application related to the CRISPR technologies resulting from the study. J.K.N. and D.X. have filed a patent application related to the delivery of CRISPR epigenome editors. J.K.N. is an inventor of patents related to CRISPRoff/on technologies, filed by The Regents of the University of California.

