## Supplementary Table 3 for "RNA m6A mediates *de novo* DNA methylation and heritable epigenetic silencing in human cells"

**Table S3. Positive hits identified by APEX2-proximity labeling (N-term APEX2-KRAB-dCas9 24 h**

| **Genes** | **Log2(Fold Change)** | **Negative log10 (p-value)** |
| --- | --- | --- |
| SFPQ | 3.09173429 | 1.99234483 |
| NUDT21 | 3.07426424 | 2.00321678 |
| CPSF6 | 3.02559513 | 2.31836165 |
| TRIM28 | 2.93729165 | 2.90783201 |
| PRCC | 2.893834 | 2.24141808 |
| EWSR1 | 2.87404141 | 1.62724835 |
| CPSF7 | 2.83265035 | 1.99926691 |
| FUBP1 | 2.81123 | 2.27423127 |
| CCNK | 2.78246309 | 2.23954692 |
| PABPN1 | 2.76403377 | 1.90082411 |
| RBM10 | 2.75736062 | 2.63407472 |
| RBMX | 2.73230079 | 1.64071864 |
| NONO | 2.69625947 | 2.04710287 |
| SRRT | 2.68718882 | 2.00567216 |
| DDX46 | 2.68536533 | 2.33293393 |
| RBM45 | 2.61298781 | 1.91426987 |
| API5 | 2.60708217 | 1.96929618 |
| SUPT5H | 2.59190791 | 2.05541497 |
| SRSF2 | 2.56019918 | 1.64043824 |
| ZC3H4 | 2.53333087 | 2.28264311 |
| SNRNP70 | 2.51965926 | 1.79706377 |
| CDK12 | 2.49927262 | 2.41377083 |
| TAF15 | 2.47744434 | 1.47160281 |
| CDC5L | 2.47636918 | 2.20819281 |
| KHDC4 | 2.45303876 | 2.12347487 |
| ALYREF | 2.45021556 | 1.61230386 |
| DDX42 | 2.4345712 | 3.41738803 |
| SNRPA | 2.43326723 | 1.64049812 |
| CCAR1 | 2.43109182 | 2.42391044 |
| KHDRBS1 | 2.42157789 | 2.06586185 |
| YLPM1 | 2.39592873 | 2.04081189 |
| ARID1A | 2.39270638 | 2.92082716 |
| NUCKS1 | 2.38281912 | 2.14191508 |
| THRAP3 | 2.36507774 | 3.32379782 |
| PQBP1 | 2.34243636 | 1.84727279 |
| DHX16 | 2.31985721 | 2.77300478 |
| SRSF1 | 2.30863901 | 1.99347832 |
| CWC27 | 2.30816867 | 2.71277666 |
| KHDRBS3 | 2.26530503 | 1.61276417 |
| DHX9 | 2.26421882 | 1.77278999 |
| FIP1L1 | 2.25717771 | 2.59201382 |
| SF3B1 | 2.25234181 | 3.06432138 |
| SUPT6H | 2.23077102 | 2.68867303 |
| PPP1R8 | 2.22537983 | 1.92801619 |
| GTF2F1 | 2.22506241 | 2.68491161 |
| DDX17 | 2.22411757 | 2.00930305 |
| SSBP3 | 2.21662693 | 4.27420556 |
| RBM14 | 2.21501119 | 1.93427721 |
| PLRG1 | 2.20618868 | 2.53280689 |
| GPALPP1 | 2.1972814 | 3.12064836 |
| DDX5 | 2.19603406 | 1.92012344 |
| MCM2 | 2.17374303 | 1.67740381 |
| PSME3 | 2.15560266 | 1.96037113 |
| RPA1 | 2.15054891 | 2.50460803 |
| BCLAF1 | 2.15054399 | 3.43553914 |
| HNRNPA3 | 2.14432669 | 2.01830123 |
| FAM50A | 2.12973003 | 1.9639213 |
| SNW1 | 2.06257355 | 2.79875451 |
| CSTF3 | 2.05171998 | 3.37648033 |
| ZNF207 | 2.03942019 | 1.6279258 |
| CBLL1 | 2.033192 | 2.06782985 |
| SF3A1 | 2.03099522 | 1.65321406 |
| ILF3 | 2.02684487 | 1.82503442 |
| CLASRP | 2.01152002 | 3.88868124 |
| CPSF1 | 2.00655142 | 2.21565224 |
| SCAF4 | 2.00128946 | 2.79202669 |
| NELFA | 1.99659225 | 2.80759911 |
| SRSF9 | 1.99584656 | 1.6694136 |
| RBM26 | 1.9944928 | 3.57280176 |
| ZMIZ1 | 1.99138506 | 1.55135681 |
| WBP4 | 1.99005502 | 1.82833015 |
| C9orf78 | 1.97904836 | 2.10405799 |
| SF3B2 | 1.97862196 | 4.33807624 |
| RBM12 | 1.97824805 | 3.49765833 |
| TRA2B | 1.97724949 | 3.02627068 |
| FUS | 1.97414508 | 2.05343536 |
| TOX4 | 1.96560587 | 4.17482465 |
| FAM98B | 1.95017245 | 1.87192727 |
| HNRNPC | 1.94395618 | 3.26993251 |
| USP39 | 1.9404041 | 2.3460654 |
| HNRNPU | 1.93718337 | 2.06830346 |
| WBP11 | 1.93300134 | 2.20728519 |
| HNRNPA0 | 1.9259551 | 2.4629788 |
| ZFR | 1.92518041 | 1.80401996 |
| CDK8 | 1.91638511 | 1.87126768 |
| DDX39B | 1.91218059 | 2.17295138 |
| RAVER1 | 1.90680434 | 2.19015912 |
| CPSF4 | 1.89571614 | 1.57529533 |
| SFSWAP | 1.88927184 | 4.6938726 |
| SNRPB | 1.88870484 | 1.55255736 |
| HDGF | 1.88257222 | 3.01031629 |
| SRSF11 | 1.8735656 | 2.00111157 |
| SRSF10 | 1.86545619 | 1.49415865 |
| PAXBP1 | 1.86310711 | 1.49141863 |
| RSRC1 | 1.85357183 | 2.46882507 |
| IK | 1.85327548 | 3.34437656 |
| CDK13 | 1.85268209 | 2.09362405 |
| NCOA5 | 1.85137558 | 1.93531755 |
| NFYC | 1.84841271 | 1.63796576 |
| VIRMA | 1.83955555 | 2.74105221 |
| ALKBH5 | 1.83933407 | 1.6335231 |
| BRD2 | 1.83306808 | 2.52987322 |
| SRRM1 | 1.82922205 | 2.17936632 |
| MORF4L1 | 1.82121245 | 2.53690829 |
| MEIS2 | 1.819832 | 1.57029227 |
| SART1 | 1.81944211 | 3.12733753 |
| SNRPD1 | 1.81898636 | 1.92593 |
| GTF2F2 | 1.81769956 | 2.99062458 |
| CDC40 | 1.8108135 | 2.1323902 |
| RBM6 | 1.80957012 | 1.72549386 |
| WDR33 | 1.80344031 | 2.57930527 |
| USF2 | 1.7910854 | 1.79259092 |
| PRPF19 | 1.7887212 | 1.65183053 |
| PPP1R10 | 1.78807163 | 2.97603426 |
| PPWD1 | 1.78129413 | 3.52630532 |
| RPRD2 | 1.77704705 | 6.26965211 |
| RBM15 | 1.77613127 | 1.72724987 |
| MEIS1 | 1.77515704 | 1.67142759 |
| DNAJC8 | 1.7717657 | 3.95402914 |
| HNRNPH2 | 1.76945519 | 1.4490495 |
| SMARCC1 | 1.76444604 | 4.0839527 |
| HSF1 | 1.74821935 | 1.6753654 |
| RAMAC | 1.7455872 | 1.90953108 |
| GTF2E2 | 1.74315672 | 1.91754975 |
| CFDP1 | 1.74273285 | 2.43327297 |
| RBM17 | 1.73677249 | 3.08619899 |
| POU3F2 | 1.73590011 | 1.62489676 |
| NELFCD | 1.71438223 | 3.01543064 |
| RPRD1A | 1.71166778 | 2.05945141 |
| PRPF4 | 1.69372294 | 4.0173031 |
| YJU2 | 1.69039151 | 1.75858622 |
| RFX1 | 1.68943926 | 2.95497652 |
| RBM5 | 1.67954738 | 3.36405929 |
| TCF3 | 1.6736702 | 2.67531686 |
| STRBP | 1.67183721 | 2.48175448 |
| LDB1 | 1.66619698 | 2.3393817 |
| TRA2A | 1.65694977 | 5.69590123 |
| ARID1B | 1.65518111 | 3.00303774 |
| HNRNPK | 1.64774052 | 1.83030228 |
| ERRFI1 | 1.64065432 | 2.72230425 |
| HNRNPH3 | 1.63582081 | 1.45961486 |
| U2AF2 | 1.63500962 | 1.8267956 |
| WTAP | 1.63293883 | 2.05243672 |
| SAP30BP | 1.6293188 | 5.33554227 |
| SF3B4 | 1.62652857 | 1.48031203 |
| RBM27 | 1.61589796 | 2.05968542 |
| ANP32A | 1.60679231 | 2.9138173 |
| POLR2A | 1.60656909 | 2.38222679 |
| SUGP1 | 1.60568226 | 2.05461664 |
| DHX8 | 1.58533132 | 2.01682272 |
| SF1 | 1.58414624 | 1.65674787 |
| DAZAP1 | 1.57102788 | 1.53062914 |
| INTS13 | 1.56939363 | 1.38361214 |
| MED12 | 1.56913831 | 1.96442733 |
| ZNF714 | 1.56758024 | 3.20038384 |
| SIN3A | 1.56416291 | 2.15267995 |
| CLK2 | 1.56175508 | 1.98072878 |
| SNRNP200 | 1.55962311 | 3.3217837 |
| BRD3 | 1.55510586 | 1.46346926 |
| RBM22 | 1.55022143 | 1.94761183 |
| XRCC1 | 1.55014992 | 2.45510578 |
| ZC3H13 | 1.549873 | 3.20538882 |
| FOXA1 | 1.54888631 | 1.88812628 |
| RBBP8 | 1.54776188 | 2.17628978 |
| SNRPD2 | 1.53783561 | 2.08907359 |
| SMU1 | 1.53664147 | 2.74101743 |
| PPIL4 | 1.53359965 | 1.75931081 |
| RBBP6 | 1.52880907 | 3.81703326 |
| GTF2E1 | 1.52560777 | 2.52178048 |
| TLK1 | 1.52557771 | 1.57384695 |
| CDKN2AIPNL | 1.52545087 | 1.79704291 |
| XRCC6 | 1.52175862 | 5.04451067 |
| DHX38 | 1.52083893 | 1.95454226 |
| LSM8 | 1.52034669 | 3.32158777 |
| JUND | 1.5196923 | 1.72648873 |
| MTREX | 1.51888944 | 4.59566344 |
| AQR | 1.5057595 | 3.59359371 |
| METTL14 | 1.50462497 | 3.55115255 |
| SAFB | 1.50400438 | 1.78326004 |
| BRD4 | 1.50219638 | 1.58446343 |
| KHSRP | 1.4955091 | 2.10002479 |
| EYA3 | 1.48978658 | 1.73626983 |
| SRSF6 | 1.48687121 | 3.26261107 |
| MCM3 | 1.48398853 | 2.74997675 |
| CD2BP2 | 1.4819171 | 2.5539637 |
| HMGB3 | 1.48167801 | 2.65706044 |
| PTBP2 | 1.48123667 | 2.19560267 |
| PNISR | 1.48113743 | 1.53077706 |
| SETD1A | 1.47994095 | 3.75342235 |
| SCAF8 | 1.47282196 | 4.25483994 |
| SMARCA4 | 1.47209808 | 2.94042032 |
| SAP18 | 1.47170167 | 2.10761976 |
| CHD2 | 1.47165884 | 2.29770429 |
| ATF7IP | 1.47143926 | 1.30313097 |
| SART3 | 1.46882952 | 4.74741803 |
| NELFE | 1.46321339 | 1.76250729 |
| HNRNPUL2 | 1.46247017 | 1.91518316 |
| CTBP2 | 1.45950622 | 1.58256833 |
| U2SURP | 1.45286683 | 2.33402843 |
| RBM15B | 1.44230603 | 1.61344067 |
| MED1 | 1.44228917 | 1.60409185 |
| PAPOLA | 1.44157663 | 3.47015732 |
| SMARCAD1 | 1.44104587 | 3.89545181 |
| FAM133B | 1.43827094 | 2.11837303 |
| GTF2I | 1.43020643 | 2.41958785 |
| RBM42 | 1.42869505 | 1.71012649 |
| SRSF5 | 1.42415481 | 2.37052629 |
| JUNB | 1.42313562 | 1.5852087 |
| KMT2C | 1.42112567 | 2.27415035 |
| ZNF8 | 1.41710678 | 2.15524714 |
| DRAP1 | 1.40199072 | 1.79959355 |
| SNRPD3 | 1.40052051 | 1.94926597 |
| RALY | 1.39715354 | 2.77044954 |
| PBX2 | 1.395287 | 4.743537 |
| ZC3H14 | 1.39139196 | 2.3688054 |
| GPATCH11 | 1.3890313 | 1.60123442 |
| RPRD1B | 1.3877588 | 2.88911671 |
| C1orf52 | 1.38631105 | 1.53066901 |
| KIAA1143 | 1.38336314 | 1.94802793 |
| NKTR | 1.38323839 | 1.68451641 |
| LEF1 | 1.3799451 | 1.67805253 |
| CCDC174 | 1.37823395 | 2.58448805 |
| GATA4 | 1.37705322 | 1.68909452 |
| HMBOX1 | 1.36718286 | 1.84086253 |
| RBM4 | 1.34770005 | 1.55202553 |
| SOX9 | 1.34182883 | 1.37026592 |
| MED14 | 1.338731 | 2.67946138 |
| ZRANB2 | 1.33678388 | 1.34976181 |
| CMTR1 | 1.33669584 | 2.27819214 |
| ZMAT2 | 1.33490641 | 1.86011905 |
| SOX2 | 1.33439133 | 1.72434209 |
| CREBBP | 1.33342608 | 2.06431056 |
| DONSON | 1.32979364 | 1.93453552 |
| LENG1 | 1.32741224 | 1.57015139 |
| SNRNP27 | 1.32651133 | 1.55418942 |
| MORC4 | 1.3259757 | 2.75404746 |
| CTBP1 | 1.32453248 | 1.80079565 |
| CCNT2 | 1.32198865 | 2.46601691 |
| CSTF2 | 1.3206322 | 1.82032706 |
| POLDIP3 | 1.31974612 | 1.70770876 |
| PRPF40A | 1.31676281 | 3.07864475 |
| XPA | 1.31625102 | 1.78820478 |
| SNRNP40 | 1.31427381 | 3.87577748 |
| HNRNPM | 1.31409043 | 1.80366783 |
| CWC15 | 1.3127048 | 1.58622065 |
| CRNKL1 | 1.31257735 | 2.03650561 |
| WAC | 1.3100998 | 4.85912008 |
| ATF1 | 1.30955642 | 2.06392523 |
| UBE2T | 1.3095199 | 1.52252137 |
| FAM76A | 1.30918549 | 2.81035145 |
| ZNF326 | 1.30583685 | 2.4949236 |
| HDGFL3 | 1.30356393 | 1.47870571 |
| PPIL2 | 1.30187202 | 1.54757893 |
| LENG8 | 1.29879203 | 2.04391846 |
| BICRA | 1.29324504 | 2.07170377 |
| RSRC2 | 1.29046396 | 4.07166146 |
| NSD3 | 1.28882998 | 2.33034716 |
| NCOR1 | 1.28780617 | 2.85778097 |
| GINS3 | 1.28090865 | 2.41730154 |
| GPKOW | 1.2801038 | 2.61505036 |
| NCBP3 | 1.27769767 | 4.60585457 |
| DLX5 | 1.27476621 | 1.47755782 |
| SF3A2 | 1.27178105 | 1.72914658 |
| KLF3 | 1.27090015 | 1.5448249 |
| SUB1 | 1.27017881 | 3.27036889 |
| RBM4B | 1.26941417 | 1.85934899 |
| VRK1 | 1.26633038 | 2.65389822 |
| IWS1 | 1.26613616 | 2.93132656 |
| PRPF8 | 1.2659317 | 2.42935949 |
| SMARCD2 | 1.26366661 | 1.66979299 |
| ZCCHC8 | 1.26040171 | 4.93005813 |
| SAFB2 | 1.26001099 | 1.84392748 |
| MCM6 | 1.25583918 | 2.10673649 |
| SMARCD3 | 1.25299481 | 3.0630576 |
| HNRNPF | 1.25129266 | 1.70334268 |
| RBM7 | 1.24694338 | 1.9744597 |
| SETD1B | 1.24488919 | 1.77201318 |
| SMARCC2 | 1.2427787 | 1.46005599 |
| NRF1 | 1.24150521 | 1.38368479 |
| DIDO1 | 1.24061026 | 1.82839334 |
| DACH1 | 1.23684314 | 1.53416602 |
| CTDSPL2 | 1.23425957 | 2.01061552 |
| SLU7 | 1.2340628 | 2.5282506 |
| NCOA3 | 1.22969385 | 1.85085888 |
| TFAP2C | 1.22775834 | 1.37523177 |
| CDK11B | 1.22709273 | 2.8340699 |
| ZNF830 | 1.224456 | 2.28269769 |
| ZMIZ2 | 1.22074895 | 1.53920342 |
| SON | 1.21899596 | 3.41608373 |
| SRSF4 | 1.21744135 | 4.93690726 |
| SLTM | 1.21549491 | 1.6996344 |
| DPF2 | 1.21537722 | 4.20689284 |
| CPSF2 | 1.21022619 | 1.85449089 |
| CHD1 | 1.20850494 | 2.26259222 |
| PUF60 | 1.20724486 | 4.70967397 |
| YTHDC1 | 1.20602473 | 2.59208874 |
| QRICH1 | 1.20534977 | 3.45532075 |
| FYTTD1 | 1.20070756 | 3.36459888 |
| NKX6-1 | 1.19512141 | 1.81533857 |
| KMT2D | 1.19184026 | 2.44374509 |
| RBM25 | 1.19160711 | 1.89616777 |
| TRIM24 | 1.18826619 | 2.72740378 |
| SMARCD1 | 1.18675702 | 3.08390934 |
| RP9 | 1.18642714 | 1.75464604 |
| LUC7L2 | 1.18357779 | 2.13254919 |
| CHTOP | 1.17864488 | 2.64332075 |
| GTF2H1 | 1.17697315 | 3.73001998 |
| HMGN5 | 1.17265692 | 3.57512035 |
| CDKN2AIP | 1.17184334 | 2.38388474 |
| NKAP | 1.17163224 | 2.02766122 |
| TP73 | 1.16932472 | 2.27188451 |
| PRPF4B | 1.1693117 | 2.66899076 |
| TOP1 | 1.1692046 | 1.6007728 |
| SMARCA2 | 1.16904972 | 3.83532502 |
| ZFC3H1 | 1.16889439 | 3.42336032 |
| CHD7 | 1.1688526 | 2.26056633 |
| XAB2 | 1.16860119 | 4.0323351 |
| PRPF38A | 1.16700193 | 1.5107293 |
| HMGXB4 | 1.1664641 | 3.26049706 |
| MED13 | 1.16644309 | 2.09487188 |
| CSTF1 | 1.16643944 | 2.47488106 |
| SRRM2 | 1.16459155 | 3.1720323 |
| FOXK2 | 1.16231764 | 2.09581564 |
| SMNDC1 | 1.16183105 | 1.79894641 |
| WDR70 | 1.16063827 | 2.03691501 |
| RBFOX2 | 1.16009601 | 2.31459333 |
| GTF2B | 1.15724858 | 1.61955565 |
| SPEN | 1.15709907 | 2.24709362 |
| SF3B6 | 1.15607536 | 3.91471919 |
| POGK | 1.15316215 | 2.79350295 |
| EP300 | 1.15234123 | 3.29148963 |
| SAP130 | 1.14847133 | 5.19685993 |
| SRSF3 | 1.14526091 | 2.75517569 |
| HMGB1 | 1.14435336 | 2.59528276 |
| POLA1 | 1.14271193 | 3.78746482 |
| RNF20 | 1.13882807 | 2.58538553 |
| AKAP8 | 1.13731413 | 1.83597644 |
| ZNF638 | 1.13713396 | 3.03833042 |
| GREB1L | 1.13126328 | 2.44617846 |
| TBP | 1.13057203 | 1.39035478 |
| RFX2 | 1.12624149 | 2.86005692 |
| ZNF250 | 1.12380652 | 1.89554003 |
| TASOR | 1.12272861 | 4.26678894 |
| IRF2BP1 | 1.11615521 | 2.28581564 |
| HMX1 | 1.11468402 | 1.67865211 |
| RBM18 | 1.11458859 | 1.7684847 |
| SRSF7 | 1.11439257 | 4.03559608 |
| TPR | 1.11321885 | 2.56545161 |
| PPP4R3B | 1.11314916 | 2.29986845 |
| PSME3IP1 | 1.11253181 | 2.3252447 |
| CWC25 | 1.11206605 | 2.08642795 |
| DDX41 | 1.10907586 | 2.13680157 |
| LUC7L | 1.10880675 | 1.8029623 |
| U2AF1 | 1.10849555 | 1.62801004 |
| ESS2 | 1.10415512 | 2.24453232 |
| TCERG1 | 1.10287694 | 2.45917947 |
| TAF3 | 1.10281764 | 1.80280467 |
| NKAPD1 | 1.10277149 | 1.96585466 |
| NIPBL | 1.10256456 | 3.66762992 |
| PRDM10 | 1.10009034 | 2.73953559 |
| MSX2 | 1.09977487 | 2.60941989 |
| SMARCE1 | 1.09863242 | 4.01092116 |
| HMCES | 1.09649857 | 1.53007053 |
| KRT5 | 1.09626245 | 1.39889228 |
| RNF40 | 1.0948848 | 6.31393955 |
| RBM20 | 1.09401308 | 4.77304711 |
| TERF2 | 1.0914573 | 1.78509734 |
| CIR1 | 1.08742684 | 1.88051403 |
| CBX5 | 1.08625056 | 1.86860189 |
| TCEAL4 | 1.08603433 | 2.22967736 |
| ZNF768 | 1.08439628 | 1.79834128 |
| SUPT16H | 1.07974559 | 1.97505302 |
| NFIA | 1.07780848 | 1.83969595 |
| MORC3 | 1.07747611 | 2.99824313 |
| ZC3H10 | 1.07591367 | 2.7350865 |
| SAE1 | 1.07478063 | 2.19567565 |
| ZC3H6 | 1.07284679 | 2.12794014 |
| NFIC | 1.07209645 | 1.74039832 |
| FAM169A | 1.06834777 | 1.42779195 |
| RNF113A | 1.06734661 | 1.92063952 |
| ERCC3 | 1.0649936 | 3.77076426 |
| DBR1 | 1.06459069 | 1.73008362 |
| NXF1 | 1.06234733 | 1.66187694 |
| DDX39A | 1.06034695 | 3.24313301 |
| NOL4L | 1.06022693 | 2.65628968 |
| NFIB | 1.05922847 | 1.64333093 |
| TCF12 | 1.05864313 | 2.87208032 |
| AFF4 | 1.05669516 | 1.87494267 |
| ERH | 1.0551839 | 1.83880292 |
| RESF1 | 1.05332818 | 2.43872941 |
| ZIC2 | 1.05235451 | 1.55566399 |
| MCM4 | 1.04927793 | 4.7233492 |
| ZFP90 | 1.04897685 | 3.17519154 |
| TEAD1 | 1.04673946 | 1.60853349 |
| RBM12B | 1.04510043 | 1.4341403 |
| SRSF12 | 1.04177752 | 2.23985928 |
| PRR12 | 1.04064226 | 2.26034051 |
| SIX4 | 1.03960197 | 2.39353655 |
| GSE1 | 1.03795514 | 2.74048214 |
| BCL9L | 1.03659197 | 1.99917559 |
| PRPF18 | 1.03574112 | 1.80204641 |
| TASOR2 | 1.03546674 | 3.3550782 |
| DNMT1 | 1.03371518 | 1.68674326 |
| POLR3F | 1.03106907 | 1.89501343 |
| PPIG | 1.03069474 | 2.26291156 |
| HMGB2 | 1.02968806 | 2.67951345 |
| TOP2B | 1.02948072 | 2.71433674 |
| ZNF609 | 1.02181116 | 2.92033773 |
| TAF1L | 1.02151335 | 1.82824858 |
| MED17 | 1.02142232 | 1.58036468 |
| SNIP1 | 1.01981766 | 2.34319909 |
| NCOA6 | 1.01821397 | 1.40270115 |
| USP37 | 1.01811173 | 4.11228686 |
| DHX15 | 1.01156687 | 2.09122383 |
| XRCC4 | 1.01098038 | 1.88249249 |
| ZNF436 | 1.01070806 | 1.30621253 |
| PSMC3IP | 1.00978555 | 1.77468824 |
| PCF11 | 1.00940427 | 3.25463457 |
| ZNF780A | 1.00836022 | 1.58124959 |
| ZBTB7A | 1.00798769 | 1.9034064 |
| JUN | 1.00680681 | 2.66597798 |
| RPA3 | 1.00020098 | 1.4687825 |
| CWF19L2 | 1.00004011 | 2.36209119 |
| BUB3 | 0.99945396 | 1.74375976 |
| RBM39 | 0.99323544 | 2.45062732 |
| TAF1 | 0.99255721 | 1.87964171 |
| PAXIP1 | 0.99224953 | 2.29890015 |
| SETD2 | 0.99011204 | 3.13880422 |
| TERF2IP | 0.98906972 | 1.4525953 |

|  |  |  |
| --- | --- | --- |
| CWC22 | 0.98793808 | 2.03418148 |
| CXXC1 | 0.98656689 | 2.99160205 |
| BCL9 | 0.98469958 | 2.52817301 |
| MAML1 | 0.98325043 | 2.2971324 |
| GPS2 | 0.98191893 | 1.37066182 |
| MED25 | 0.98012487 | 2.94328875 |
| BOD1L1 | 0.97887003 | 2.85122047 |
| PBRM1 | 0.97798602 | 2.12177718 |
| NSRP1 | 0.97625674 | 1.59673 |
| REV1 | 0.97535662 | 2.70131151 |
| RFC1 | 0.97491532 | 1.89627029 |
| POLE3 | 0.97422084 | 1.74420378 |
| TCF20 | 0.97406736 | 1.96776585 |
| INO80C | 0.97176873 | 2.06347916 |
| CREB1 | 0.97094674 | 1.47890657 |
| PWP2 | 0.96877783 | 1.93772076 |
| ACIN1 | 0.96772742 | 2.2256016 |
| STAG2 | 0.96763665 | 1.47434054 |
| PRPF3 | 0.96620044 | 1.87485952 |
| TTC14 | 0.96541662 | 2.62052909 |
| IRF2 | 0.96436895 | 1.97656728 |
| KDM6A | 0.96228096 | 4.66198484 |
| NFATC2IP | 0.95945689 | 1.64409887 |
| BAP18 | 0.95853738 | 1.64345744 |
| PNKP | 0.95795481 | 3.85326992 |
| SSRP1 | 0.95709126 | 1.39136233 |
| TCEA1 | 0.95669156 | 2.89322243 |
| DNAJC17 | 0.95644799 | 1.45161567 |
| NCOR2 | 0.95361429 | 2.24807803 |
| RAVER2 | 0.95315683 | 3.25693691 |
| TLK2 | 0.95169611 | 3.54925481 |
| DLX1 | 0.9508995 | 1.51133419 |
| RNMT | 0.95082578 | 1.42786044 |
| PRIM2 | 0.94982371 | 3.07093884 |
| PRPF6 | 0.94978718 | 1.49413343 |
| XRN2 | 0.9480426 | 1.9287216 |
| THOC1 | 0.94214416 | 2.47308724 |
| PPP4R2 | 0.94149185 | 2.57767203 |
| ZBTB43 | 0.9394147 | 4.07095792 |
| SUPT20H | 0.93938401 | 2.39104918 |
| ZNF689 | 0.93937722 | 1.51449211 |
| METTL3 | 0.93923848 | 2.00982391 |
| ZNF263 | 0.93833784 | 3.55547594 |
| ELF1 | 0.93554855 | 1.36390784 |
| EXOSC10 | 0.93472339 | 3.09508336 |
| CUX2 | 0.9340242 | 2.60513361 |
| BRPF3 | 0.93364916 | 2.52960036 |
| RFX3 | 0.93147745 | 2.56332868 |

|  |  |  |
| --- | --- | --- |
| RBM8A | 0.93094669 | 2.3916112 |
| TRIM27 | 0.92933934 | 1.68528531 |
| SOX12 | 0.92932384 | 1.36601752 |
| RNPC3 | 0.92878187 | 2.55381769 |
| INO80 | 0.92860289 | 2.19800645 |
| POGZ | 0.92642089 | 2.21529333 |
| MED15 | 0.92471386 | 2.63966248 |
| HES1 | 0.92320775 | 1.67183483 |
| GATA6 | 0.92212392 | 1.44593795 |
| PRPF31 | 0.91652255 | 2.26555183 |
| MAGOHB | 0.91536909 | 2.05025245 |
| ZNF639 | 0.91117977 | 1.43109224 |
| SCAF11 | 0.91014757 | 2.16446968 |
| QSER1 | 0.90983219 | 3.28832851 |
| INTS10 | 0.90766757 | 2.65102983 |
| PDCD7 | 0.90617444 | 2.5789996 |
| NCOA2 | 0.90532096 | 3.71978806 |
| MCM5 | 0.90504575 | 1.60741687 |
| MBTD1 | 0.90482887 | 2.44494025 |
| DHX40 | 0.90468001 | 1.60907359 |
| WRNIP1 | 0.90451646 | 2.6397014 |
| HDAC2 | 0.90435529 | 2.40036818 |
| LSM10 | 0.9033317 | 1.73254804 |
| PAX9 | 0.90279929 | 1.77942967 |
| KMT2E | 0.90214911 | 2.50065655 |
| DCAF13 | 0.90079122 | 2.45578332 |
| DGCR8 | 0.89935083 | 2.56754183 |
| MIER1 | 0.89869889 | 3.5335534 |
| MLLT1 | 0.89592893 | 1.63559781 |
| TP53BP1 | 0.89418787 | 3.15399931 |
| LIG1 | 0.89119648 | 2.33356068 |
| SUDS3 | 0.88654618 | 1.70557354 |
| PKNOX1 | 0.88621092 | 1.51630096 |
| SBNO2 | 0.88573556 | 2.18567935 |
| MORF4L2 | 0.88382197 | 1.48194364 |
| GATAD1 | 0.8834068 | 1.75202539 |
| ING3 | 0.88325469 | 2.68074722 |
| CASZ1 | 0.88292137 | 1.33107106 |
| SPAG7 | 0.88182058 | 1.45243188 |
| HEXIM1 | 0.8812136 | 1.79502362 |
| PARN | 0.87924548 | 4.73799149 |
| USP7 | 0.87842653 | 2.06582768 |
| NKX2-4 | 0.87740096 | 1.97363835 |
| ZNF318 | 0.87499209 | 5.21436147 |
| PPIE | 0.87447735 | 2.0895529 |
| KCTD18 | 0.87396084 | 2.99341341 |
| RAD21 | 0.86830919 | 1.64972905 |
| BRMS1L | 0.86762554 | 1.70371886 |

|  |  |  |
| --- | --- | --- |
| TET2 | 0.86745972 | 3.18851508 |
| SNRPE | 0.86396898 | 1.58734417 |
| PARP1 | 0.86381167 | 3.34004347 |
| NELFB | 0.86361937 | 2.93653592 |
| IRF2BPL | 0.86262797 | 1.48323859 |
| PIAS4 | 0.85493646 | 2.29116594 |
| DMAP1 | 0.85419788 | 2.32895601 |
| SMC6 | 0.8535483 | 1.68875952 |
| REXO1 | 0.85232339 | 2.08766397 |
| BDP1 | 0.85214625 | 2.1857721 |
| THOC2 | 0.85169295 | 2.81870898 |
| GPATCH1 | 0.85107278 | 2.96069924 |
| CDC14B | 0.85104127 | 1.61448741 |
| DERPC | 0.85014111 | 2.08059769 |
| ATF7 | 0.84836191 | 1.88506811 |
| ZNF398 | 0.84765257 | 1.49370116 |
| BCOR | 0.84628861 | 2.63639344 |
| SMN2 | 0.84570978 | 2.61852014 |
| ZNF746 | 0.84547207 | 1.49450174 |
| ZHX2 | 0.84410128 | 3.47181187 |
| AKAP8L | 0.8435974 | 2.00553012 |
| ZBTB45 | 0.84237516 | 2.42421581 |
| SNRPB2 | 0.8422851 | 3.4246413 |
| C1orf131 | 0.84124972 | 1.79243504 |
| XRCC5 | 0.84066716 | 2.33849884 |
| ZHX3 | 0.83784623 | 3.26927854 |
| WDR3 | 0.83628105 | 2.67552726 |
| LIN54 | 0.83484704 | 2.55344391 |
| KDM4A | 0.83451381 | 1.30532261 |
| PAPOLG | 0.83297298 | 3.23941639 |
| SPTY2D1 | 0.83212407 | 1.64562483 |
| MAX | 0.83192014 | 2.70323855 |
| CECR2 | 0.83125459 | 1.98044668 |
| MAD1L1 | 0.83024728 | 2.94362519 |
| USP48 | 0.82922202 | 1.36121953 |
| TAF5 | 0.82853873 | 2.21751636 |
| PHC3 | 0.82696529 | 1.98460825 |
| CENPI | 0.82582292 | 1.60125629 |
| RTF1 | 0.82424232 | 2.26536671 |
| FOXK1 | 0.82319027 | 1.86725579 |
| NUSAP1 | 0.82147868 | 1.85790049 |
| CCDC12 | 0.82017358 | 2.62997692 |
| ARID4A | 0.81885485 | 1.4537092 |
| MARCHF7 | 0.81846226 | 2.37756996 |
| CENPF | 0.81789997 | 2.26886199 |
| GINS1 | 0.81420804 | 2.05084641 |
| PRDM15 | 0.81420327 | 1.93969072 |
| YY1 | 0.81308327 | 2.95251061 |

|  |  |  |
| --- | --- | --- |
| ZC3H18 | 0.81134723 | 2.42122967 |
| TFAP2A | 0.80975627 | 1.3476849 |
| ZBTB34 | 0.80810343 | 2.1118878 |
| ZCCHC9 | 0.80783734 | 2.13932427 |
| JMJD1C | 0.80742232 | 3.21527459 |
| HELLS | 0.80662295 | 3.42595642 |
| ZEB1 | 0.80382452 | 2.80251817 |
| MLLT10 | 0.80170068 | 2.72535394 |
| NAB2 | 0.79897217 | 1.67540291 |
| MSH6 | 0.79882557 | 2.52777648 |
| CBX3 | 0.79807074 | 2.24273719 |
| GTF2IRD1 | 0.79794983 | 2.99780224 |
| TRIM33 | 0.79669291 | 2.4503183 |
| TTC37 | 0.79603772 | 2.2605552 |
| C5orf22 | 0.79387555 | 1.67171837 |
| SCAF1 | 0.79308313 | 3.44121593 |
| HSF2 | 0.79250244 | 2.57597161 |
| MEAF6 | 0.79118156 | 1.6009062 |
| TAF6 | 0.79112375 | 2.2035386 |
| ASH2L | 0.79076534 | 3.85410381 |
| ZRSR2 | 0.79063357 | 2.85006291 |
| GTF2H4 | 0.78883894 | 1.84507797 |
| FAM76B | 0.78861516 | 1.79213932 |
| ADARB1 | 0.78768714 | 1.79448567 |
| PHRF1 | 0.78718955 | 2.71380438 |
| RADX | 0.78684744 | 2.82919785 |
| INTS4 | 0.78648496 | 3.01901903 |
| POLR3D | 0.78519417 | 2.15302298 |
| SCML2 | 0.78510831 | 2.77073755 |
| SOX13 | 0.78484131 | 1.41043255 |
| BRD7 | 0.78444817 | 2.42750615 |
| APOBEC3B | 0.78395096 | 2.26305442 |
| ADNP | 0.78371042 | 3.04501276 |
| MPHOSPH8 | 0.78347984 | 3.03896993 |
| CEP164 | 0.78288802 | 1.46478404 |
| HTATSF1 | 0.78077288 | 1.39299434 |
| NACC1 | 0.77891534 | 1.91135067 |
| RACGAP1 | 0.77281376 | 4.37471615 |
| RGS10 | 0.76957447 | 1.77381982 |
| FRG1 | 0.76833809 | 1.36598529 |
| SNRNP48 | 0.76807943 | 3.07414614 |
| RIF1 | 0.76744355 | 4.25482237 |
| PHF21A | 0.76728769 | 2.13765434 |
| PDS5B | 0.76621947 | 1.83965502 |
| SETD5 | 0.76502928 | 3.9676166 |
| ATF2 | 0.76428156 | 2.71666548 |
| RECQL5 | 0.76290594 | 3.16379514 |
| SREK1 | 0.76278732 | 1.44932004 |

|  |  |  |
| --- | --- | --- |
| ZEB2 | 0.76100141 | 3.86829801 |
| TGM1 | 0.7607171 | 1.6765901 |
| TRPS1 | 0.76037805 | 2.33652661 |
| GEMIN4 | 0.76025922 | 1.88367796 |
| HDAC3 | 0.75558264 | 1.53051255 |
| MIDEAS | 0.75516981 | 1.86905077 |
| BRD9 | 0.75401527 | 3.25824376 |
| SMARCAL1 | 0.75363898 | 2.2220161 |
| ZNF711 | 0.75210283 | 1.3253524 |
| DUSP11 | 0.74786554 | 2.1087169 |
| NFRKB | 0.74741176 | 2.24981114 |
| SREK1IP1 | 0.7472736 | 2.05901973 |
| ERCC1 | 0.745313 | 2.97868133 |
| ZNF608 | 0.74069875 | 1.51370204 |
| SMARCB1 | 0.7389129 | 1.94659479 |
| RBM33 | 0.73845507 | 2.44929162 |
| ZNF217 | 0.73802585 | 2.17071368 |
| MSH2 | 0.73763766 | 2.2270121 |
| CASP8AP2 | 0.73691439 | 2.12745116 |
| PRPF39 | 0.7361394 | 2.23635054 |
| THOC5 | 0.73535623 | 3.17030341 |
| FKBP5 | 0.73284084 | 2.4710522 |
| PAF1 | 0.7317704 | 2.89610973 |
| GPATCH8 | 0.73106746 | 4.07768777 |
| GTF3C3 | 0.7310573 | 2.21334877 |
| KLF12 | 0.73089803 | 1.53902238 |
| MRPS7 | 0.73086569 | 1.37247218 |
| ZBTB10 | 0.73056328 | 3.62311572 |
| PTOV1 | 0.72887786 | 1.62521391 |
| TIMELESS | 0.72780558 | 2.75745095 |
| PHF2 | 0.72767459 | 3.36160301 |
| ZMYM4 | 0.72728214 | 2.42214063 |
| POLE | 0.72669723 | 3.50214151 |
| INTS12 | 0.72616077 | 1.80926375 |
| MRGBP | 0.72483949 | 2.50521845 |
| MMS22L | 0.7236956 | 1.40165571 |
| DNMT3B | 0.7236724 | 1.63999597 |
| PHIP | 0.72296453 | 1.48515209 |
| TBL1XR1 | 0.72285996 | 2.42843972 |
| MTA3 | 0.72256036 | 3.12522519 |
| MAU2 | 0.72196218 | 2.9119239 |
| FEN1 | 0.72195733 | 1.95490241 |
| ZNF148 | 0.72142343 | 2.90605051 |
| DDX23 | 0.71902255 | 2.24584578 |
| EP400 | 0.71783765 | 3.97588558 |
| GATAD2B | 0.7156624 | 2.17812519 |
| FOXJ2 | 0.71434268 | 1.76928159 |
| RNASEH1 | 0.71399354 | 3.59019794 |

|  |  |  |
| --- | --- | --- |
| KDM3B | 0.71300802 | 2.34884495 |
| MSL3 | 0.70987889 | 3.26848199 |
| FRA10AC1 | 0.70958144 | 1.67977848 |
| ZBTB8A | 0.70885228 | 1.74098637 |
| PARP2 | 0.70749211 | 2.19094523 |
| PRAME | 0.70726458 | 1.53814962 |
| SUGP2 | 0.70532585 | 4.57158401 |
| GABPA | 0.70442917 | 1.82998146 |
| ZC3H3 | 0.70390689 | 3.60808197 |
| KDM6B | 0.70375232 | 1.89258869 |
| KDM5C | 0.70318223 | 3.27083752 |
| ARID3B | 0.70314697 | 4.86821859 |
| LEO1 | 0.70286622 | 1.83659474 |
| CHD8 | 0.70286267 | 1.70439131 |
| ZNF16 | 0.70273162 | 2.13648744 |
| BRCA1 | 0.70137081 | 4.10490976 |
| BPTF | 0.70021971 | 3.65818278 |
| GTF3C1 | 0.69984456 | 2.34705442 |
| BAG4 | 0.69936955 | 1.46126977 |
| TAF4 | 0.69869463 | 1.56020936 |
| ZBTB3 | 0.69775116 | 1.4363888 |
| KDM5A | 0.69761027 | 1.99248857 |
| USP42 | 0.69397224 | 3.13884727 |
| BRF1 | 0.69341316 | 2.03237233 |
| RBMX2 | 0.69274216 | 1.63739599 |
| ZNF629 | 0.69246199 | 2.67229279 |
| ZMYM2 | 0.69177215 | 2.16339936 |
| PRC1 | 0.69147869 | 1.81359799 |
| MAGED1 | 0.69137855 | 1.74227175 |
| E2F6 | 0.69119167 | 1.87252003 |
| CCDC117 | 0.68961942 | 3.03834226 |
| POLR3G | 0.68924283 | 1.348668 |
| BLM | 0.68783245 | 2.03334611 |
| ZC3H8 | 0.68703084 | 1.65485359 |
| PHF20L1 | 0.68696722 | 1.74207527 |
| TENT2 | 0.68659758 | 2.77700196 |
| PBX3 | 0.68636738 | 2.25172407 |
| THAP11 | 0.68466761 | 1.5410842 |
| NKX2-1 | 0.68452475 | 1.54710793 |
| ZNF777 | 0.68389778 | 1.82649209 |
| FAM126B | 0.68176114 | 1.90840113 |
| JADE3 | 0.6816097 | 1.78909148 |
| JMJD6 | 0.67945646 | 2.01300192 |
| ZBTB9 | 0.67731141 | 1.78812119 |
| ADAR | 0.6768737 | 1.7214625 |
| KDM1A | 0.67576979 | 2.66323056 |
| BANP | 0.67543282 | 2.34031408 |
| ORC2 | 0.67533235 | 2.25132087 |

|  |  |  |
| --- | --- | --- |
| MTA2 | 0.67483451 | 2.99789312 |
| PCIF1 | 0.67293675 | 2.50244993 |
| CDK7 | 0.67192914 | 2.16016463 |
| ZNF317 | 0.67040525 | 1.68928868 |
| SP1 | 0.67032044 | 1.35164778 |
| BCLAF3 | 0.66987494 | 1.73416129 |
| UTP3 | 0.66947679 | 1.66341837 |
| EMSY | 0.66912521 | 2.3795336 |
| L3MBTL2 | 0.66802771 | 2.82679067 |
| CCDC82 | 0.66766436 | 3.33798945 |
| NCBP1 | 0.66664671 | 1.88846207 |
| TOP2A | 0.66597104 | 2.41389158 |
| RFWD3 | 0.66558372 | 1.69242083 |
| USP3 | 0.66378505 | 2.25167382 |
| RCOR1 | 0.66376566 | 2.18041464 |
| APEX1 | 0.66326992 | 1.81662204 |
| GATA2 | 0.6632698 | 1.37865746 |
| MECOM | 0.66229911 | 2.64840523 |
| MCM8 | 0.66172224 | 1.40294049 |
| E2F7 | 0.66086835 | 1.38324637 |
| ZBTB33 | 0.66073206 | 2.18907355 |
| WDR5 | 0.65944531 | 2.35630293 |
| UHRF1 | 0.65909965 | 1.8453363 |
| PRR3 | 0.6590454 | 1.68843525 |
| BRD8 | 0.65872366 | 2.1602328 |
| THAP4 | 0.65814523 | 1.89130513 |
| SPINDOC | 0.65786213 | 1.8192681 |
| LUC7L3 | 0.65773582 | 2.96189858 |
| CUX1 | 0.65643126 | 2.58847117 |
| POLR2B | 0.65636024 | 2.82306181 |
| POLE2 | 0.65588745 | 1.84373655 |
| KDM5B | 0.65467018 | 2.90260504 |
| DCAF12 | 0.65421289 | 1.60510855 |
| SBNO1 | 0.65369345 | 1.89522741 |
| MPHOSPH10 | 0.65326814 | 2.5651252 |
| FOXC1 | 0.65292364 | 1.54945308 |
| MOV10 | 0.6527572 | 2.22369278 |
| HCFC2 | 0.65249478 | 2.1388785 |
| OTUD5 | 0.65235752 | 1.70792921 |
| NCAPD3 | 0.65200159 | 2.29147796 |
| ZNF7 | 0.65172624 | 2.13535205 |
| CLK4 | 0.6510043 | 1.3938767 |
| WDHD1 | 0.65052702 | 1.5039531 |
| NCAPG2 | 0.65039458 | 2.00304731 |
| SOX5 | 0.6502285 | 3.81175833 |
| ECT2 | 0.64830083 | 2.32074154 |
| WDR36 | 0.64829631 | 2.40284129 |
| TSHZ3 | 0.64759531 | 2.96358584 |

|  |  |  |
| --- | --- | --- |
| CBX4 | 0.6472681 | 1.79374964 |
| ASXL1 | 0.64706244 | 4.11129474 |
| ZMYM3 | 0.64693966 | 1.56422201 |
| EPC2 | 0.64536787 | 2.82820035 |
| ARID4B | 0.64535077 | 2.52988409 |
| NR2F6 | 0.64493332 | 1.58809766 |
| BRCA2 | 0.64233375 | 1.44168905 |
| PHF8 | 0.64204038 | 2.98323152 |
| TNRC18 | 0.64141334 | 2.12679013 |
| ZBTB1 | 0.64073898 | 3.62408565 |
| WDR82 | 0.64046939 | 1.9478249 |
| ARID2 | 0.64020009 | 1.94617233 |
| RBM41 | 0.64001313 | 2.11808373 |
| EHMT1 | 0.63967523 | 2.39793155 |
| TRAF4 | 0.63717927 | 1.36666414 |
| CHD1L | 0.6370334 | 2.74077026 |
| NUFIP1 | 0.63682133 | 2.56798049 |
| ASF1A | 0.6366802 | 1.56570467 |
| DDB2 | 0.63636938 | 2.16380638 |
| NEIL3 | 0.63293046 | 2.0477278 |
| POLR2C | 0.63247813 | 1.44994247 |
| NRDE2 | 0.63244938 | 1.75296119 |
| ZNF219 | 0.63212481 | 1.80889504 |
| SP3 | 0.63171068 | 1.42631978 |
| BUD31 | 0.63089624 | 1.53619637 |
| HSPB1 | 0.6307409 | 2.48053127 |
| RPA2 | 0.62933949 | 1.48451381 |
| KLF5 | 0.62924139 | 1.59806491 |
| ZNF592 | 0.62779342 | 2.6364116 |
| ZDBF2 | 0.62694603 | 2.71202366 |
| CTR9 | 0.62662038 | 1.31383062 |
| PLAGL2 | 0.62611021 | 4.21915552 |
| LSM6 | 0.6251765 | 1.5500189 |
| RAD50 | 0.62503467 | 1.52840885 |
| MTBP | 0.62495184 | 1.52610072 |
| ZKSCAN2 | 0.62410588 | 1.33410606 |
| N4BP2L2 | 0.62337033 | 2.78461094 |
| TADA2A | 0.62295993 | 1.51283725 |
| ZSWIM3 | 0.62161514 | 1.6052431 |
| MGA | 0.62129976 | 1.55062094 |
| STK19 | 0.6211236 | 3.23777011 |
| PCGF2 | 0.62058615 | 1.90212584 |
| DNAJB6 | 0.62001321 | 1.48690976 |
| ZNF282 | 0.61971874 | 2.23032541 |
| PDCD4 | 0.61904144 | 1.31318282 |
| DDX20 | 0.61879511 | 1.45455749 |
| TBL1X | 0.6175591 | 1.66849175 |
| GATAD2A | 0.61748885 | 4.00958382 |

|  |  |  |
| --- | --- | --- |
| INTS1 | 0.61717543 | 1.71956682 |
| GLE1 | 0.61717414 | 1.34362515 |
| G2E3 | 0.61650064 | 2.3299716 |
| ERCC6 | 0.61629209 | 2.40242255 |
| SKIL | 0.61596873 | 1.5649599 |
| TAF9 | 0.61590297 | 1.39355854 |
| CHD4 | 0.61432811 | 3.27656648 |
| ARHGEF40 | 0.61402017 | 1.52491811 |
| FTO | 0.6134691 | 1.98560966 |
| KNL1 | 0.613087 | 1.79667545 |
| TSPYL2 | 0.6130387 | 1.56541828 |
| RSBN1L | 0.61275601 | 3.22516817 |
| TARDBP | 0.61137126 | 1.47202068 |
| ASXL2 | 0.60981087 | 2.48111871 |
| ZNF771 | 0.60906842 | 1.89518055 |
| PCGF1 | 0.60806351 | 2.81591796 |
| KMT2B | 0.60758498 | 2.69343359 |
| BICRAL | 0.60704108 | 1.84575388 |
| SRBD1 | 0.60680768 | 2.77372717 |
| LSM3 | 0.60674401 | 2.12556193 |
| RECQL4 | 0.60608753 | 1.44180413 |
| INTS3 | 0.60601531 | 2.46003071 |
| RBAK | 0.60515776 | 1.76665594 |
| MED27 | 0.60453804 | 2.28415907 |
| CCDC9B | 0.60395703 | 1.5592073 |
| LSM11 | 0.60278037 | 1.58785386 |
| DEPDC1 | 0.60256964 | 2.73697132 |
| ZNF497 | 0.60115091 | 1.93574034 |
| RBPJ | 0.60064117 | 1.76723066 |
| ZNF775 | 0.60026798 | 1.53514783 |
| GTF2H3 | 0.59960792 | 2.37791588 |
| UNC80 | 0.59938367 | 1.4653711 |
| ZNF48 | 0.59913923 | 2.32867157 |
| CHTF18 | 0.59848878 | 1.85918616 |
| CHEK2 | 0.59756483 | 1.83947287 |
| ZFHX4 | 0.59532989 | 2.08134789 |
| DNTTIP1 | 0.59506146 | 1.67691139 |
| KANSL1 | 0.59441798 | 1.66879669 |
| RNPS1 | 0.59321544 | 2.31859991 |
| TOPBP1 | 0.59243841 | 2.05067501 |
| TINF2 | 0.591846 | 2.18380903 |
| GTF2H2;GTF2I | 0.59094519 | 1.4525399 |
| POLL | 0.59049833 | 1.46511449 |
| CHD3 | 0.58991578 | 2.62714374 |
| KAT7 | 0.58830687 | 1.99128869 |
| TRRAP | 0.58816972 | 2.8014852 |
| FOXP1 | 0.58802988 | 2.6234993 |
| SNRPA1 | 0.5854084 | 2.27677895 |

|  |  |  |
| --- | --- | --- |
| RCOR3 | 0.58409639 | 1.77286575 |
| INCENP | 0.58402372 | 1.35222502 |
| FAM102B | 0.58318736 | 1.98810285 |
| TOPORS | 0.58131901 | 2.17201205 |
| KDM2B | 0.58108282 | 3.09570154 |
| POLR2L | 0.58029448 | 2.19454652 |
| FIZ1 | 0.5787453 | 1.96495476 |
| ATRX | 0.57871922 | 3.10495948 |
| CHAMP1 | 0.5783589 | 3.1346213 |
| NKRF | 0.57814812 | 2.40945879 |
| RERE | 0.57795147 | 1.83105362 |
| SATB2 | 0.57786348 | 1.57473489 |
| SKIV2L | 0.57730607 | 3.75966425 |
| RLIM | 0.57601651 | 1.93103049 |
| ZNF703 | 0.57590004 | 2.29727136 |
| KAT5 | 0.57443336 | 2.42893773 |
| HDAC1 | 0.5742065 | 2.01742091 |
| USP28 | 0.57415332 | 1.97354931 |
| LRRC41 | 0.57400954 | 2.83102446 |
| DOT1L | 0.57284311 | 2.47744278 |
| LCOR | 0.57227 | 2.6230985 |
| ZNF184 | 0.57225138 | 2.95340653 |
| DHX34 | 0.57171845 | 2.12184705 |
| PHF20 | 0.5710686 | 4.37215829 |
| ZFXH3 | 0.57071444 | 1.68147511 |
| NEIL2 | 0.57057849 | 1.78147744 |
| MED16 | 0.56916571 | 2.97207981 |
| POLR3C | 0.56713708 | 2.08626695 |
| SKI | 0.56698534 | 2.61696343 |
| AKAP17A | 0.566538 | 2.75813712 |
| RFC2 | 0.56627675 | 2.54838352 |
| APOBEC3F | 0.56621859 | 1.75789363 |
| PCGF3 | 0.56590396 | 3.12035757 |
| PWWP2A | 0.56440087 | 2.87604138 |
| RECQL | 0.5631792 | 1.57644395 |
| ZBTB21 | 0.56180402 | 1.90365726 |
| DNA2 | 0.5614578 | 1.76350261 |
| PHAX | 0.56087935 | 1.73391156 |
| ZNF280D | 0.56082055 | 1.9767846 |
| ZNF335 | 0.56056121 | 1.33626459 |
| ZNF189 | 0.56018187 | 2.40907756 |
| LIG3 | 0.5596203 | 1.54374522 |
| SF3A3 | 0.55909897 | 1.44556648 |
| MED24 | 0.55821643 | 1.54427778 |
| YY1AP1 | 0.55483337 | 1.43812141 |
| BARD1 | 0.55382212 | 2.12778303 |
| NAV2 | 0.55371555 | 1.35796569 |
| SRSF8 | 0.55310167 | 1.92199302 |

|  |  |  |
| --- | --- | --- |
| RBM19 | 0.55213105 | 2.22388879 |
| EPC1 | 0.55203024 | 1.89363562 |
| FANCD2 | 0.55079249 | 1.93544469 |
| DUSP14 | 0.55037742 | 2.58132461 |
| ERCC5 | 0.54936183 | 2.94391437 |
| PIAS1 | 0.5492957 | 2.25119799 |
| MLF2 | 0.54852101 | 2.04906573 |
| ZKSCAN3 | 0.54818065 | 1.54694342 |
| IDI1 | 0.54763675 | 1.67996062 |
| CHD6 | 0.54673269 | 1.52395431 |
| NYNRIN | 0.54511893 | 1.7466378 |
| TADA2B | 0.54432848 | 3.30997858 |
| SKP2 | 0.54425181 | 2.09570235 |
| EHMT2 | 0.54423703 | 2.30384628 |
| PHF12 | 0.54418307 | 2.58504597 |
| C19orf44 | 0.54382669 | 1.63341278 |
| POLR2G | 0.54374357 | 1.78612008 |
| MDC1 | 0.54160209 | 1.43993814 |
| POLD1 | 0.54154947 | 1.35647466 |
| CENPT | 0.54036474 | 1.79510265 |
| HCFC1 | 0.5400312 | 4.04886157 |
| ZNF646 | 0.53837414 | 2.83791804 |
| CIZ1 | 0.53618948 | 1.44625729 |
| UBR7 | 0.53603923 | 2.99505328 |
| PIAS2 | 0.53521171 | 1.72511997 |
| LRRC42 | 0.53508868 | 1.36688336 |
| ZNF35 | 0.53453651 | 1.69124326 |
| UTP14A | 0.53311033 | 1.95771515 |
| KMT2A | 0.53302459 | 1.74676228 |
| NACC2 | 0.53298256 | 1.74469226 |
| HDGFL2 | 0.53278559 | 2.08662721 |
| RNASEH2C | 0.53107453 | 1.65935917 |
| MYBL2 | 0.53104912 | 1.92634633 |
| BAZ1A | 0.53064788 | 1.73012451 |
| TET1 | 0.53039888 | 1.89162501 |
| FOXJ3 | 0.52814433 | 1.48400795 |
| ERCC4 | 0.52779315 | 2.28181153 |
| S100A7 | 0.52767414 | 1.84769178 |
| KANSL3 | 0.52763051 | 1.77748761 |
| GTF3C2 | 0.52600187 | 1.77182169 |
| NCOA1 | 0.52568937 | 1.94757714 |
| NFKBIL1 | 0.52551525 | 1.98704104 |
| HOXA13 | 0.52510246 | 1.40456822 |
| ZGPAT | 0.52421531 | 1.96240778 |
| ZNF408 | 0.52323832 | 1.63516984 |
| DDX50 | 0.52312175 | 1.52911355 |
| PHF3 | 0.52281965 | 2.47922445 |
| ZMYND8 | 0.52253176 | 2.08810684 |

|  |  |  |
| --- | --- | --- |
| HAT1 | 0.52137436 | 1.8687017 |
| CCDC106 | 0.52086078 | 1.33592058 |
| ZZZ3 | 0.52074803 | 2.75474854 |
| ZNF385A | 0.51921772 | 1.43080841 |
| ELL | 0.51857871 | 1.8853661 |
| CSTF2T | 0.51834941 | 1.70056323 |
| FOXG1 | 0.51820976 | 2.38638366 |
| HIC2 | 0.51680198 | 2.13531016 |
| HLTF | 0.51666197 | 3.01577297 |
| SPRTN | 0.51641352 | 1.39056377 |
| ZNF770 | 0.51614218 | 1.35914402 |
| MED23 | 0.51602882 | 2.05822826 |
| PHC1 | 0.51488818 | 2.54249031 |
| SMARCA5 | 0.51481595 | 2.00370736 |
| RRP9 | 0.51282191 | 2.54107337 |
| PSMB4 | 0.51264385 | 1.61915835 |
| UTP18 | 0.51213234 | 4.03600859 |
| ARMCX5 | 0.51021463 | 1.40683323 |
| KDM4C | 0.50995792 | 1.86210282 |
| ZHX1 | 0.50930463 | 2.7111717 |
| UTP20 | 0.50922117 | 2.58564029 |
| SLFN5 | 0.50885451 | 1.49294126 |
| RFX7 | 0.50675693 | 3.12815015 |
| MNAT1 | 0.50557869 | 2.47740489 |
| JADE1 | 0.50342857 | 2.60183231 |
| RNGTT | 0.50338519 | 2.50948353 |
| RAD54L2 | 0.5032524 | 2.6573114 |
| KAT8 | 0.50299414 | 1.47931372 |

Log2FoldChange was calculated by two independent experimental replicates. Ribosomal protein
