## Supplementary Table 4 for "RNA m6A mediates *de novo* DNA methylation and heritable epigenetic silencing in human cells"

**Table S4. Positive hits identified by APEX2-proximity labeling (C-term dCas9-KRAB-APEX2 24 h**

| **Genes** | **Log2(Fold Change)** | **Negative log10 (p-value)** |
| --- | --- | --- |
| ZNF10 | 6.62 | 3.19 |
| NUDT21 | 1.61 | 3.55 |
| TRIM28 | 1.53 | 3.44 |
| DNMT3A | 1.47 | 3.33 |
| HIC2 | 1.33 | 2.66 |
| CPSF7 | 1.32 | 3.64 |
| POU3F2 | 1.30 | 3.24 |
| DNAJB1 | 1.29 | 2.84 |
| GNG12 | 1.29 | 2.65 |
| FUBP1 | 1.28 | 4.18 |
| RP9 | 1.27 | 2.27 |
| CIR1 | 1.27 | 2.40 |
| CPSF6 | 1.26 | 3.71 |
| TRIM27 | 1.26 | 3.20 |
| ZKSCAN4 | 1.23 | 2.67 |
| RIF1 | 1.22 | 2.81 |
| SMARCAD1 | 1.21 | 1.95 |
| TAX1BP1 | 1.20 | 2.57 |
| NFYC | 1.19 | 4.07 |
| CCNYL1 | 1.19 | 2.17 |
| RFX1 | 1.18 | 2.07 |
| TRIM24 | 1.17 | 3.34 |
| TP53BP1 | 1.17 | 2.16 |
| ZWINT | 1.17 | 2.82 |
| LTV1 | 1.17 | 1.97 |
| EFR3A | 1.16 | 2.54 |
| ZBTB21 | 1.15 | 3.25 |
| GNAI2 | 1.15 | 2.18 |
| NACC1 | 1.15 | 3.98 |
| LZTS2 | 1.13 | 2.47 |
| DDX41 | 1.13 | 2.76 |
| GNB2 | 1.12 | 2.26 |
| EWSR1 | 1.12 | 2.08 |
| MACROH2A1 | 1.11 | 1.62 |
| WTAP | 1.11 | 2.19 |
| ULBP3 | 1.11 | 1.85 |
| RSRC2 | 1.10 | 3.81 |
| ULBP2 | 1.10 | 2.41 |
| ARID1A | 1.09 | 2.99 |
| LUC7L | 1.09 | 2.95 |
| CASZ1 | 1.09 | 3.44 |
| TMF1 | 1.07 | 2.91 |
| ZEB1 | 1.07 | 1.61 |
| TRAFD1 | 1.07 | 4.46 |
| SUMO1 | 1.06 | 1.98 |
| UBXN4 | 1.06 | 2.55 |
| YES1 | 1.06 | 2.37 |
| H3C13 | 1.04 | 1.68 |
| TRPS1 | 1.04 | 2.51 |
| BEND3 | 1.03 | 4.13 |
| ZNF24 | 1.03 | 2.91 |
| CPSF4 | 1.02 | 3.83 |
| POLD1 | 1.02 | 3.28 |
| YLPM1 | 1.02 | 2.81 |
| DDX20 | 1.02 | 2.54 |
| NR2F1 | 1.02 | 2.77 |
| ZBTB33 | 1.01 | 3.01 |
| ARID3B | 1.01 | 3.27 |
| TASOR2 | 0.99 | 2.68 |
| MSH2 | 0.99 | 2.87 |
| PRKDC | 0.98 | 2.96 |
| INTS8 | 0.98 | 2.70 |
| PCGF6 | 0.97 | 3.01 |
| YY1 | 0.97 | 1.85 |
| DNAJC7 | 0.97 | 2.14 |
| RBM25 | 0.97 | 2.19 |
| ZSCAN29 | 0.97 | 2.54 |
| DDI2 | 0.96 | 2.30 |
| GNAI1 | 0.96 | 2.27 |
| BTAF1 | 0.96 | 1.75 |
| MBIP | 0.96 | 2.73 |
| BAG6 | 0.96 | 3.07 |
| KDM6A | 0.95 | 1.92 |
| TNKS1BP1 | 0.95 | 2.34 |
| CEP95 | 0.95 | 2.29 |
| GEMIN4 | 0.95 | 2.13 |
| LYN | 0.95 | 2.30 |
| MIA2 | 0.95 | 1.92 |
| CCHCR1 | 0.95 | 2.15 |
| HIRA | 0.94 | 3.84 |
| SAP130 | 0.94 | 1.32 |
| TAF6L | 0.94 | 3.17 |
| GNAI3 | 0.94 | 1.97 |
| PUF60 | 0.93 | 2.81 |
| ZNF148 | 0.93 | 3.25 |
| ZZZ3 | 0.92 | 3.31 |
| ZMYM2 | 0.92 | 2.42 |
| GTF3C3 | 0.92 | 2.15 |
| RAB11FIP1 | 0.92 | 3.46 |
| MGA | 0.92 | 4.01 |
| BTBD1 | 0.92 | 2.67 |
| PRDM16 | 0.91 | 3.56 |
| DNAJB6 | 0.91 | 2.41 |
| POLR3D | 0.91 | 2.85 |
| PPP1R12C | 0.91 | 2.46 |
| HMBOX1 | 0.91 | 1.66 |
| KIAA0753 | 0.91 | 2.63 |
| CUX1 | 0.91 | 3.12 |
| MSH6 | 0.90 | 2.36 |
| VIRMA | 0.90 | 2.73 |
| ATF6B | 0.90 | 1.67 |
| WDR33 | 0.90 | 2.80 |
| JAK1 | 0.90 | 2.40 |
| EPC2 | 0.90 | 2.64 |
| HDAC2 | 0.89 | 2.17 |
| HDAC1 | 0.89 | 2.60 |
| AKAP12 | 0.89 | 2.30 |
| ARID3A | 0.89 | 3.06 |
| TRIM33 | 0.89 | 2.18 |
| KIAA1671 | 0.88 | 2.73 |
| DVL3 | 0.88 | 2.70 |
| PRPF6 | 0.88 | 3.08 |
| MAGED1 | 0.88 | 2.53 |
| MAGED4B | 0.88 | 3.30 |
| ARID1B | 0.87 | 1.86 |
| ZC3H13 | 0.87 | 2.62 |
| SRRM2 | 0.87 | 2.12 |
| SMARCE1 | 0.86 | 2.18 |
| RCOR1 | 0.86 | 3.49 |
| NKAPD1 | 0.86 | 2.38 |
| KAT14 | 0.86 | 2.98 |
| BAG5 | 0.86 | 2.49 |
| SF3B6 | 0.86 | 2.89 |
| TBX3 | 0.86 | 1.93 |
| ZBTB10 | 0.86 | 2.88 |
| ATG9A | 0.85 | 2.25 |
| NPLOC4 | 0.85 | 2.18 |
| NR2C1 | 0.85 | 2.32 |
| PRPF40A | 0.85 | 3.30 |
| SMARCA4 | 0.84 | 2.58 |
| PHF21A | 0.84 | 1.77 |
| DEPDC1B | 0.84 | 2.06 |
| TRIM37 | 0.84 | 3.21 |
| PAXIP1 | 0.84 | 1.99 |
| CSTF3 | 0.84 | 2.06 |
| HELLS | 0.84 | 1.89 |
| ZEB2 | 0.84 | 5.00 |
| MYBL2 | 0.84 | 3.50 |
| BCOR | 0.84 | 3.64 |
| SUPT6H | 0.83 | 3.37 |
| POLR2A | 0.83 | 3.07 |
| CASP8AP2 | 0.83 | 2.32 |
| SMARCA2 | 0.83 | 3.20 |
| ZNF609 | 0.82 | 4.20 |
| DLG5 | 0.82 | 2.56 |
| DMAP1 | 0.82 | 2.90 |
| ODF2L | 0.82 | 2.38 |
| CYP51A1 | 0.82 | 2.12 |
| INTS2 | 0.82 | 2.30 |
| ELF1 | 0.82 | 2.50 |
| NBR1 | 0.82 | 3.22 |
| MORC3 | 0.81 | 2.54 |
| SIN3A | 0.81 | 2.76 |
| RPRD2 | 0.81 | 2.98 |
| NT5DC2 | 0.81 | 2.37 |
| AHNAK2 | 0.81 | 2.59 |
| GNA11 | 0.81 | 1.88 |
| EP400 | 0.80 | 2.85 |
| PTPN1 | 0.80 | 4.26 |
| CNKSR3 | 0.80 | 1.60 |
| ATF7IP | 0.80 | 2.08 |
| ZNF593 | 0.80 | 2.07 |
| RPA1 | 0.80 | 3.69 |
| CCDC120 | 0.80 | 2.40 |
| MEF2D | 0.80 | 3.03 |
| MPHOSPH9 | 0.80 | 1.73 |
| SETD2 | 0.79 | 2.98 |
| ZNF516 | 0.79 | 1.58 |
| ZNF318 | 0.79 | 2.98 |
| NOTCH2 | 0.79 | 1.80 |
| RAB18 | 0.79 | 2.16 |
| KDM1A | 0.78 | 2.17 |
| MACROH2A2 | 0.78 | 1.32 |
| ZNF638 | 0.78 | 2.06 |
| FIP1L1 | 0.78 | 2.76 |
| CCT7 | 0.78 | 2.59 |
| OSER1 | 0.78 | 1.53 |
| L3MBTL2 | 0.77 | 2.73 |
| KIDINS220 | 0.77 | 2.21 |
| CECR2 | 0.77 | 2.32 |
| BIRC6 | 0.77 | 2.58 |
| GTF2I | 0.77 | 2.88 |
| ORC3 | 0.77 | 2.61 |
| DNAJC1 | 0.77 | 1.42 |
| MORC4 | 0.76 | 2.68 |
| SETD1A | 0.76 | 1.86 |
| ZNF16 | 0.76 | 2.62 |
| ZKSCAN8 | 0.76 | 2.82 |
| RINT1 | 0.76 | 2.68 |
| BACH1 | 0.76 | 2.17 |
| PHC3 | 0.76 | 2.02 |
| SF3A3 | 0.76 | 2.37 |
| TBC1D7 | 0.76 | 1.50 |
| TP53 | 0.75 | 2.30 |
| ZIC2 | 0.75 | 1.87 |
| RBM17 | 0.75 | 2.76 |
| RCOR3 | 0.75 | 2.47 |
| ZHX2 | 0.75 | 1.39 |
| SMCHD1 | 0.75 | 2.85 |
| NELFCD | 0.75 | 1.85 |
| JUN | 0.75 | 3.24 |
| TRRAP | 0.75 | 2.18 |
| PTPN14 | 0.75 | 1.78 |
| CSRP2 | 0.75 | 2.27 |
| UCK2 | 0.75 | 2.58 |
| RB1CC1 | 0.75 | 3.39 |
| RFX7 | 0.74 | 2.33 |
| DDX46 | 0.74 | 1.91 |
| RFWD3 | 0.74 | 2.04 |
| HUWE1 | 0.74 | 2.18 |
| TCAF1 | 0.74 | 2.07 |
| C15orf39 | 0.74 | 1.89 |
| LUC7L3 | 0.74 | 2.30 |
| WIZ | 0.73 | 3.55 |
| MECOM | 0.73 | 1.68 |
| ZNF655 | 0.73 | 2.43 |
| ZNF629 | 0.73 | 2.88 |
| RAD21 | 0.73 | 1.51 |
| ETV6 | 0.73 | 2.21 |
| NEK2 | 0.73 | 2.64 |
| SNW1 | 0.73 | 1.95 |
| HOMEZ | 0.73 | 1.73 |
| EEPD1 | 0.73 | 1.54 |
| FAM120B | 0.73 | 2.86 |
| HERC2 | 0.73 | 2.83 |
| POLG | 0.73 | 1.77 |
| TCEA1 | 0.73 | 2.88 |
| POLR2B | 0.73 | 2.12 |
| RECQL | 0.73 | 1.87 |
| LRRC49 | 0.73 | 2.17 |
| CSTF2 | 0.73 | 3.36 |
| NCAPG2 | 0.72 | 2.31 |
| DPF2 | 0.72 | 2.96 |
| KMT2D | 0.72 | 1.77 |
| SMARCC1 | 0.72 | 2.10 |
| ZMYM3 | 0.72 | 1.40 |
| DNMT1 | 0.72 | 2.64 |
| SMC2 | 0.72 | 2.29 |
| EXO1 | 0.72 | 2.18 |
| EXOC7 | 0.72 | 2.33 |
| HIVEP2 | 0.72 | 1.65 |
| VRK2 | 0.72 | 4.50 |
| SUPT5H | 0.72 | 2.76 |
| SCFD1 | 0.72 | 1.59 |
| DIS3 | 0.72 | 1.59 |
| FOXO1 | 0.72 | 3.09 |
| TAF9 | 0.72 | 2.49 |
| PHRF1 | 0.72 | 2.31 |
| CDK9 | 0.72 | 3.53 |
| KDM2B | 0.71 | 2.99 |
| CREBBP | 0.71 | 2.27 |
| ZBED4 | 0.71 | 1.99 |
| ZNF768 | 0.71 | 2.84 |
| PIBF1 | 0.71 | 2.25 |
| C17orf80 | 0.71 | 1.70 |
| RAB11FIP5 | 0.71 | 2.19 |
| L3MBTL3 | 0.71 | 2.81 |
| PROSER1 | 0.71 | 1.96 |
| UBE3C | 0.71 | 2.87 |
| TADA1 | 0.71 | 2.66 |
| RAD54L2 | 0.71 | 4.28 |
| TRIP11 | 0.71 | 2.22 |
| CEP170 | 0.71 | 2.10 |
| MTBP | 0.71 | 2.23 |
| SPTLC2 | 0.71 | 1.61 |
| UFSP2 | 0.71 | 2.41 |
| CLK3 | 0.71 | 1.94 |
| KMT2C | 0.70 | 2.95 |
| ZFHX4 | 0.70 | 2.50 |
| DOCK7 | 0.70 | 2.18 |
| SUPT3H | 0.70 | 2.96 |
| TLK1 | 0.70 | 1.83 |
| MCM2 | 0.70 | 2.33 |
| PSMG1 | 0.70 | 1.70 |
| SRSF11 | 0.70 | 3.47 |
| XRCC3 | 0.70 | 2.80 |
| UBE2I | 0.70 | 2.19 |
| CREB3L2 | 0.70 | 2.30 |
| KCTD3 | 0.70 | 2.53 |
| POLR2H | 0.70 | 2.44 |
| POLA1 | 0.70 | 1.62 |
| KIAA1586 | 0.70 | 2.21 |
| CABIN1 | 0.69 | 2.31 |
| LUC7L2 | 0.69 | 2.28 |
| WDHD1 | 0.69 | 2.67 |
| LRCH2 | 0.69 | 2.48 |
| FANCD2 | 0.69 | 1.80 |
| SUPT7L | 0.69 | 2.39 |
| TLK2 | 0.69 | 2.71 |
| WDR6 | 0.69 | 1.60 |
| MED26 | 0.69 | 2.43 |
| NUP93 | 0.69 | 1.95 |
| SMC1A | 0.69 | 3.01 |
| TADA2A | 0.69 | 2.57 |
| SCAF4 | 0.69 | 1.99 |
| U2SURP | 0.69 | 2.45 |
| MAGED2 | 0.69 | 3.18 |
| FOXK2 | 0.69 | 3.38 |
| RBM26 | 0.69 | 2.49 |
| SATB2 | 0.69 | 2.80 |
| ZNF687 | 0.68 | 2.52 |
| EPS15 | 0.68 | 2.12 |
| LMO7 | 0.68 | 3.18 |
| SF3B1 | 0.68 | 1.94 |
| CDC25C | 0.68 | 2.25 |
| CENPH | 0.68 | 4.25 |
| DDX23 | 0.68 | 2.04 |
| RECQL4 | 0.68 | 1.42 |
| TAF9B | 0.68 | 2.34 |
| PBX1 | 0.67 | 3.69 |
| PIAS1 | 0.67 | 3.03 |
| SHKBP1 | 0.67 | 2.83 |
| INTS6 | 0.67 | 2.77 |
| MCMBP | 0.67 | 1.96 |
| PSMD1 | 0.67 | 1.77 |
| PCF11 | 0.67 | 2.76 |
| ARID4B | 0.67 | 2.64 |
| TP53BP2 | 0.67 | 2.55 |
| DDRKG1 | 0.67 | 1.50 |
| SFPQ | 0.67 | 2.07 |
| ZMYM4 | 0.67 | 3.32 |
| CHD8 | 0.66 | 2.12 |
| KDM2A | 0.66 | 2.45 |
| ZNF217 | 0.66 | 2.22 |
| ZMIZ2 | 0.66 | 1.70 |
| TBC1D15 | 0.66 | 2.41 |
| ERCC2 | 0.66 | 2.09 |
| TAF5L | 0.66 | 2.91 |
| GTF3C1 | 0.66 | 2.96 |
| UBAC2 | 0.66 | 1.38 |
| DCAF1 | 0.66 | 2.39 |
| SNAPC4 | 0.66 | 2.03 |
| WDR48 | 0.66 | 1.36 |
| ATRIP | 0.66 | 2.12 |
| WRN | 0.66 | 2.84 |
| LENG8 | 0.66 | 2.73 |
| BRIP1 | 0.66 | 2.30 |
| KAT2A | 0.66 | 2.85 |
| SH3D19 | 0.66 | 3.07 |
| ZNF219 | 0.65 | 3.32 |
| MED13 | 0.65 | 1.88 |
| DDX42 | 0.65 | 2.62 |
| PRR12 | 0.65 | 1.67 |
| CD2BP2 | 0.65 | 4.02 |
| TELO2 | 0.65 | 1.54 |
| FHOD1 | 0.65 | 1.49 |
| ENAH | 0.65 | 2.05 |
| SAFB | 0.65 | 3.29 |
| INTS1 | 0.65 | 2.02 |
| GANAB | 0.65 | 2.14 |
| HLTF | 0.65 | 3.38 |
| SMARCC2 | 0.64 | 2.72 |
| ZNF579 | 0.64 | 3.49 |
| DCAF7 | 0.64 | 2.76 |
| CPSF3 | 0.64 | 1.70 |
| CPSF2 | 0.64 | 3.05 |
| DENR | 0.64 | 2.59 |
| EXOC6 | 0.64 | 1.80 |
| NELFA | 0.64 | 1.93 |
| RAD23B | 0.64 | 2.64 |
| BRD4 | 0.63 | 2.86 |
| TERF2IP | 0.63 | 2.37 |
| TPI1 | 0.63 | 2.01 |
| RHEB | 0.63 | 3.37 |
| RBBP7 | 0.63 | 2.01 |
| CLK2 | 0.63 | 2.36 |
| XAB2 | 0.63 | 2.03 |
| IRF2BP1 | 0.63 | 2.31 |
| USP32 | 0.63 | 2.07 |
| ZFYVE16 | 0.63 | 1.83 |
| SALL2 | 0.63 | 1.77 |
| PRPF8 | 0.63 | 1.89 |
| WDR62 | 0.63 | 2.41 |
| MARCHF7 | 0.63 | 1.96 |
| INTS12 | 0.63 | 3.62 |
| PTK7 | 0.63 | 1.53 |
| SHPRH | 0.63 | 4.36 |
| ZNHIT6 | 0.63 | 2.38 |
| KDM4A | 0.62 | 3.28 |
| FOXC1 | 0.62 | 2.67 |
| ZNF771 | 0.62 | 2.81 |
| ZHX3 | 0.62 | 4.43 |
| NUP133 | 0.62 | 2.30 |
| BAG2 | 0.62 | 1.57 |
| SUDS3 | 0.62 | 3.43 |
| EDRF1 | 0.62 | 2.14 |
| LRCH1 | 0.62 | 2.09 |
| TAF4 | 0.62 | 2.42 |
| VPS4A | 0.62 | 2.27 |
| USP5 | 0.62 | 1.84 |
| ZKSCAN1 | 0.62 | 2.44 |
| SLU7 | 0.61 | 2.18 |
| MED4 | 0.61 | 2.85 |
| RICTOR | 0.61 | 1.84 |
| E2F6 | 0.61 | 2.15 |
| HOXC9 | 0.61 | 2.89 |
| GIT2 | 0.61 | 1.45 |
| YEATS4 | 0.61 | 2.42 |
| BUB3 | 0.61 | 1.71 |
| POLE | 0.61 | 2.30 |
| POLR2C | 0.61 | 1.80 |
| UBE3A | 0.61 | 2.37 |
| INTS7 | 0.61 | 1.72 |
| SENP1 | 0.61 | 1.95 |
| CDCA7L | 0.60 | 2.94 |
| MYSM1 | 0.60 | 1.71 |
| CEP97 | 0.60 | 1.78 |
| HOXA10 | 0.60 | 1.72 |
| AUP1 | 0.60 | 2.21 |
| ERCC6L | 0.60 | 2.70 |
| BCL11A | 0.60 | 2.37 |
| CEP135 | 0.60 | 1.97 |
| CEP112 | 0.60 | 1.58 |
| SLX4 | 0.60 | 2.18 |
| NUP88 | 0.60 | 2.02 |
| WNK1 | 0.60 | 1.89 |
| TP73 | 0.60 | 1.55 |
| KANSL1 | 0.60 | 3.85 |
| PPP1R10 | 0.60 | 1.77 |
| KANSL3 | 0.59 | 1.69 |
| SF3A1 | 0.59 | 3.07 |
| BCLAF1 | 0.59 | 1.68 |
| MCM7 | 0.59 | 2.79 |
| NUP85 | 0.59 | 1.82 |
| RB1 | 0.59 | 2.19 |
| CENPT | 0.59 | 2.49 |
| YJU2 | 0.59 | 1.50 |
| NUP160 | 0.59 | 1.89 |
| ZHX1 | 0.59 | 3.51 |
| NUP214 | 0.59 | 1.94 |
| NEURL4 | 0.58 | 3.75 |
| WDR37 | 0.58 | 2.86 |
| PPP5C | 0.58 | 2.50 |
| KIAA0586 | 0.58 | 2.96 |
| ATF6 | 0.58 | 1.89 |
| PBX2 | 0.58 | 1.96 |
| DNAJA1 | 0.58 | 2.45 |
| RBM10 | 0.58 | 2.14 |
| MED23 | 0.58 | 1.77 |
| ZC3H14 | 0.58 | 2.37 |
| MED1 | 0.57 | 1.79 |
| CCDC47 | 0.57 | 3.34 |
| SUGP2 | 0.57 | 2.89 |
| TAF15 | 0.57 | 2.34 |
| TCERG1 | 0.57 | 1.50 |
| MARK1 | 0.57 | 2.21 |
| RRM1 | 0.57 | 2.67 |
| GCC2 | 0.57 | 1.45 |
| VRK3 | 0.57 | 2.69 |
| RBM14 | 0.57 | 3.48 |
| LRRC59 | 0.57 | 2.89 |
| RNF40 | 0.57 | 2.55 |
| CEP78 | 0.57 | 2.54 |
| ERICH5 | 0.57 | 1.49 |
| GATA4 | 0.57 | 2.49 |
| WDR18 | 0.57 | 3.39 |
| DCAF8 | 0.56 | 2.48 |
| SAP30BP | 0.56 | 2.43 |
| DDX39A | 0.56 | 3.78 |
| THOC2 | 0.56 | 2.33 |
| HCFC1 | 0.56 | 2.37 |
| CHEK1 | 0.56 | 2.09 |
| NCOR1 | 0.56 | 2.90 |
| DEPDC1 | 0.56 | 2.87 |
| WDR41 | 0.56 | 1.41 |
| EPC1 | 0.56 | 2.35 |
| ZSCAN26 | 0.56 | 2.19 |
| UFD1 | 0.56 | 2.21 |
| CSTF2T | 0.56 | 2.33 |
| TADA3 | 0.56 | 3.04 |
| CDK5RAP3 | 0.56 | 2.22 |
| DOCK11 | 0.56 | 2.15 |
| BAIAP2L1 | 0.56 | 2.59 |
| PSAT1 | 0.56 | 2.17 |
| FOXP1 | 0.56 | 2.81 |
| ESS2 | 0.56 | 3.06 |
| SRCAP | 0.56 | 2.57 |
| RADX | 0.56 | 1.95 |
| TDP2 | 0.55 | 2.01 |
| EEF1E1 | 0.55 | 1.31 |
| PDXDC1 | 0.55 | 1.84 |
| ZBTB40 | 0.55 | 2.79 |
| SP2 | 0.55 | 2.06 |
| CHTF18 | 0.55 | 1.78 |
| CCNT1 | 0.55 | 2.22 |
| SALL1 | 0.55 | 3.61 |
| NTHL1 | 0.55 | 2.57 |
| PRIM1 | 0.55 | 3.22 |
| TJP2 | 0.55 | 2.43 |
| PRKAA2 | 0.55 | 3.00 |
| RMC1 | 0.55 | 1.51 |
| ZFYVE9 | 0.55 | 1.37 |
| EDC4 | 0.55 | 1.90 |
| RFX2 | 0.55 | 2.25 |
| LRIF1 | 0.55 | 2.41 |
| RBM22 | 0.54 | 2.55 |
| TFAP4 | 0.54 | 1.87 |
| ARVCF | 0.54 | 1.54 |
| LIG3 | 0.54 | 2.35 |
| VCPIP1 | 0.54 | 2.04 |
| SLK | 0.54 | 2.31 |
| CIZ1 | 0.54 | 2.72 |
| NCOR2 | 0.54 | 2.13 |
| NUP205 | 0.54 | 1.45 |
| CDK1 | 0.54 | 2.11 |
| BRD8 | 0.54 | 1.59 |
| SF1 | 0.54 | 2.64 |
| NFAT5 | 0.54 | 1.74 |
| SCAF11 | 0.54 | 2.53 |
| FLYWCH1 | 0.54 | 1.89 |
| SMU1 | 0.54 | 2.39 |
| NELFE | 0.54 | 1.88 |
| NHSL1 | 0.54 | 1.89 |
| NUP107 | 0.54 | 1.71 |
| BBS7 | 0.54 | 1.77 |
| SAFB2 | 0.53 | 2.97 |
| SF3A2 | 0.53 | 1.83 |
| PEX1 | 0.53 | 2.06 |
| RTF2 | 0.53 | 1.61 |
| MTA3 | 0.53 | 2.02 |
| IRS2 | 0.53 | 1.61 |
| NR2F6 | 0.53 | 2.08 |
| DMWD | 0.53 | 2.14 |
| CALCOCO2 | 0.53 | 2.02 |
| SMARCAL1 | 0.53 | 3.85 |
| MED12 | 0.53 | 2.62 |
| ARGLU1 | 0.53 | 1.83 |
| MCM8 | 0.53 | 3.59 |
| ARMC5 | 0.53 | 1.64 |
| UBA6 | 0.53 | 1.98 |
| HEATR1 | 0.53 | 2.45 |
| MCM9 | 0.53 | 1.85 |
| SCAF8 | 0.53 | 1.95 |
| FBXO22 | 0.53 | 2.16 |
| RSPRY1 | 0.53 | 1.35 |
| DACH1 | 0.53 | 2.00 |
| NELFB | 0.53 | 1.44 |
| ATG2B | 0.53 | 1.86 |
| XRCC1 | 0.53 | 3.09 |
| SMARCD3 | 0.53 | 2.54 |
| CDR2 | 0.53 | 2.10 |
| QRICH1 | 0.53 | 1.97 |
| RBM12 | 0.52 | 4.29 |
| UBE4B | 0.52 | 2.04 |
| MAP7D1 | 0.52 | 2.43 |
| HERC1 | 0.52 | 1.95 |
| TAF5 | 0.52 | 2.13 |
| SFSWAP | 0.52 | 2.47 |
| DDX28 | 0.52 | 2.74 |
| PPP6R3 | 0.52 | 2.18 |
| STK38L | 0.52 | 2.45 |
| GATA2 | 0.52 | 1.59 |
| PPP2R3C | 0.52 | 1.34 |
| KAT8 | 0.52 | 2.26 |
| NUP50 | 0.52 | 3.04 |
| RPRD1A | 0.52 | 2.33 |
| TASOR | 0.52 | 1.42 |
| TANC2 | 0.52 | 1.80 |
| FAM83B | 0.51 | 1.91 |
| KLF12 | 0.51 | 2.20 |
| DHX16 | 0.51 | 2.09 |
| TFG | 0.51 | 1.54 |
| CDR2L | 0.51 | 1.92 |
| NFIC | 0.51 | 2.51 |
| TOPBP1 | 0.51 | 1.93 |
| NDC80 | 0.51 | 2.05 |
| GATAD2A | 0.51 | 1.92 |
| DOCK9 | 0.51 | 2.25 |
| PCNA | 0.51 | 3.02 |
| DHX35 | 0.50 | 2.84 |
| CEP164 | 0.50 | 4.43 |
| USP9X | 0.50 | 1.90 |
| DOCK6 | 0.50 | 1.63 |
| POLR3E | 0.50 | 3.47 |
| THRAP3 | 0.50 | 1.34 |
| EXOC5 | 0.50 | 1.70 |
| EXOC4 | 0.50 | 2.17 |
| NEK9 | 0.50 | 1.56 |
| HIVEP1 | 0.50 | 2.87 |

Log2FoldChange was calculated by two independent experimental replicates. Ribosomal protein
