## Supplementary Table 5 for "RNA m6A mediates *de novo* DNA methylation and heritable epigenetic silencing in human cells"

**Table S5. Positive hits identified by APEX2-proximity labeling (N-term APEX2-KRAB-dCas9 at 48 hour**

| **Genes** | **Log2(Fold Chang Negative log10 (p-value)** |  |
| --- | --- | --- |
| ZNF10 | 9.231645269 | 3.875162391 |
| RESF1 | 1.973599905 | 1.3697017 |
| U2AF2 | 1.971612017 | 1.33479325 |
| PRPF19 | 1.81986818 | 1.304489089 |
| ZNF7 | 1.516900069 | 1.432730309 |
| SMARCAD1 | 1.476576252 | 1.432138526 |
| CCNL1 | 1.461126739 | 2.180645545 |
| MTREX | 1.359408426 | 2.232273848 |
| NOL10 | 1.357282598 | 1.411601005 |
| ZMYM6 | 1.356636808 | 1.344149377 |
| SIN3A | 1.338940125 | 1.369727193 |
| MCM6 | 1.329426539 | 1.919009561 |
| AATF | 1.311728489 | 1.467179949 |
| ZNF711 | 1.310247762 | 3.550208741 |
| H1-5 | 1.297129343 | 1.335334444 |
| RFX2 | 1.2830109 | 2.30720729 |
| CECR2 | 1.260370721 | 2.110533817 |
| PALB2 | 1.250110159 | 1.409742103 |
| KMT2C | 1.243331022 | 1.58271081 |
| RBM14 | 1.209841132 | 2.488476701 |
| TRIM24 | 1.205716114 | 1.601193449 |
| RSRC1 | 1.199198699 | 1.303381944 |
| HIVEP1 | 1.155919227 | 1.338439711 |
| GTPBP10 | 1.142220129 | 1.700544025 |
| ZFP91 | 1.137554854 | 1.512600563 |
| PRDM10 | 1.125766831 | 1.565745406 |
| THOC6 | 1.124082424 | 1.593792037 |
| MED13 | 1.12237701 | 2.456803766 |
| PCID2 | 1.114136236 | 1.487627986 |
| SNU13 | 1.102495115 | 1.397654588 |
| XRCC4 | 1.100951929 | 1.953553188 |
| LEO1 | 1.09616476 | 1.329720743 |
| SMARCD3 | 1.091846377 | 2.011246762 |
| CPSF2 | 1.06907964 | 1.447311675 |
| WDR82 | 1.053257125 | 1.482827879 |
| CTR9 | 1.050741301 | 1.857100619 |
| PRAME | 1.05045428 | 1.562544469 |
| SF3A2 | 1.050392736 | 1.32026449 |
| TCERG1 | 1.046107643 | 2.066906961 |
| BCL9L | 1.037465697 | 1.889235916 |
| EPC1 | 1.035735634 | 1.307311148 |
| TAF1 | 1.031281885 | 1.441544389 |
| FYTTD1 | 1.027850306 | 2.561354781 |
| PDS5B | 1.023328528 | 1.851668222 |
| RCOR1 | 1.022359049 | 1.474877939 |
| GTF3C3 | 1.017466479 | 1.991940571 |
| THOC2 | 1.007783634 | 1.44284849 |
| HOXA13 | 1.006641695 | 1.548647609 |
| TSHZ3 | 1.005807615 | 1.50040035 |
| L3MBTL2 | 0.999371559 | 1.302363927 |
| GTF2H4 | 0.997883724 | 1.722474331 |
| CDKN2AIPNL | 0.994829294 | 1.584337086 |
| ALYREF | 0.99334234 | 1.361170206 |
| HDAC2 | 0.991864997 | 1.476517004 |
| BICRAL | 0.9841158 | 1.911859519 |
| SMC6 | 0.983908423 | 1.468365806 |
| CREBBP | 0.981285015 | 1.878406047 |
| WDR36 | 0.971175055 | 1.454512217 |
| SNRPB2 | 0.969461647 | 1.561782464 |
| MGA | 0.964830445 | 1.680337384 |
| CWF19L2 | 0.961859386 | 1.665377507 |
| GTF2H2;GTF2I | 0.959690156 | 1.799975048 |
| POLDIP3 | 0.954126953 | 1.399797803 |
| RNF2 | 0.949429389 | 1.411629284 |
| GTF2H3 | 0.942118673 | 1.458597698 |
| INO80 | 0.941304842 | 1.591365946 |
| ASH2L | 0.936038253 | 2.016656272 |
| GEMIN4 | 0.933850834 | 2.763277575 |
| SMARCD2 | 0.93308144 | 1.363063504 |
| PCGF2 | 0.92252093 | 1.830103838 |
| SLU7 | 0.922376877 | 2.034803024 |
| GTF3C1 | 0.915764121 | 1.319402577 |
| THAP4 | 0.914081378 | 1.396244131 |
| INTS10 | 0.905718949 | 1.509938694 |
| BPTF | 0.90345997 | 1.795675059 |
| CIC | 0.903223035 | 1.472144635 |
| RBM28 | 0.89188162 | 2.819430786 |
| DDX56 | 0.889100445 | 1.673422842 |
| MLLT10 | 0.883640437 | 1.413887907 |
| ZFP90 | 0.880121562 | 2.254671804 |
| ARID3A | 0.879516328 | 1.531060757 |
| ARID3B | 0.875635475 | 1.309351228 |
| ZBTB7A | 0.87080024 | 1.572422119 |
| API5 | 0.868723995 | 1.544966861 |
| LRRC41 | 0.868564153 | 2.07898562 |
| MIER1 | 0.857952423 | 1.681316216 |
| INTS12 | 0.857600491 | 1.526709919 |
| BBX | 0.857518184 | 1.544823939 |
| DCLRE1A | 0.851779321 | 1.390913037 |
| CRAMP1 | 0.84931463 | 1.395494813 |
| ZBTB39 | 0.847730768 | 1.790836027 |
| HCFC1 | 0.836124653 | 2.04699547 |
| DDX18 | 0.826550362 | 1.519810563 |

|  |  |  |
| --- | --- | --- |
| AHCTF1 | 0.8215696 | 1.507913406 |
| CHD4 | 0.814898214 | 1.57678811 |
| LUC7L2 | 0.814006855 | 2.020902587 |
| TAF1L | 0.81079598 | 1.970526453 |
| REST | 0.806369551 | 1.636712479 |
| MCM9 | 0.798583673 | 1.350597733 |
| BCORL1 | 0.797833629 | 1.544225173 |
| SAGE1 | 0.797438154 | 1.746685407 |
| MED25 | 0.788894097 | 1.340075726 |
| ZEB1 | 0.788529552 | 1.801296443 |
| TCF20 | 0.787283914 | 1.431372987 |
| RFC4 | 0.77910037 | 1.940259875 |
| RB1 | 0.769037185 | 1.720462041 |
| ZFP62 | 0.767701207 | 2.335822847 |
| MYEF2 | 0.765650812 | 1.511681405 |
| KDM4B | 0.762734132 | 1.63538578 |
| WDHD1 | 0.759404154 | 1.739495842 |
| ING3 | 0.757984664 | 1.349738706 |
| PKNOX1 | 0.756538708 | 1.520001002 |
| CEBPZ | 0.756445026 | 1.55250281 |
| BHLHB9 | 0.75484247 | 1.553976586 |
| DKC1 | 0.753009664 | 1.777756971 |
| EP400 | 0.752635029 | 1.790304584 |
| CDKN2A | 0.752628684 | 1.49623888 |
| PHC2 | 0.750550761 | 1.446834661 |
| ZKSCAN2 | 0.746996444 | 1.562924011 |
| TAF1A | 0.744284459 | 1.502957567 |
| METTL16 | 0.742076394 | 1.308256874 |
| PHF12 | 0.740021093 | 1.468958966 |
| ATAD5 | 0.731203509 | 1.485719916 |
| JADE3 | 0.730002075 | 1.444521552 |
| BAG2 | 0.724594508 | 1.316922802 |
| ZNF512 | 0.715642805 | 1.375647574 |
| ZMYM3 | 0.714920364 | 1.418753935 |
| QSER1 | 0.712709445 | 1.434263288 |
| KMT2E | 0.710250332 | 1.427784544 |
| CCDC82 | 0.708854003 | 1.496372759 |
| SFMBT2 | 0.707516527 | 1.976279664 |
| KDM3B | 0.70207072 | 1.439769807 |
| ZBTB3 | 0.698194907 | 1.702110751 |
| RADX | 0.697842522 | 1.723033131 |
| C7orf50 | 0.695001846 | 1.318144834 |
| HMG20B | 0.691015208 | 1.30288358 |
| SHPRH | 0.682920823 | 1.840799723 |
| PRPF31 | 0.681402318 | 1.451731032 |
| POLR2L | 0.679178947 | 1.782591268 |
| EXO1 | 0.671330067 | 1.405909828 |
| FBXL6 | 0.665084998 | 2.778419229 |

|  |  |  |
| --- | --- | --- |
| METTL3 | 0.655184835 | 1.703883798 |
| DDX47 | 0.652625976 | 1.302548559 |
| CHAF1B | 0.651684123 | 1.818620005 |
| DDX27 | 0.649570115 | 1.680910005 |
| MTF2 | 0.638126497 | 1.742001423 |
| SF3A3 | 0.634315705 | 1.927089183 |
| CPSF3 | 0.63316747 | 1.319739061 |
| NFIX | 0.631967077 | 1.639476931 |
| EXOSC9 | 0.63176263 | 1.89199558 |
| REXO4 | 0.629416935 | 1.844261965 |
| HMX1 | 0.628134688 | 1.509677266 |
| NKX2-1 | 0.624523803 | 1.637325934 |
| FOXF2 | 0.62395551 | 1.416324136 |
| DDX11L8 | 0.61766836 | 1.357320068 |
| MSH3 | 0.615749874 | 1.375820636 |
| SDE2 | 0.611412332 | 1.531550716 |
| UTP18 | 0.610967643 | 2.334166806 |
| DDX5 | 0.605753849 | 1.444861155 |
| INCENP | 0.601968048 | 1.427758434 |
| ZNF503 | 0.601227344 | 1.536978646 |
| C9orf78 | 0.599824907 | 1.423477014 |
| EZH2 | 0.597186754 | 2.071558597 |
| ACD | 0.595923649 | 1.449188868 |
| WDR46 | 0.592037067 | 1.769125769 |
| DZIP3 | 0.57828085 | 2.00302717 |
| RLF | 0.57823856 | 1.855829258 |
| FAM76B | 0.578084091 | 1.356373948 |
| PRR3 | 0.571856659 | 1.688724072 |
| RFWD3 | 0.567646231 | 3.121671204 |
| WDR75 | 0.566420224 | 1.408453595 |
| ZBTB8A | 0.565952242 | 1.889001918 |
| PHF6 | 0.56149958 | 2.353507757 |
| RBM19 | 0.536412043 | 1.430767298 |
| SARNP | 0.535909953 | 1.608622951 |
| TAF7 | 0.535588231 | 1.823112904 |
| NFIB | 0.534808012 | 1.537507619 |
| NOL11 | 0.528802718 | 1.699072402 |
| GATAD2B | 0.52533188 | 1.495778829 |
| KDM5C | 0.525284154 | 1.313568353 |
| ZFP37 | 0.519677429 | 1.387126622 |
| TFDP1 | 0.515409734 | 1.666852508 |
| ECT2 | 0.509897063 | 1.958129265 |
| YEATS4 | 0.509073941 | 1.933024917 |
| ZNF22 | 0.508660122 | 1.821016262 |
| ENY2 | 0.507278245 | 2.518515205 |

Log2FoldChange was calculated by two independent experimental replicates. Ribosomal proteins and

s)
