## Supplementary Table 6 for "RNA m6A mediates *de novo* DNA methylation and heritable epigenetic silencing in human cells"

**Table S6. List of sgRNA sequences**

| **Gene** | **Protospacer (5' to 3')** |
| --- | --- |
| CD55 | CTGCGACTCGGCGGAGTCC |

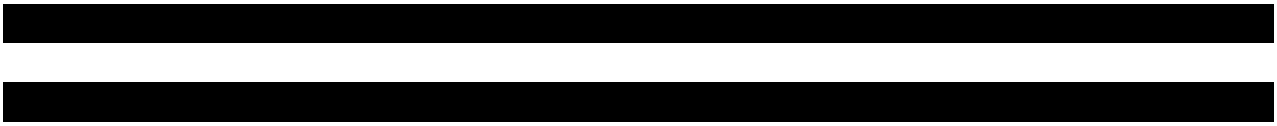

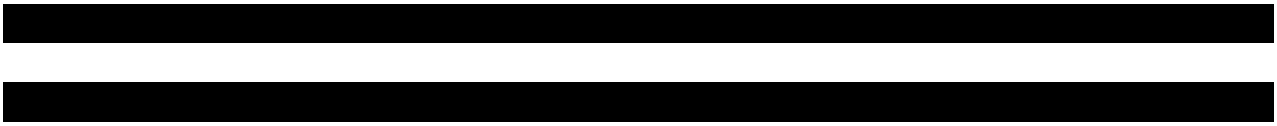

[REDACTED]

[REDACTED]
