## Supplementary Table 7 for "RNA m6A mediates *de novo* DNA methylation and heritable epigenetic silencing in human cells"

**Table S7. sgRNA and donor sequences for generation of tagged HEK293T cell lines for immunoprecipita**

| Gene | sgRNA | ssODN |
| --- | --- | --- |
| SEDTB1 | AAAAGCATGTCTTCCCTTCC | GTTATAAGGACTCCAATTTAATTTGTTTTCTGTTCCATGC<br>AGAGGACAAAAGCATGGGTGGTGGTGGTTCTGGTGGT<br>GGTGGTTCTACCGAGCTCAACTTCAAGGAGTGGCAAAA<br>GGCCTTTACCGATATGATGagcagttTgCCTGGGTGCATTG<br>GTTTGGATGCAGCAACAGCTACAGTGGAGTCTGAAGAG<br>ATTGC |
| DNMT3A | AGTATTTTGCGTGTGTGTAA | CATGGAGCGTGCCAGTCATCCGCCACCTCTTCGCTCCGCT<br>GAAGGAGTATTTTGCGTGTGTGGGTGGTGGTGGTTCT<br>GGTGGTGGTGGTTCTACCGAGCTCAACTTCAAGGAGTG<br>GCAAAAGGCCTTTACCGATATGATGTAAGGGACATGGG<br>GGCAAACTGAGGTAGCGACACAAAGTTAAACAAACAAA<br>CAAAAAACA |

tion experiments

[REDACTED]

[REDACTED]

[REDACTED]

[REDACTED]

\_\_\_\_\_

\_\_\_\_\_

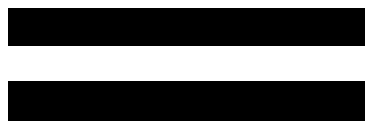
