## Supplementary Table 8 for "RNA m6A mediates *de novo* DNA methylation and heritable epigenetic silencing in human cells"

**Table S8. List of antibodies used in this study**

| **Antibodies**<br>**(target)** |  | **Company** | **Catalog #** | **Experiment** | **Dilution** |
| --- | --- | --- | --- | --- | --- |
| CD55 | APC anti-human<br>CD55 Antibody | BioLegend | 311312 | Flow<br>cytometry |  |
| CD81 | CD81 (TAPA-1)<br>Antibody | BioLegend | 349510 | Flow<br>cytometry |  |
| SETDB1 | Rabbit pAb<br>SETDB1 antibody | Proteintech | BC009362 | Western<br>Blot | 1:1000<br>dilution |
| DNMT3A | Rabbit pAb<br>DNMT3A antibody | Proteintech | 20954-1-AP | Western<br>Blot | 1:2000<br>dilution |
| VIRMA | Rabbit pAb<br>VIRMA antibody | Proteintech | 25712-1-AP | Western<br>Blot | 1:2000<br>dilution |
| WTAP | Mouse mAb<br>WTAP antibody | Proteintech | 60188-1-Ig | Western<br>Blot | 1:1000<br>dilution |
| METTL3 | Rabbit METTL3<br>antibody | Proteintech | 15073-1-AP | Western<br>Blot | 1:2500<br>dilution |
| YTHDC1 | Rabbit YTHDC1<br>antibody | Proteintech | 29441-1-AP | Western<br>Blot | 1:1000<br>dilution |
| YTHDF2 | Rabbit YTHDF2<br>antibody | Proteintech | 24744-1-AP | Western<br>Blot | 1:2500<br>dilution |
| GAPDH | Mouse GAPDH<br>antibody | abcam | Ab245355 | Western<br>Blot | 1:1000<br>dilution |
| B-Actin | Mouse mAb B-actin antibody | Cell Signaling | 3700S | Western<br>Blot | 1:2000<br>dilution |
| H3K9me3 | Histone H3K9me3<br>antibody (pAb) | Active Motif | 39766 | ChIP-qPCR |  |

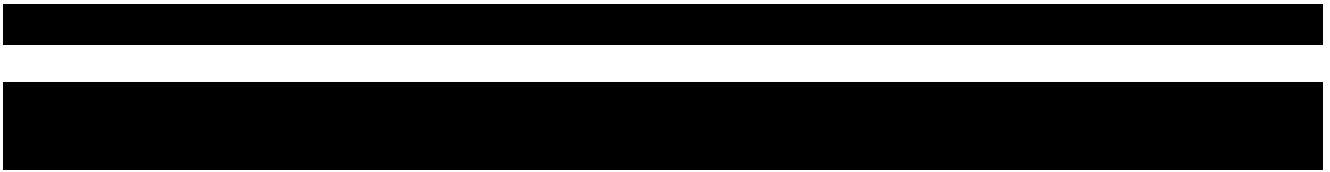

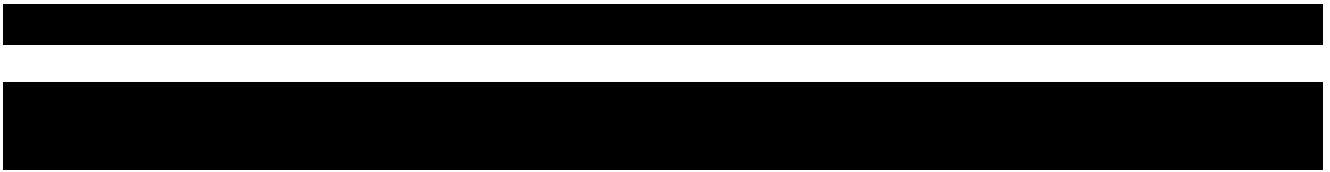

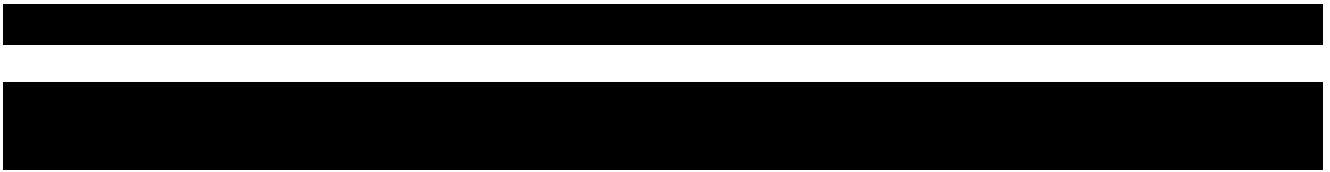

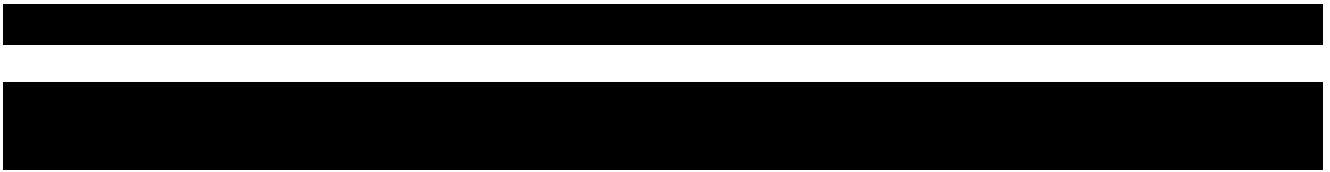

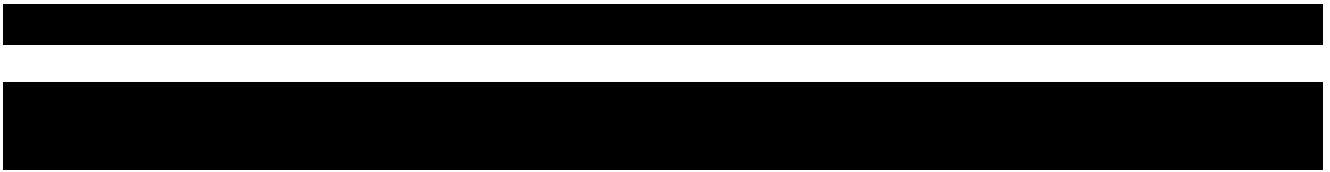

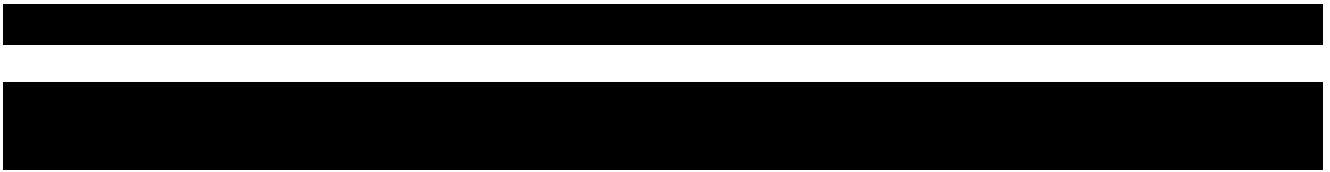

[REDACTED]

[REDACTED]
