## Supplementary Table 9 for "RNA m6A mediates *de novo* DNA methylation and heritable epigenetic silencing in human cells"

**Table S9. DNA methylation profiling primers**

| Target Name | Oligo 1 sequence | Oligo 2 sequence | Amplicon Size (kb) |  |
| --- | --- | --- | --- | --- |
| CD55 | TGGGTTGATGGGTGT<br>AGTAAATTAT | ACACAACAAAAACCCCA<br>AATACTAA | 1.43 |  |
| CLTA | TATTTGTTGATTGGGT<br>AGTTTTTGA | AACATCACACACAAATT<br>AAAACATC | 1.215 |  |
| L1Hs 5' UTR | GGGTGATTTTTGTATT<br>TTTATTTGAG | CAAAAACAAACAAACCT<br>CCTTAAAC | 0.416 | from RepBase<br>L1HS consensus |
| LTR5H | GTGTTGTGTTAATTTA<br>GGGTAAATG | CCCTACACACCTATAAA<br>TATTTCTC | 0.744 | from RepBase<br>LTR5_H<br>consensus |
| HERVH (primer set A) | TGGGAGATTAATTTTT<br>TGTTTTTTT | ATATCAAACACCTCAAA<br>CCATTTAC | 1.067 | from RepBase<br>HERVH<br>consensus |
| HERVH (primer set B) | GGGGATTTTTTTTGG<br>GAGAT | ATCAAAAAACCTTAAAC<br>CAAATTC | 2.006 | from dfam<br>consensus<br>DF000000183.4 |

[REDACTED]

[REDACTED]
