## Supplementary Table 10 for "RNA m6A mediates *de novo* DNA methylation and heritable epigenetic silencing in human cells"

**Table S10. Sequences of Quantitative PCR primers used in this study**

| Target | Forward primer<br>sequence | Reverse primer<br>sequence |  |
| --- | --- | --- | --- |
| HERV-K pol | TCACATGGAAACAG<br>GCAAAA | AGGTACATGCGT<br>GACATCCA | RT-qPCR |
| L1Hs-ORF1 | GACATCTACACCGA<br>AAACCC | TCGTCAAAATCAT<br>TCTCCATCC | RT-qPCR |
| ATF4 | CTTGATGTCCCCCT<br>TCGACC | CTTGTCGCTGGA<br>GAACCCAT | RT-qPCR |
| GAPDH | GAAGGTGAAGGTC<br>GGAGTCAAC | CAGAGTTAAAAG<br>CAGCCCTGGT | RT-qPCR |
| ACT-B | CGACAGGATGCAG<br>AAGGAG | ACATCTGCTGGA<br>AGGTGGA | RT-qPCR |
| CD55 | CACGAGGCTTCTG<br>CTTACTG | GCGCCGGGTTA<br>GAACAA | ChIP-qPCR |
| GAPDH | CACCGTCAAGGCT<br>GAGAACG | ATACCCAAGGGA<br>GCCACACC | ChIP-qPCR |

[REDACTED]

[REDACTED]
